# Seed dormancy increased population persistence in permissive environments, but not in stressful environments in an annual plant

**DOI:** 10.1101/2025.08.22.671647

**Authors:** Brandie Quarles-Chidyagwai, Kathleen Donohue

## Abstract

**Background and Aims:** Seed dormancy can delay germination timing to more favorable growth conditions, not only increasing seedling survival, but potentially increasing lifetime fitness. As such, seed dormancy can be a form of seasonal environmental tracking. In addition, seed dormancy can act as a bet-hedging strategy by spreading the germination risk across time, within or between years. Through both environmental tracking and bet-hedging, seed dormancy can stabilize population demography, potentially enhancing long-term population persistence.

**Methods:** To test whether populations that express seed dormancy are more likely to persist than populations not capable of dormancy, we established genetically variable, experimental field populations of *Arabidopsis thaliana* that differ in their capacity to control the seasonal timing of germination through seed dormancy. Four environmental treatments were imposed to test for demographic differences across environments and to test whether dormancy mitigates the effects of environmental variation.

**Key Results:** Seasonal seed dormancy influenced demography and population persistence primarily via early seedling or rosette mortality. Dormant populations had larger seedling populations and higher population persistence over the three years in the most permissive environmental treatments. However, stressful environments diminished the demographic effects of dormancy. These dynamics, in turn, resulted in dormant populations unexpectedly exhibiting more variation across environmental treatments than non-dormant populations. Therefore, dormancy’s enhancement of demographic performance may be caused more by allowing populations to take advantage of favorable conditions than by helping them to escape poor conditions.

**Conclusions:** This study shows that seasonal seed dormancy may help populations persist over time, but not under all environmental conditions. In more permissive environments, dormancy can reduce population bottlenecks and maintain larger populations. Some conditions, however, may be too adverse for seed dormancy to overcome.

## Introduction

Seed dormancy is the process through which plants can control their germination timing and delay growth until more permissive conditions occur (Finch-Savage and Leubner-Metzger 2006). Seasonal dormancy precisely regulates seasonal timing of germination within a year, while between-year dormancy delays germination across years, resulting in a seed bank. Both types of dormancy are forms of developmental arrest, like diapause, a process that has been documented across a wide span of organisms to allow them to better track and survive variable environmental conditions (Baskin and Baskin 2004; Furness *et al*. 2015; Sgrò *et al*. 2016; Penfield 2017). Accordingly, dormancy is predicted to have significant demographic consequences. Specifically, as a form of bet-hedging, seasonal and between-year dormancy can buffer populations against large between-year fluctuations in population size (Evans and Dennehy 2005; Živković and Tellier 2012; Miller *et al*. 2025). In addition, seasonal dormancy can act as a form of phenological environmental tracking where plants use environmental cues to regulate seasonal timing of germination within a year (Donohue *et al*. 2010; D’Aguillo *et al*. 2019). Through phenological tracking, seasonal dormancy can increase lifetime fitness at the individual level and reduce extinction risk at the population-level (Donohue *et al*. 2005a; Willis *et al*. 2014). As the effects of climate change become more prevalent, it is critical to investigate the mechanisms or traits that may facilitate population persistence in the face of variable and changing seasonal environments.

Bet-hedging is defined as the spread of demographic risk, i.e. reducing the variance in fitness at the cost of reducing mean fitness (Seger and Brockman 1987; Philippi and Seger 1989). Seed dormancy acts as a bet-hedging strategy by spreading the demographic risk of germination across time (Evans and Dennehy 2005; Venable 2007; Gremer and Venable 2014; Torres-Martínez *et al*. 2017). Through bet-hedging, dormancy can stabilize population demography by buffering against environmental variation, thereby increasing the effective population size (Ne) (Lundemo *et al*. 2009; Falahati-Anbaran *et al*. 2011; Gianella *et al*. 2021). Most studies of dormancy as a bet-hedging strategy have focused on between-year dormancy, fewer have focused on the effects of variable germination timing within a year, i.e. seasonal dormancy (except see Donohue et al., 2005b; Gremer et al., 2016).

Phenological tracking, when species use environmental cues to shift the timing of developmental events to track seasonal environments that are more favorable for survival and reproduction, is a form of habitat selection (Wolkovich and Donahue 2021). Many studies have focused on reproductive life stages, documenting the effects of shifts in flowering time (Bertin 2008; Tooke and Battey 2010; Cleland *et al*. 2012; Jagadish *et al*. 2016; Tun *et al*. 2021; Iler *et al*. 2021). However, theoretical (Burghardt *et al*. 2016) and empirical (D’Aguillo *et al*. 2019; D’Aguillo and Donohue 2023) work has also found that cued seasonal dormancy is a form of habitat selection as it significantly influences the seasonal environmental conditions plants experience post-germination (reviewed in Donohue, 2003). Therefore, seasonal seed dormancy can not only influence survival immediately after germination, but also survival to later stages and reproductive success by determining the seasonal environmental conditions that a plant will experience throughout its life cycle.

Through phenological tracking, seasonal seed dormancy can reduce extinction risk and potentially increase population size. Cued seed dormancy has been shown to increase survival and reproduction in some species (González-Astorga and Núñez-Farfán 2000; Donohue *et al*. 2005a; Picó 2012). Increased individual fitness could maintain population sizes over time, or even result in population growth. Furthermore, higher fitness, as a result of efficient tracking of seasonal environmental conditions can reduce population extinction risk (Willis *et al*. 2014).

Since more frequent stressful environments are expected with climate change (Chaudhry and Sidhu 2022), it is important to investigate the effects of dormancy under various environmental conditions. Little is known about how widespread dormancy’s effects on population demography can be across diverse environments. Through habitat selection, seasonal dormancy may mitigate the effects of environmental variation on population demography by ensuring that seedlings experience only a subset of the variable environments that are available for them to germinate into (D’Aguillo *et al*. 2019). Therefore, if dormancy can act as a form of phenological tracking, more consistent performance across environments that vary over time may be expected. Alternatively, if an environment is chronically stressful it may repress demographic performance regardless of seed dormancy. More research is needed to test whether dormancy can provide a demographic advantage in seasonally and chronically stressful conditions. The demographic benefits of dormancy may result from its ability to allow plants to take advantage of favorable conditions, or by allowing plants to avoid stressful conditions. If the former, we would expect that dormancy would have a larger effect on population demography in more permissive environments whereas, if the latter, we would expect dormancy to have a greater effect in more stressful environments.

The environment can also have a direct effect on the amount of seasonal dormancy that is expressed (Lu *et al*. 2016). A field experiment investigating differences in germination across geographical and seasonal variation found differences in the frequency of non-dormant phenotypes expressed across environments, suggesting that dormancy induction and maintenance is environmentally dependent (Donohue *et al*. 2005b). If the amount of seasonal dormancy, or environmental tracking, that is expressed is dependent on the environment, the demographic benefits of dormancy would not be expected in all environmental scenarios.

In this study, we established genetically variable, experimental field populations of *Arabidopsis thaliana* (Brassicaceae) that differed in alleles at 2-3 major dormancy loci to evaluate the demographic effects of seed dormancy. Specifically, we tested 1) whether seed dormancy would increase the size of the seedling class, stage-specific survival, and reproductive output, 2) whether seed dormancy would reduce fluctuations in total seedling number across years and the occurrence of local population extinction, and 3) whether the demographic effects of seed dormancy would be repressed in stressful environments. Since seed dormancy can be expressed as either seasonal or between-year dormancy, we estimated differences in both forms of dormancy between populations and performed population projection models to test whether seasonal dormancy and/or between-year dormancy would mitigate the effects of environmental variation on projected population growth.

## Materials and Methods

### Study Species

*Arabidopsis thaliana* is an annual plant in the Brassicaceae family. It exhibits natural variation in seed dormancy and has a plethora of available genetic resources and information that can be used to manipulate dormancy (Bentsink *et al*. 2010; Kronholm *et al*. 2012; Montesinos-Navarro *et al*. 2012; Debieu *et al*. 2013; Postma *et al*. 2016). *A. thaliana* typically has a winter annual life history, germinating in the autumn and flowering in the spring (Baskin and Baskin 1983). However, variation in germination timing can result in a spring-annual life history – germinating and flowering in the spring – or a rapid-cycling life history – having multiple generations per year (Thompson 1994). In the native European range of *A. thaliana,* seasonal seed dormancy tends to be higher in the South compared to the North (Debieu *et al*. 2013; Martínez-Berdeja *et al*. 2020; Zacchello *et al*. 2022). There is also genetic variation for seasonal dormancy within regions that covaries with seasonal climatic differences (Montesinos-Navarro *et al*. 2012; Zacchello *et al*. 2022). *A. thaliana* was introduced to the United States and there is a naturalized population in Durham, NC near where the experiment was set up. In this naturalized population, *A. thaliana* exhibits a winter annual life cycle with peak germination in mid-October (Mauricio and Rausher 1997; Leverett 2017).

### Genetic Composition of the Experimental Populations

To test whether seasonal seed dormancy alters population dynamics, we used recombinant inbred lines (RILs) to establish genetically variable populations that differ in seed dormancy. Previous studies identified a small number of large-effect QTLs for seed dormancy in several RIL sets in *A. thaliana* (Alonso-Blanco *et al*. 2003; Laserna *et al*. 2008; Meng *et al*. 2008; Bentsink *et al*. 2010; Huang *et al*. 2010; Postma and Ågren 2015). We selected three RIL sets to use for this study: Castelnuovo-12 x Rodasen-47 (“Italy x Sweden” hereafter), Bay-0 x Shahdara (Bayreuth, Germany x Shakhdara, Tajikistan; “Germany x Tajikistan” hereafter), and Cal-0 x Tac-1 (Calver, England, UK x Tacoma, Washington, USA; “UK x USA” hereafter). Seeds for each RIL set were ordered from the Arabidopsis Biological Resource Center (see “ABRC Stock #” in Supplementary Table 1). Each RIL set had 2-3 dormancy QTLs of large effect, identified by prior QTL studies (Table S1). The RIL sets serve as independent assessments of dormancy manipulation as they differ in their genetic backgrounds and the dormancy loci with segregating alleles.

Lines that were homozygous for alleles associated with high dormancy were selected to make up “Dormant” populations, and lines that were homozygous for alleles associated with non-dormancy were selected to make up “Non-Dormant” populations. To maintain genetic variation at other loci, lines were selected to ensure that allele frequencies across the rest of the genome were as close to 50% as possible. The number of lines that met the criteria for being in the Dormant and Non-Dormant population varied by RIL set (see “Homozygous lines” in Table S2). In addition, to provide segregating genetic variation, we conducted crosses within the Dormant and Non-Dormant populations, separately, to permit effective recombination within the experimental populations (See “Heterozygous lines” in Table S2). The number of heterozygous lines varied by RIL set based on the success of the attempted crosses. Each population pool in Italy x Sweden and Germany x Tajikistan contained approximately 15% heterozygosity, but the UK x USA lines did not yield any heterozygotes. To allow for the evolution of dormancy itself during the experiment, “Mixed” populations were composed of equal numbers of each “dormant” and “non-dormant” line, providing a population with 50% dormant and 50% non-dormant individuals. While there is some degree of linkage disequilibrium around the loci fixed for dormancy alleles, allele frequencies throughout the rest of the genome were close to 50% in all RIL sets, based on marker allele frequencies from prior studies, which were provided with the RIL sets (Supplementary Fig. 1).

### Seed Bulking Conditions and Characterization of Seed Dormancy in the Lab

To standardize the maternal environment, the selected lines were first grown in the Duke University Phytotron. Three replicates of each individual line within each RIL set were grown in growth chambers (EGC Model GCW30; Chagrin Fall, OH, USA). More specifically, plants were stratified in the dark at 4°C [5-6 days for Italy x Sweden and 4 days for Germany x Tajikistan and UK x USA], grown at 22°C with 16-hour days until the first 2 true leaves developed, vernalized at 4°C with 12-hour days (8 weeks for Italy x Sweden, 3 weeks for Germany x Tajikistan and UK x USA), and then grown at 22°C with 16-hour days until maturity. Fresh seeds were harvested once the plants had completely dried out and matured all their fruits. Immediately after the establishment of the field experiment (see below), 26-36 days after seed harvest, germination assays were conducted in the lab to ensure that the experimental populations differed in seed dormancy (Italy x Sweden 2-4 days, Germany x Tajikistan and UK x USA 0-2 days after field establishment; for more details see Supplementary Information).

### Field Experimental Design

144 replicate population cages were established in a fallow old-field site on Duke Forest property, Durham, NC (36.009169, −79.018746). For all 3 RIL sets, three dormancy treatments (Dormant, Non-dormant, and Mixed) were established with 8 replicate populations in a “Control” environmental treatment (3 x 3 x 8 = 72 experimental populations; Fig. 1). Three additional environmental treatments (see below) were established for Italy x Sweden, with 8 replicates per dormancy treatment (3 x 3 x 8 = 72 experimental populations). The field was divided into 8 spatial blocks with one replicate of each RIL set, dormancy, and environmental treatment combination per block.

**Figure 1.**
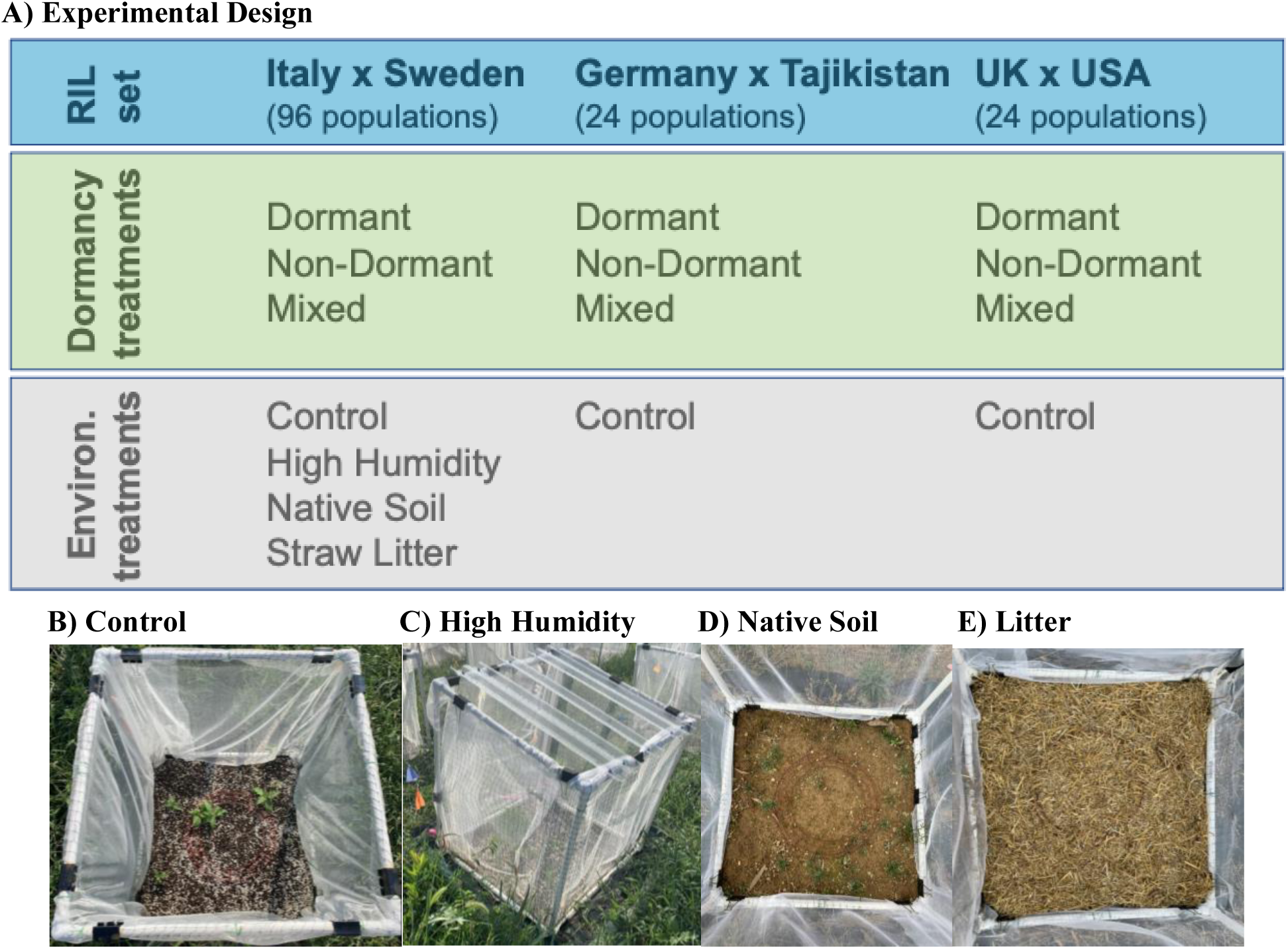
Field Experimental Design. There were 3 RIL sets, 3 Dormancy treatments, and 4 Environmental treatments. There were 8 replicates per RIL set x Dormancy treatment x Environmental treatment combination. The total number of populations per RIL set is listed. The Control treatment B) had greenhouse soil, the High Humidity treatment C) had roof panels on top of the cages and trenches around the bottom of the cages, the Native Soil treatment D) had steam pasteurized North Carolina clay soil, and the Litter treatment E) had sterilized straw laid on top of the greenhouse soil.

Holes (61cm x 61cm x ∼10.2cm deep) were dug in the field and filled with greenhouse soil (Fafard #2) for Control, Litter, and High Humidity treatments or steam pasteurized North Carolina clay soil (taken from the field site) for the Native Soil treatment. Cages were 61cm high, placed 61-91cm apart, and constructed out of PVC pipes, deer netting, and white organza fabric to prevent seed contamination across experimental populations; the tops of cages were left uncovered. The fabric was replaced every year before the natural seed dispersal season to ensure maximum protection against seed contamination. Since *A. thaliana* is a poor competitor that frequently inhabits disturbed habitats, we staked strips of shade cloth, 6-inches wide, around the outside of cages and regularly weeded inside and outside cages to eliminate the effects of inter-specific competition.

Phytotron-produced seeds, described above, were dispersed into the field at two different time points, summer, and fall. Italy x Sweden populations’ summer dispersal occurred on May 20, 2019 (seeds 26-32 days after-ripened) during the natural seed dispersal season for *A. thaliana*, and the summer dispersal for Germany x Tajikistan and UK x USA occurred on July 9, 2019 (seeds 26-29 days after-ripened). After observing low germination in the fall of 2019, we conducted another round of dispersal for all RIL sets on November 17, 2019 (seeds 7 months after-ripened for Italy x Sweden and 5 months after-ripened for Germany x Tajikistan and UK x USA). The exact numbers of seeds dispersed into each population are given in Table S2 and ranged from 616 to 640. Populations were then allowed to naturally cycle through the end of 2021.

### Environmental Manipulations

Four environmental treatments were established to test how the effects of dormancy vary across environments and how dormancy alters the demographic effects of environmental variation (Fig. 1). “Control” populations used greenhouse soil with no additional manipulation. To manipulate soil moisture, roof panels were installed on top of cages and trenches were dug around the bottom of cages and filled with plastic to prevent lateral movement of water. We call this treatment “High Humidity” because it led to higher soil moisture and temperature, as described below. “Native Soil” populations had native soil from the field site, with no additional manipulation, to test for edaphic environmental effects. The native soil was steam pasteurized to kill any pre-existing seeds (see Taylor et al., 2017). For a “Litter” treatment, sterilized straw was laid on top of the soil during the first winter season and left there throughout the experiment. Litter accumulation has been shown to create stressful conditions, especially during seed recruitment (Jessen *et al*. 2023). Greenhouse soil was used for the Control treatment for practical reasons (it was not possible to transport and sterilize soil from every population plot, given the resources) and to impose a stark contrast between amended and unamended soil, since *A. thaliana* frequently colonizes agricultural sites with intense soil amendment.

To test for environmental differences among the treatments, soil temperature (DS1921G Thermochron iButton; Maxim Integrated Products, Inc) and soil water content (SWC, i.e. volumetric water content; HOBO Micro Station Data Logger, EC5 and 10HS Soil Moisture Smart Sensors; Onset Computer Corporation, Bourne, MA, USA) were recorded every four hours for all environmental treatments throughout the experiment. One replicate per environmental treatment (regardless of dormancy treatment) was measured in each of 4 blocks for soil temperature and 6 blocks for SWC. From the environmental sensor data, we calculated a daily average, maximum, and minimum soil temperature, and moisture. Negative SWC values resulted from air pockets developing in very dry soil, so all negative SWC values were converted to zero before calculating the daily average, max, and min (Campbell 2019). In addition, to ensure that the roof panels on the High Humidity treatment did not significantly influence light availability, we also measured the photosynthetic photon flux density (PPFD; MQ-200 Quantum Meter; Apogee Instruments Inc, Logan, Utah, USA) in one Control and one High Humidity population in all 8 blocks. These measurements were conducted one time in September 2021 at solar noon. Four samples were taken in each of the measured population cages to account for spatial variation within a population cage.

To assess how the environmental treatments differed from each other across the three years, an ANOVA with environmental treatment, month-year, and interactions between those effects as fixed factors and block as a random factor was used for soil temperature and moisture (PROC MIXED, SAS 9.4 SAS Institute, Inc. 2015). Tukey’s tests were used to test for significant pairwise differences between treatments within each month-year. An ANOVA with just environmental treatment (Control versus High Humidity) as a fixed effect and block as a random effect was used for PPFD. To improve normality, PPFD was natural log transformed.

### Estimating Population-level Demographic Parameters

To monitor population size and age structure over time, we collected population-level data for all 144 population cages for three years. Circles with a diameter of 24.77-cm were placed in the center of population cages to standardize the area of the census, since for some life stages the area of the whole population was too large to census on a regular basis. Population censuses were conducted weekly during the summer (late May – early September), at least every two weeks during the fall germination period (usually late September – October), at least monthly during the winter rosette growth period (November – January), and at least monthly during the spring reproductive period (February – early May). After the third year’s peak germination period in autumn, most of the plants that germinated were transplanted and taken to the phytotron for leaf tissue and seed collection for future studies. Therefore, the year 3 census data ends in December 2021. This represents 3-5 generations, since *A. thaliana* can sometimes exhibit more than one generation per year.

During each population census, the numbers of seedlings, rosettes, and adults that were bolting, flowering, fruiting, or mature/dead inside the circle were recorded. A seedling was classified as a plant with only cotyledons or with 2 cotyledons and 2 developing true leaves; a rosette was a plant with fully developed true leaves; a bolting plant was one which had recently transitioned to reproduction but which had not yet flowered; a flowering plant was a plant with at least one open flower; a fruiting plant was a plant with at least one fruit (including mature and not yet mature fruits); and a mature/dead plant was one with completely mature fruits/seeds. In addition, in years 1 and 2, the census also recorded the number of reproductive individuals outside the circle for an estimate of total reproductive population size of the entire population. From these population census data, we were able to calculate estimates of the size of each developmental stage/class over time, transitions between life stages, and population persistence. All data manipulation and calculations of derived variables were conducted using the tidyverse package (Wickham *et al*. 2019) in R v. 4.0.1 (R Core Team 2020).

Size of the seedling class was estimated as the total number of seedlings that emerged during a given year. The estimate of total seedling number assumes that all seedlings survived between censuses, either remaining as seedlings or transitioning to the rosette stage. In addition, since the censuses occurred every 1-2 weeks, it is possible that a new rosette could be a fast-growing new germinant that germinated in between censuses. Thus, the number of new germinants at each census was estimated as the total number of seedlings plus the number of new rosettes, minus the number of seedlings from the previous census. The estimate of the size of the seedling class was calculated as the sum of new germinants over all censuses within a year.

The size of the rosette class was calculated by taking the sum of the number of new rosettes (Rosettes_(t)_ – Rosettes_(t-1)_) inside the circle at each census. Note that some rosettes could be counted as both new germinants in the seedling class estimates and new rosettes in the rosette class estimates. The total number of reproductive individuals was calculated by taking the sum of the fruiting and mature/dead each week (both inside and outside the circle, separately) and adding up all the new reproductive individuals ((Fruiting_(t)_ + Mature/Dead_(t)_) – (Fruiting_(t-1)_ + Mature/Dead_(t-1)_) across censuses. When calculating the number of new individuals for each adult life stage, if the difference between censuses was negative, it was converted to 0. Fluctuations in population size over time were calculated by taking the difference in the number of plants between consecutive years (year 1 vs. 2 or year 2 vs. 3) at each life stage. Due to the transplanting after peak germination in year 3, we only have complete data for size of the seedling class in year 3.

Since dormancy is expected to have the largest effect on early seedling survival (i.e., seedling establishment), we estimated 2-week seedling survival probabilities for each population. “Seedling establishment” represents the proportion of the total number of germinants that survived for two weeks, either by transitioning to rosette or remaining a seedling (for more details see Supplementary Information).

To assess how dormancy affected survival to later life stages, we also calculated survival from germinant to rosette and survival from rosette to reproduction. Survival from germinant to rosette was estimated as the total number of rosettes divided by the number of seedlings (both inside the circle only). Survival from rosette to reproduction was calculated as the total number of reproductive plants divided by the total number of rosettes (both inside the circle only).

To evaluate population persistence over the three years, we classified the populations based on whether any individuals in the population germinated or reproduced in years 1, 2, and 3 (see Fig. 2 for more details). There were multiple ways for a population to fail, but only one way for them to “persist.”

**Figure 2.**
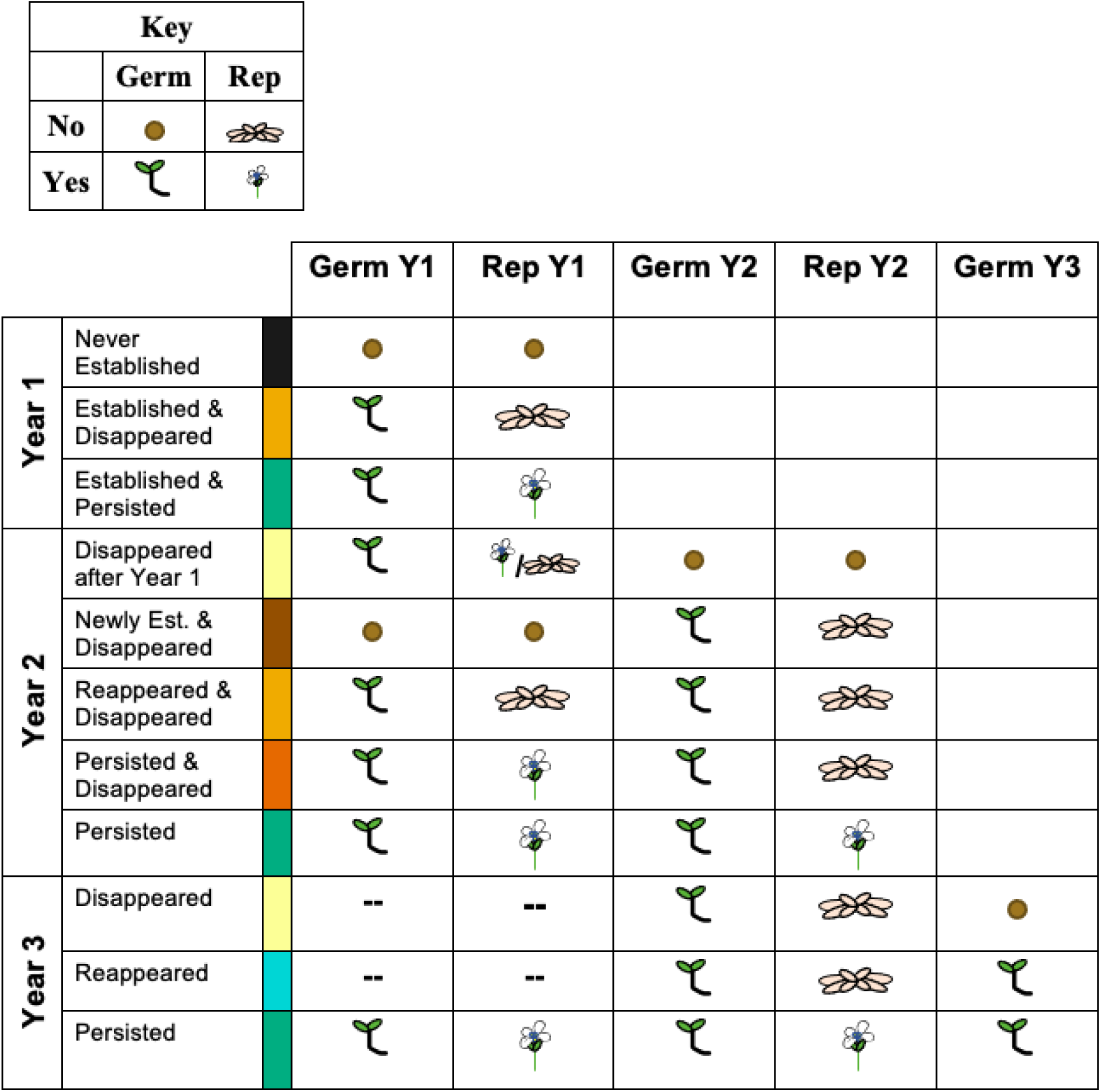
Classifying population persistence. Populations were classified by whether they germinated and reproduced in years 1, 2, and 3. Seeds represent no germination, green seedlings represent successful germination, discolored (pink) rosettes represent no reproduction after successful germination, and flowers represent successful reproduction. All populations that did not have any germination in year 3 had disappeared in year 2, regardless of what happened in year 1. “Reappeared” populations in year 3 include populations that “Disappeared after Year 1” and populations that germinated and disappeared in year 2.

### Statistical Analysis of Population-level Demographic Parameters

For all analysis of the population cage demographic data, census years start after the end of the dispersal season in early summer. There could be multiple generations of *A. thaliana* included in a census year due to rapid cyclers. Note that all generations within a census year contributed to the sizes of each life stage class. Rapid cycling summer and spring generations represent a small proportion of each life stage class.

To test for the effects of dormancy on demographic performance across RIL sets, only the Control treatment was analyzed. To test how the effect of dormancy differed across environmental treatments, only the Italy x Sweden RIL set was analyzed. Sub-models were subsequently used to test the effects of dormancy within each RIL set and environmental treatment. Additional sub-models were used to test the effect of the environment within each dormancy treatment. Tukey’s tests (for ANOVAs), or Hochberg tests (for non-ANOVAS), were used to correct the P-values for the sub-models. All statistical analyses were conducted in SAS (SAS 9.4 SAS Institute Inc. 2012), unless otherwise noted.

We used an ANOVA to test whether dormancy influenced the size of the seedling class, and fluctuations in seedling number between year. PROC MIXED was used, with dormancy treatment, RIL set or environmental treatment, year, any interactions between those effects as fixed effects, and block as a random effect. “Year” is equivalent to “Transition” (year 1 to 2 and year 2 to 3) for fluctuations in the size of the seedling class. To improve normality, we used a log-10 transformation for seedling number (+1) for comparison across RIL sets and a natural log transformation was used for comparisons across environmental treatments.

In addition to the quantitative analysis, we also performed a qualitative analysis of the change in the size of the seedling class between years. The logistic regressions had three response levels: increase, no change, or decrease in seedling number. PROC LOGISTIC was used with a logit-link function Block was also added as a fixed factor in the model.

A logistic regression was also used to analyze the population-level stage-specific survival estimates; a firth option was added to PROC LOGISTIC since the data was binary. The logistic regression analysis was performed in two ways 1) with all the interactions between dormancy, RIL set or environment, and season or year, and 2) with only the main effects because SAS converts the analysis from type III to joint tests when interactions are included in the model. This method correctly models interactions, but not individual main effects. Hochberg tests were not used on the sub models for the seasonal seedling survival data since each year-season was independent.

Since not all populations had survival to each stage, or germination in each season for seasonal seedling establishment, stage-specific survival was not factorial across all eight blocks. Thus, for these data, block was changed to be the two halves of the field with the most difference in water retention (“new block”; observational). New block was included as a fixed effect in all the models of survival from germinant to rosette, but due to remaining factorial issues, it was only included in some of the sub-models for seasonal seedling establishment and survival from rosette to reproduction. For the Italy x Sweden-only analysis, year 1 summer was removed from the seasonal seedling establishment analysis due to quasi-complete separation of data points issues with the model convergence. Year 2 was excluded from the analysis of survival from rosette to reproduction due to low survival to rosette in year 2, and thus low sample sizes.

Population persistence was analyzed in two ways: first with a logistic regression across years with two response levels—persist or not, and second with chi-square contingency tests (PROC FREQ with a chisq option) within each year with all the categories described in Fig. 2 included. The within-year chi-square contingency tests were conducted within each RIL set or environmental treatment, for each pairwise combination of dormancy treatments.

### Individual-level Traits

Due to potential under-replication in the limited circle census area, we augmented the estimates of some vital rates by sampling individuals outside the circle. Up to 25 individual *A. thaliana* plants inside and outside the circle were marked as early in life as possible in Italy x Sweden populations in 5 blocks (total of 60 populations). Plants were marked with toothpicks and small bird bands. Due to high seedling mortality, additional small rosettes were marked to maintain sufficient sample sizes. We recorded survival to reproduction and the total number of siliques per marked individual. Silique number for individuals that survived to reproduce serves as our estimate of per-capita reproductive output. Mean lifetime fitness for each population was estimated as the average silique number, including zeros from individuals that did not survive to reproduction.

An ANOVA testing for the effects of dormancy, environment, and dorm*envt within year 1 was used for per-capita reproductive output; population was included as a random effect. Year 2 was removed from the full-effects model due to low survival to reproduction across all treatments that year. Per-capita reproductive output was natural log transformed to improve normality. Tukey’s tests were used to test pairwise differences between dormancy treatments.

The probability of individual survival to reproduction was analyzed using a logistic regression that tested for the effects of dormancy, environment, year, and all interactions between those effects with each individual in the model represented by 0 (died) or 1 (survived to reproduction). Lifetime fitness could not be normalized, as it was highly zero-inflated. No non-parametric test would be able to account for the three-way interaction between dormancy, environment, and year, so Friedman’s tests were used within years and environmental treatments. Specifically, PROC FREQ was used to test for the effects of dormancy treatment with Cochran-Mantel-Haenszel (cmh2) and rank scores options used to define the Friedman’s test. Block was not included in the analysis of any of the individual-level traits due to a lack of consistent data across blocks. Hochberg tests were used for sub models of survival to reproduction and lifetime fitness.

To verify that silique number was an accurate estimate of per-capita reproductive output, we counted the number of seeds per silique in a subsample of siliques during year 1 (for sampling methods see Supplementary Information). Due to variation in the availability of unbroken siliques, there were 74 dormant and 66 non-dormant siliques sampled. An ANOVA with block as a random factor showed that seed number per silique did not significantly differ between dormancy treatments (Dorm mean: 29.32 +/− 1.23, NonDorm mean: 29.44 +/− 1.00; F=0.00, P=0.9738; PROC MIXED, SAS 9.4 SAS Institute, Inc. 2015). Therefore, silique number is proportional to total seed number for each dormancy treatment.

### Assessing Between-Year Dormancy and Seed Survival in the Soil

Even though we selected lines that differed primarily in seasonal dormancy, it is also possible that they may have differed in between-year dormancy as well. However, given that the field populations naturally cycled, we had no way of precisely estimating the frequency of between-year dormancy in the population cages. To address this limitation, we set up seedbank pot-cages for the Italy x Sweden RIL set. Each pot-cage represented an environmental treatment (Control, High Humidity, Native Soil) and had 2 clay pots (26.9cm diameter each), one with dormant seeds and the other with non-dormant seeds. There were 48 pots total, 2 dormancy treatments x 3 environmental treatments x 8 replicates (embedded into the same block system as the population cages). On May 9, 2021 (start of year 3 of the population cage experiment), 114 seeds were planted per pot with 2 replicates per homozygous line and 9 replicates per heterozygous line (same lines used in the population cages), resulting in about 15% heterozygosity per pot, as was used in the population cages (Table S2). Germination was monitored weekly in all pots until December 2022 to estimate the frequency of between-year dormancy (for more details see Supplementary Information). Seeds that did not germinate were assumed to have died, giving estimates of seed mortality in the soil. Note that we did not explicitly evaluate seed viability, so our estimate of seed mortality could include both viable and inviable seeds.

### Assessing Germination Timing in the Field

To confirm that the differences in seasonal dormancy in the lab germination assays translated to germination timing differences in the field, we calculated seasonal germination proportions both from the population cages and SeedBank pot-cages. The number of germinants in each season was divided by the total number of germinants in that year for each population or pot. Seasons were set according to the timing of peaks of germination: Summer: mid-March (Julian date = 78) to late September (Julian date = 264); Early Autumn: late September (Julian date = 265) to late October (Julian Date = 298); Late Autumn: late October (Julian date = 299) to mid-March (Julian date = 77). Spring was incorporated into Summer and Winter was incorporated into Late Autumn due to low germination during those seasons. A chi-square contingency test (PROC CATMOD) was used to test for differences in the season of germination between dormancy and RIL/environmental treatments for each year. Due to low variation in seasonal germination timing in year 3, only years 1 and 2 were analyzed for the Control treatment in the population cages. Similarly, due to low overall germination and low variation in seasonal germination timing, year 2 was excluded from the SeedBank pots analysis.

### Projections of population growth rate and sensitivities

To estimate long-term persistence, or the asymptotic lambda, we constructed a 2-year structured population projection matrix model. The model includes two years of vital rates to take into account contributions from the seed bank and to allow for manipulations of environmental variability across years. The annual census is envisioned to be just after new seeds are dispersed. Specifically, total population size (N) is equal to the sum of “fresh/new” seeds (N_f_) and below-ground seeds in the seed bank (N_sb_), resulting in a 2 x 2 projection matrix (Fig. S3). Fresh seeds survive until germination time and germinate at rate G_f_, and those that do not (fraction 1 – G_f_) go into the seed bank. Seeds survive in the seed bank at rate (S_seed_) and germinate out of it at rate G_sb_. Seeds that germinated in a given year survive to reproduction at rate S_rep_ and individuals that survived to reproduction produce F seeds. Reproductive success (w) is equal to S_rep_*F. For more details on the model’s framework and the calculation of vital rates, see the Supplementary Information.

To examine the consequences of having a seed bank and the effects of environmental variation, we had 4 model scenarios. Environmental variation was represented by “favorable” and “unfavorable” years. The year 1 and year 2 measurements of reproductive success from the population cages were used for “favorable” and “unfavorable” years, respectively. Scenarios 1 and 2 had two favorable years of survival and reproduction in a row, with scenario 1 having no between-year dormancy and scenario 2 with the maximum between-year dormancy, as estimated from the seed-bank pots. Scenarios 3 and 4 had 1 favorable year followed by an unfavorable year of survival and reproduction, with scenario 3 having no between year dormancy and scenario 4 having the maximum between-year dormancy. An average matrix was computed for each dormancy*environmental treatment and model scenario, based on field census data. The popbio package in R was used to calculate the expected asymptotic population growth rate (λ) and matrix element sensitivities for all dormancy*environmental treatments and model scenarios (Caswell 2001; Stubben and Milligan 2007).

For an estimate of variation in lambda, we computed bootstrapped 95% confidence intervals for each dormancy*treatment*model scenario using the boot package in R (Canty *et al*. 2024). The bootstrap sampled each block, with replacement, to calculate the average vital rates for each dormancy*treatment combination and then used those averages to calculate the lambdas for each model scenario. The function boot.ci was used with the “percentile” method to calculate the 95% confidence intervals (Carpenter and Bithell 2000).

## Results

### Differences in seed dormancy and germination timing of the founding populations

Four weeks after seed harvest, and four or fewer days after planting in the field, the Dormant populations had lower germination proportions than the Non-Dormant populations in all RIL sets under controlled conditions in the lab (It x Sw *X*^2^ =178.85, P < 0.001; Ger x Taj *X*^2^ = 101.54 P < 0.001; UK x USA *X*^2^ =27.36; P < 0.001), confirming that the populations differed significantly in dormancy at the time the populations were established in the field (Fig. S4).

In the field, Non-Dormant populations of Italy x Sweden had more germination in the Summer, and Dormant populations had more germination in the Autumn (Table S3,4; Fig. S5,6). This confirms that the dormancy treatments differed in seed dormancy in the lab and in seasonal germination timing the field.

### Differences in environments across the environmental treatments

The environmental treatments significantly affected soil temperature and moisture, with the Native Soil treatment having the most chronically stressful and the Control and High Humidity treatments having the most permissive environment (Table S5; Fig. S7A, B). The Native Soil treatment had the most extreme soil temperatures, with higher maxima and lower minima than the Control treatment, especially during the fall season. The Native Soil treatment was also the driest, especially in the fall to early spring seasons. The Native Soil treatment experienced drought conditions (Min SWC < 0.1) before all other treatments and it was the only treatment to experience drought in the Spring of year 1. In addition, the Native Soil treatment was the only treatment to experience temperatures below 6°C in year 1, the first to experience hotter temperatures (Max °C > 30) in year 2, and the last to transition to cooler temperatures in year 3. Therefore, the Native Soil treatment not only experienced more extreme temperatures and drier soils but also experienced hotter temperatures and drier soils sooner than the other treatments.

The High Humidity treatment was warm in the summer (significantly warmer than control), early fall, and spring, but moist from summer through winter. It was significantly moister than the other treatments for most of the experimental duration. The High Humidity treatment was the first treatment to experience soil saturation (Max SWC >0.3) in years 1 and 2, was the only saturated treatment in the spring of year 1, and the last treatment to transition to drier soil in the spring of year 2. The High Humidity was not just wetter than the other treatments but also experienced wetter conditions sooner and for longer durations. All treatments had similar PPFD, confirming that the roofs on the High Humidity treatment did not significantly alter light availability (Fig. S7C).

The Litter treatment was drier than the Control treatment in April 2021 but was otherwise similar to the Control treatment in both temperature and soil moisture. However, the Litter treatment was cooler than the Native Soil and High Humidity treatments on average. We could not quantify the potential light stress caused by the Litter treatment.

### Demographic Effects of Dormancy in the Control Treatment

#### Demographic transitions across life stages

*Seed to seedling transition:* Estimated seed mortality was high and between-year dormancy was low in all SeedBank pots (Table S6; Fig. S8B, C). Dormant populations had higher germination in year 2 than Non-Dormant populations (based on maximum germination estimates; Table S6, Fig. S8B). These results suggest that dormancy loci we selected for mostly influenced germination timing within a year but did not significantly influence between-year dormancy.

Through bet-hedging and phenological tracking, seed dormancy is expected to increase population size by reducing the risk of post-germination mortality. In the control treatment of the population cages, Dormant and Mixed populations had a larger class of seedlings than Non-Dormant populations in year 3 in the Italy x Sweden and UK x USA RIL sets (Table 1, S7; Fig. S9A). This likely reflects lower immediate mortality of new germinants in Dormant populations.

**Table 1.**
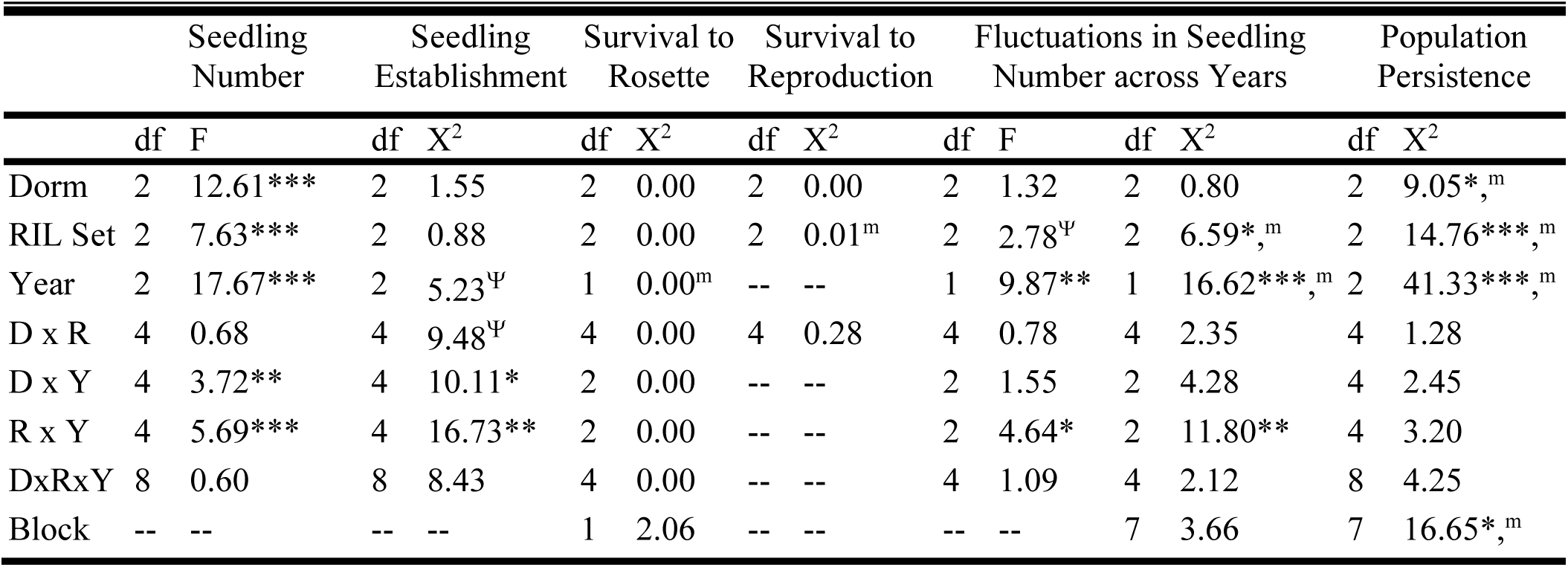
Effects of dormancy across all RIL sets in the control treatment. The effect of dormancy, RIL set, year, and all interactions among those factors for seedling number, seedling establishment, survival to rosette, survival to reproduction, fluctuations in seedling number between years, and population persistence. “Year” represents “Transition” (ex: year 1 to 2) for seedling number fluctuations and “Year-Season” for seedling establishment. Year 1, Summer was excluded from the analysis of seedling establishment since the Germany x Tajikistan and UK x USA RIL sets did not have any germination in that season; block was also excluded. Year 2 was excluded from the analysis of survival to reproduction due to very low survival to reproduction in all treatments that year; block was also excluded. The eight blocks were reduced to two “new blocks” for the survival to rosette analysis. Seedling number was analyzed using a type III ANOVA while the other traits were analyzed with a logistic regression. The seedling number fluctuations data was analyzed using both a type III ANOVA of the quantitative data and a logistic regression of the qualitative direction of the change. Population persistence was analyzed as a binomial, persist or not. P-values are represented as ***P < 0.001, **P < 0.01, *P < 0.05, ^Ψ^P <0.08. Significant main effects that were observed in the main-effect-only logistic regression models are represented as ^m^.

*Stage-specific survival:* Via phenological tracking, seasonal dormancy is also expected to increase survival throughout the life cycle. Dormant populations had higher seedling establishment than Non-Dormant populations in year 1 autumn, but the difference was only significant in the Germany x Tajikistan RIL (Table 1, S8; Fig. S9B). Dormant populations of Germany x Tajikistan also had higher survival to rosette than Non-Dormant populations in year 1 (Table S7; Fig. S9C). Dormancy did not affect survival to reproduction, at the population level, in any RIL set or year (Table 1, S7; Fig. S9D). Of note, survival to rosette and reproduction was low or zero across treatments in year 2.

Since our estimates of population-level stage-specific survival were limited to individuals growing inside the circle, we augmented those estimates by following individuals inside and outside the circle. Unlike the population-level results, individuals in Non-Dormant Italy x Sweden populations had lower survival to reproduction than the other dormancy treatments in the Control treatment (Table 3; Fig. 4A, Fig. S10). In addition, unlike the population-level results, individual survival to reproduction was greater than zero for most treatments in year 2, but survival to reproduction did generally decrease from year 1 to 2.

*Fecundity and total lifetime fitness:* As a form of habitat selection, seasonal dormancy may increase individual fitness. However, per-capita reproductive output did not vary significantly between dormancy treatments in the Italy x Sweden RIL set, the only RIL set in which it was measured (Table 3; Fig. 4B). Thus, the higher lifetime fitness in Dormant and Mixed Control populations, compared to Non-Dormant populations in this RIL set is primarily driven by the differences in survival to reproductive adult rather than per-capita fruit production (Fig. 4C).

#### Population stability and persistence

*Fluctuations in population size between years:* Through both bet-hedging and habitat selection, seasonal dormancy can buffer population size changes or reduce declines in population size across years. Changes in seedling number between year 1 and year 2 did not differ among dormancy treatments for any RIL set (Table 1; Fig. S9E). However, the difference in seedling number between years 2 and 3 did differ, such that Non-Dormant populations declined while Dormant and Mixed populations increased in Italy x Sweden (Table S7).

*Population persistence across years:* All the potential effects of seasonal dormancy on population size, stage-specific survival, and fluctuations in population size should combine to increase population persistence. In the control treatment, Non-Dormant populations had the lowest persistence over time (Table 1, S7; Fig. 6, Fig. S9F). Dormant and Mixed Italy x Sweden populations significantly persisted throughout the three years, in contrast to the Non-Dormant populations, which went extinct but reappeared in year 3. Thus, dormancy resulted in more consistent above-ground populations in the Italy x Sweden RIL set. In contrast, Non-Dormant populations had lower persistence than the other treatments only in year 1 for the Germany x Tajikistan RIL set. In line with the fitness results at the population and individual level, there was higher population failure in year 2.

In summary, Non-Dormant populations had smaller seedling classes than Dormant and Mixed populations by year 3, Non-Dormant populations had lower survival during early life stages than Dormant populations (Germany x Tajikistan in year 1), and Non-Dormant populations also had lower persistence than Dormant or Mixed populations.

### Demographic Effects of Dormancy in Different Environmental Treatments

With climate change increasing the occurrence of stressful environments it is important to know whether seasonal dormancy gives populations an advantage by allowing seedlings to avoid the most stressful conditions when conditions are poor in general, or whether stressful conditions repress the effects of dormancy. In contrast, seasonal dormancy may allow populations to take better advantage of permissive conditions by promoting seed germination under the best conditions, thereby giving dormancy a stronger effect in more favorable conditions than in more stressful conditions. Below we compare the demographic effects of seasonal dormancy that we found in the Control treatment to the other environmental treatments.

The Native Soil and Litter treatments were the most stressful, and the Control and High Humidity treatments were the most favorable environmental treatments. The Control and High Humidity treatments had the largest seedling classes with the Native Soil and Litter treatments with the smallest seedling classes (Table 2; Fig. 3A). In addition to small seedling classes, the Litter treatment also had the lowest survival across life stages, and the lowest population persistence over time (Fig. 3, 6).

**Figure 3.**
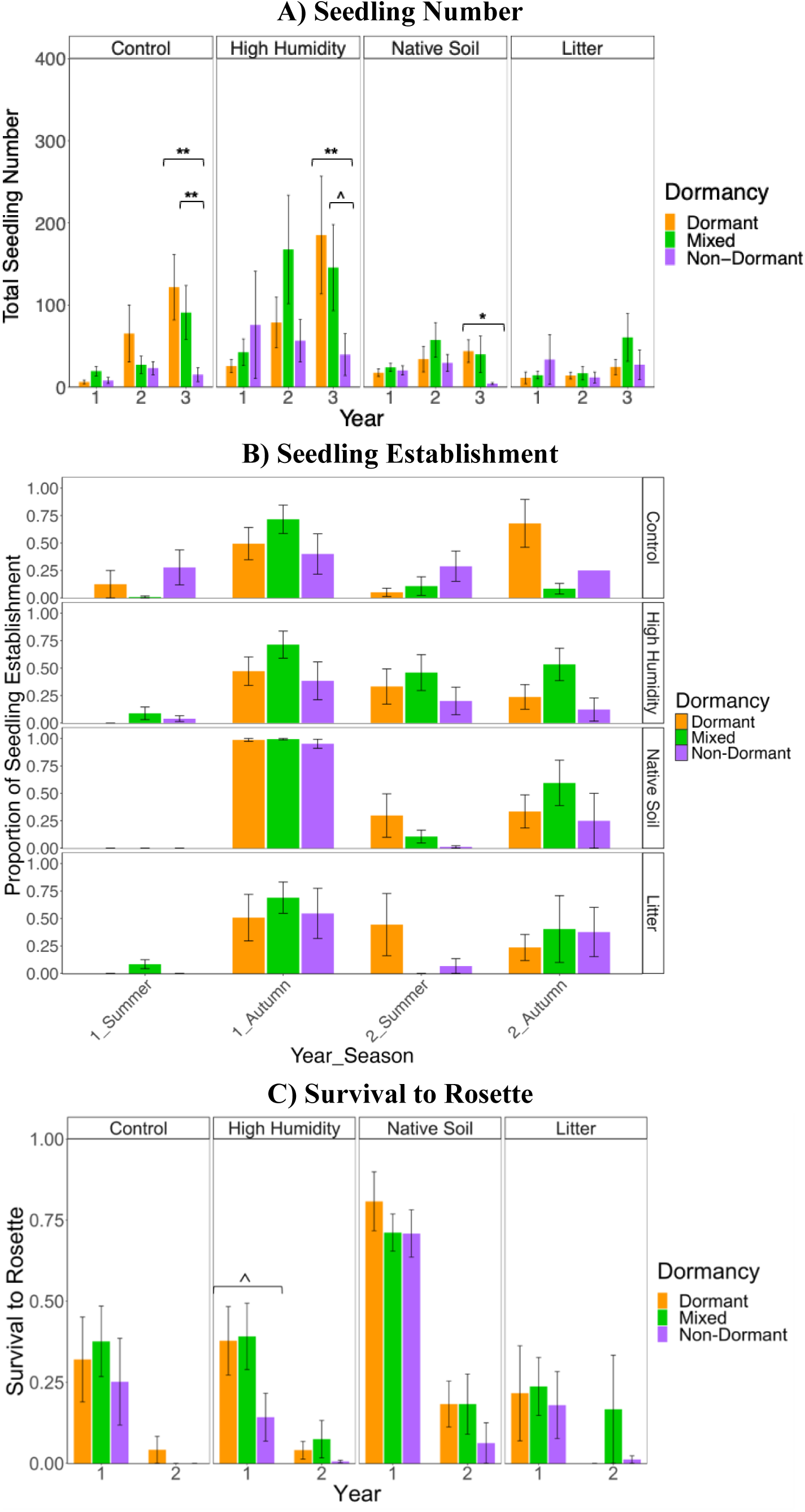
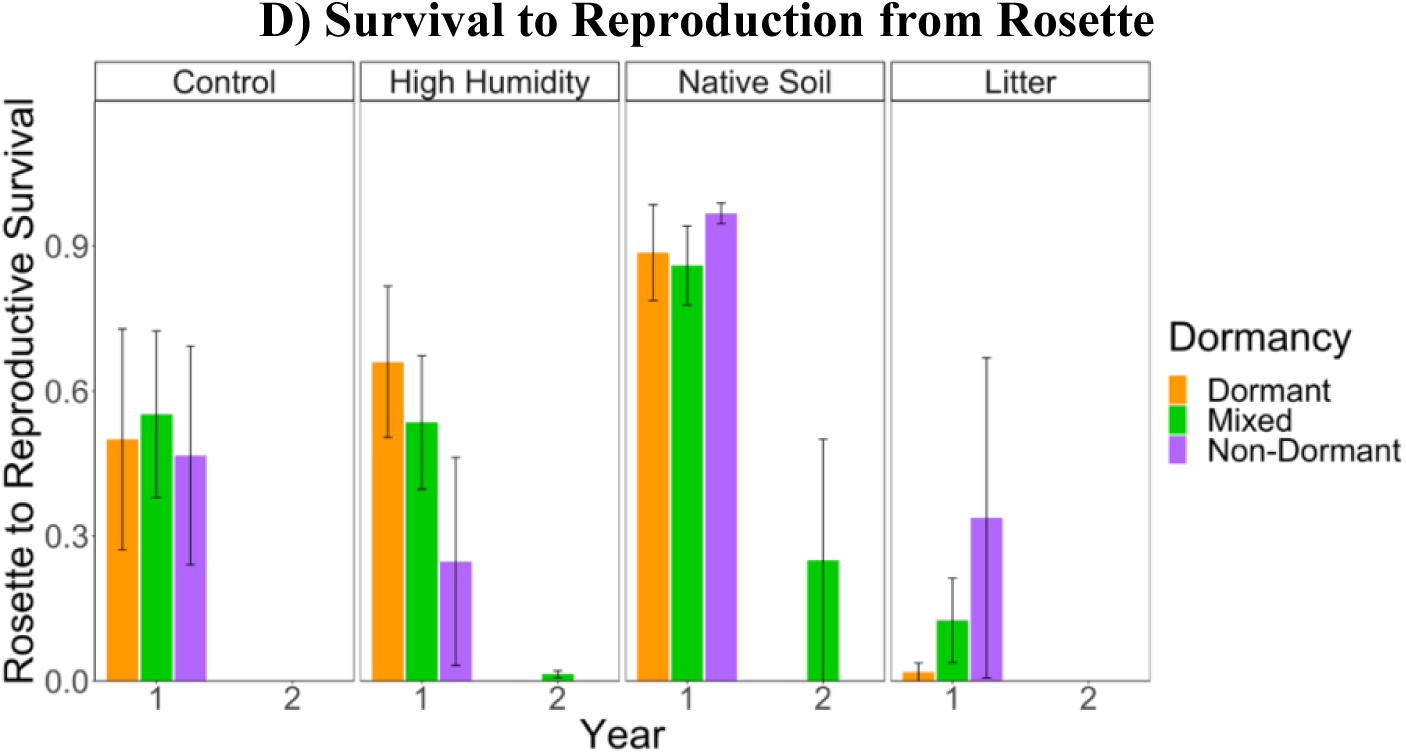
Key demographic metrics calculated from population-level data across environmental treatments for the Italy x Sweden RIL set. See Supplemental Information for results of the other RIL sets. Dormancy differences in A) the size of the seedling class, B) seedling establishment, C) proportion of survival to rosette, and D) proportion of survival to reproduction from rosette for each environmental. Dormant populations are in orange, Mixed in green, and Non-Dormant in purple. Raw means with standard errors are shown. Brackets with asterisks indicate significant differences among dormancy treatments. P-values are represented as ***P < 0.001, **P < 0.01, *P < 0.05, ^P<0.05 prior to correction test.

**Table 2.**
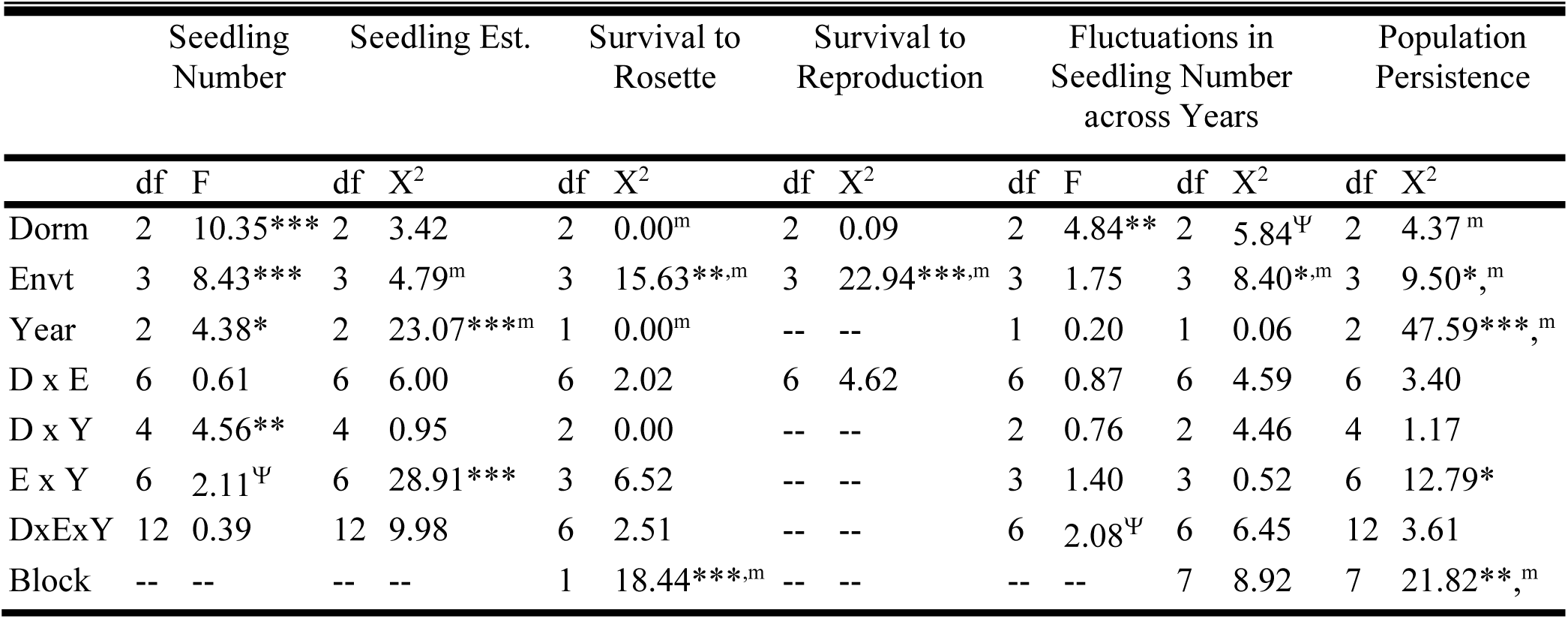
Effects of dormancy across environmental treatments in the Italy x Sweden RIL set. The effect of dormancy, environmental treatment, year, and all interactions among those factors for seedling number, seedling establishment, survival to rosette, survival to reproduction, seedling number fluctuations between years, and population persistence. “Year” represents “Transition” (ex: year 1 to 2) for seedling number fluctuations and “Year-Season” for seedling establishment; Year 1, Summer was removed from the seedling establishment analysis due to quasi-complete separation of data points issues with the model convergence; block was excluded as well. Year 2 was excluded from the analysis of survival to reproduction due to very low survival to reproduction in all treatments in year 2; block was excluded as well. The eight blocks were reduced to two “new blocks” for the survival to rosette analysis. Seedling number was analyzed using a type III ANOVA, and the other traits were analyzed with a logistic regression. The seedling number fluctuations data was analyzed using both a type III ANOVA of the quantitative data and a logistic regression of the qualitative direction of the change. Population persistence was analyzed as a binomial, persist or not. P-values are represented as ***P < 0.001, **P < 0.01, *P < 0.05, ^Ψ^P <0.08. Significant main effects that were observed in the main-effect-only logistic regression models are represented as ^m^.

The differences across dormancy treatments were more pronounced in the more favorable conditions (Control and High Humidity) than in the more stressful ones (Native Soil and Litter). As in the Control treatment, Non-Dormant High Humidity populations had smaller seedling classes by year 3, lower lifetime fitness, and lower population persistence than the other dormancy treatments (Table 2, 3, S9; Fig. 3A, 4C, 6). In addition, the effects of dormancy persisted later in life in the High Humidity treatment, with Non-Dormant populations having marginally lower survival from germinant to the rosette in year 1 than the other dormancy treatments (Table S9; Fig. 3C). Dormancy did increase the size of the seedling class in the Native Soil treatment, but Dormant populations remained small relative to Dormant populations in the Control and High Humidity treatments (Table S9; Fig. 3A). In addition, there were no differences in population persistence across dormancy treatments in the Native Soil treatment (Table S9; Fig. 6). The Litter treatment did not show any demographic differences among dormancy treatments, except with higher individual survival in Non-Dormant populations than the other dormancy treatments in year 2 (Table S9; Fig. 3, 4A, 5, 6).

**Figure 4.**
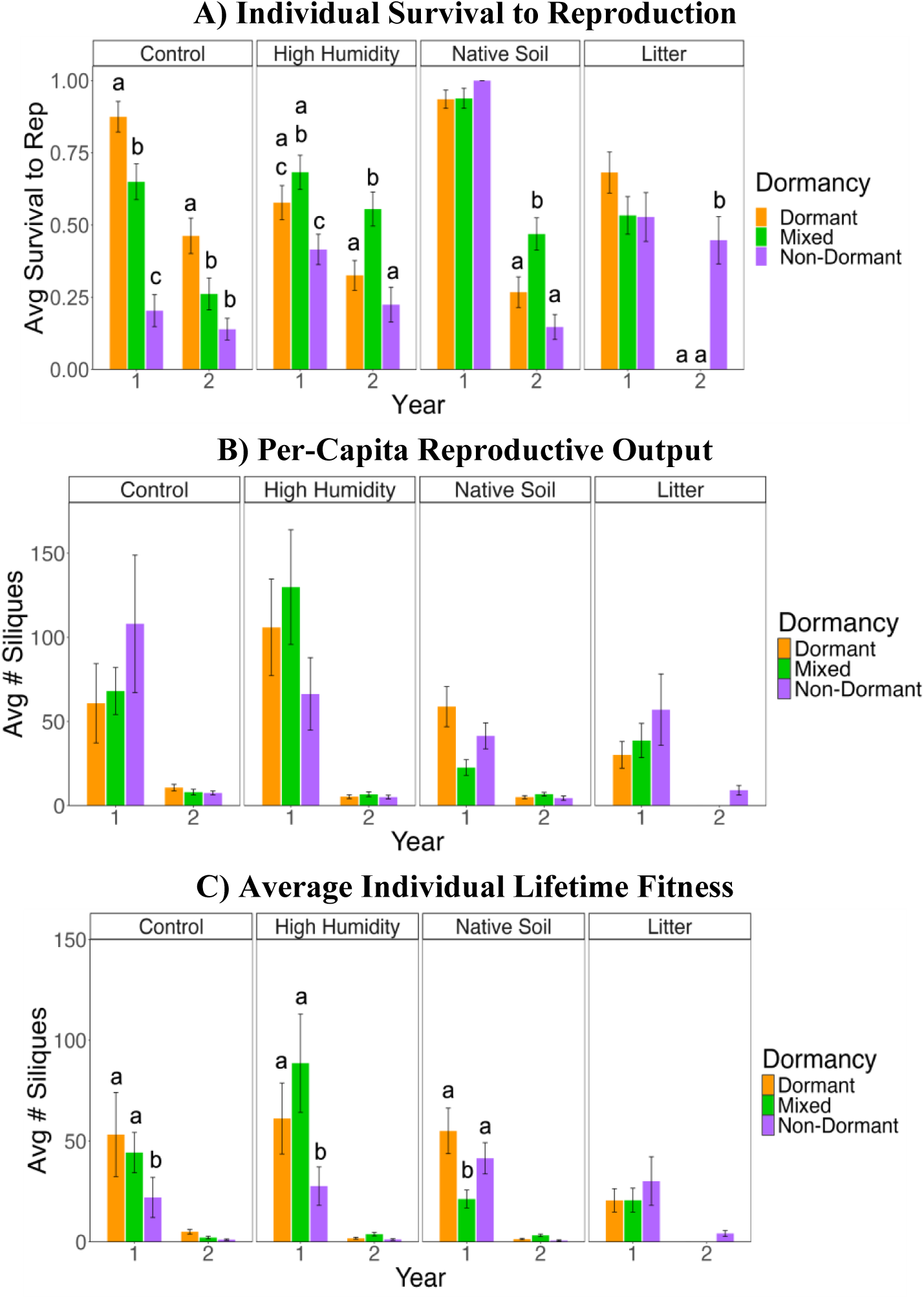
Individual A) survival to reproduction, B) per-capita reproductive output, and C) average lifetime fitness. Significant pairwise differences between dormancy treatments, within an environmental treatment are indicated by different letters.

**Figure 5.**
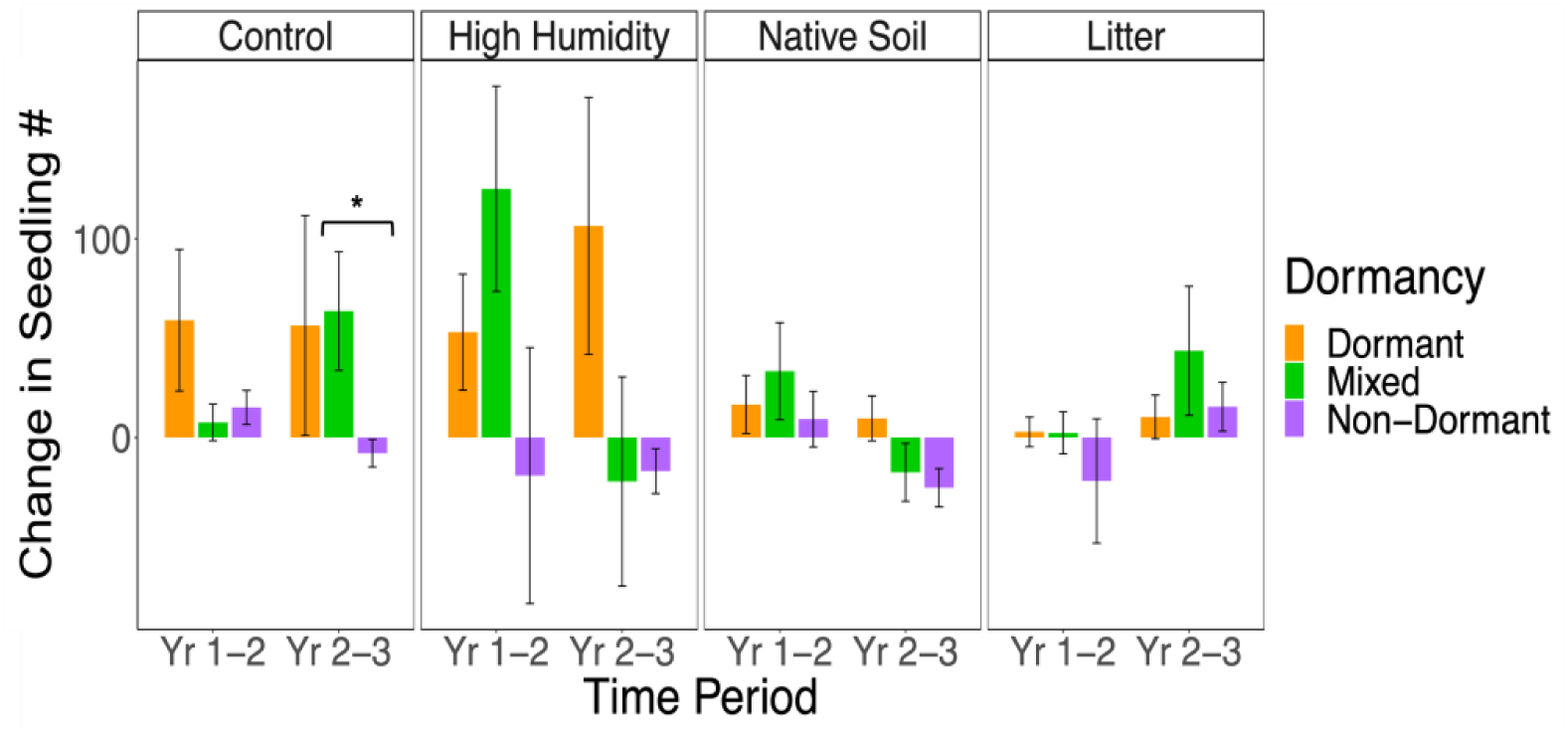
Fluctuations in the size of the seedling class between years, for each environmental treatment. The change in seedling number between years is shown for each environmental treatment in the Italy x Sweden RIL set. Bars above 0 represent an increase in seedling number and bars below 0 represent a decrease in seedling number. Dormant populations are in orange, Mixed in green, and Non-Dormant in purple. Raw means with standard errors are shown. P-values are represented as ***P < 0.001, **P < 0.01, *P < 0.05.

**Table 3.**
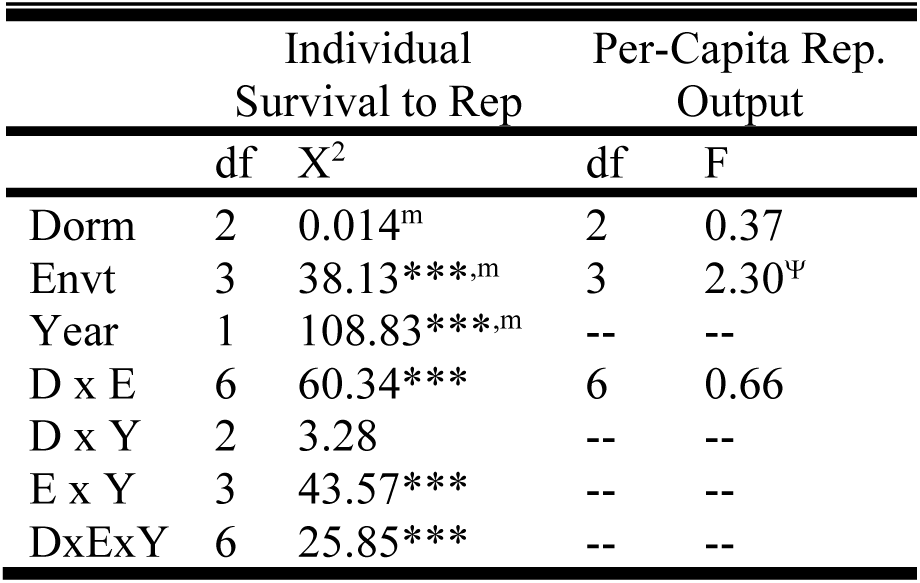
Effects of dormancy on individual-level fitness estimates in the Italy x Sweden RIL set. Year 2 was excluded from the analysis of per-capita reproductive output due to very low survival to reproduction in all treatments in year 2. Block was excluded from all models. Survival to reproduction was analyzed using a logistic regression and per-capita reproductive output was analyzed using a type III ANOVA. P-values are represented as ***P < 0.001, **P < 0.01, *P < 0.05, ^Ψ^P <0.08. Significant main effects that were observed in the main-effect-only logistic regression models are represented as ^m^. Note that there was no full model for individual lifetime fitness. The effect of dormancy was evaluated for lifetime fitness using Friedman’s tests within each year and environment.

|  | Individual<br>Survival to Rep |  | Per-Capita Rep.<br>Output |  |
| --- | --- | --- | --- | --- |
|  | df | X<sup>2</sup> | df | F |
| Dorm | 2 | 0.014<sup>m</sup> | 2 | 0.37 |
| Envt | 3 | 38.13***,<sup>m</sup> | 3 | 2.30<sup>Ψ</sup> |
| Year | 1 | 108.83***,<sup>m</sup> | -- | -- |
| D x E | 6 | 60.34*** | 6 | 0.66 |
| D x Y | 2 | 3.28 | -- | -- |
| E x Y | 3 | 43.57*** | -- | -- |
| DxExY | 6 | 25.85*** | -- | -- |

### Effects of Dormancy on Response to Environmental Variation

If dormancy stabilizes population performance under environmental variation, we expect that the effect of the environmental treatments would be less pronounced in the Dormant populations than the Non-Dormant populations. Overall, the Native Soil and Litter treatments were the most stressful treatments, with both Litter and Native Soil having the lowest seedling class sizes, and the Litter treatment having the highest population extinction in years 2 and 3. Dormancy was not able to counteract the detrimental effects of those environments on population demography. However, despite mostly non-significant interactions between dormancy and the environment, we detected significant effects of environmental variation in seedling number, survival to rosette, and per-capita reproductive output in Dormant and Mixed populations, but not in Non-Dormant populations, (Table S10). Therefore, contrary to expectation, Dormant populations had greater variation in performance across environments, and this was because they were able to perform better in favorable environments than Non-dormant populations.

**Figure 6.**
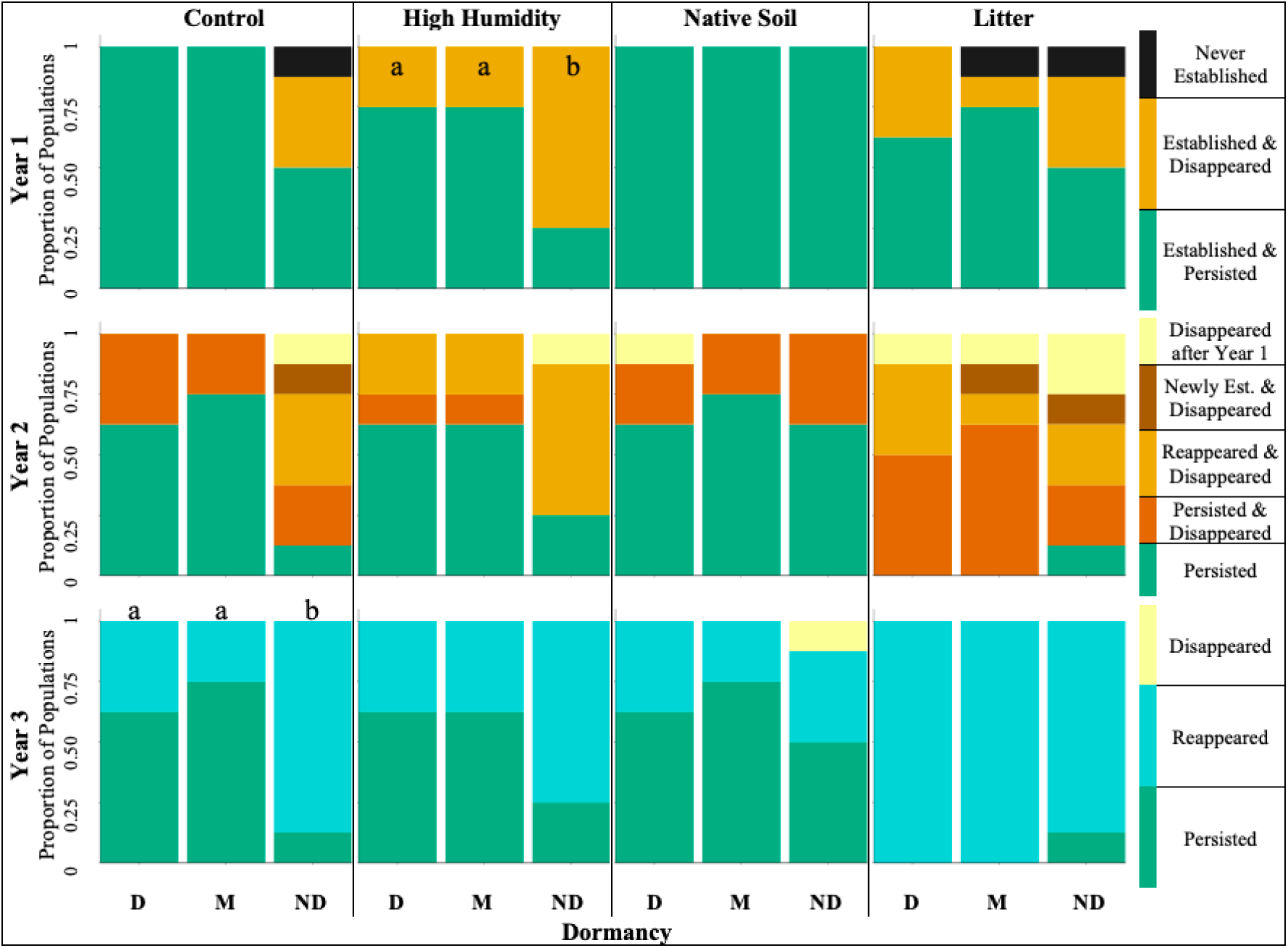
Above-ground population persistence across environmental treatments. The proportion of populations that persisted (green shading) or went extinct (warm shading) is shown for each environmental treatment in the Italy x Sweden RIL set. Note that there are multiple ways/timepoints a population could have gone extinct and that some populations showed signs of seed-bank dynamics by “reappearing” after going extinct in a previous year (see Figure 3 for more details on the classifications). Also, note that for year 3, there is no data on whether populations with germination also successfully persisted to reproduction, as data collection stopped in the winter of year 3. Letters indicate significant differences between dormancy treatments.

### Demographic Projections of Population Growth Rates as a Function of Between-year Dormancy and Environmental Variation

The above demographic dynamics were potentially the combined effect of seasonal and between-year dormancy. Here we explicitly compare the expected demographic outcomes with versus without between-year dormancy with population models parameterized with the above demographic data. In the population projection models, Non-Dormant populations had lower population growth rates than populations with dormancy (Mixed or Dormant) under almost all conditions (Table S11; Fig. 7). For the scenario that most closely resembles what occurred during our experiment, with fluctuations in environmental quality across years and a seed bank (model 4), Dormant populations had higher expected population growth than Non-Dormant populations in all environmental treatments, except the Litter treatment. It is important to note that population growth was greater than 1 for all dormancy treatments in both models 1 and 2, which had consistently favorable conditions, except in the Litter treatment.

**Figure 7.**
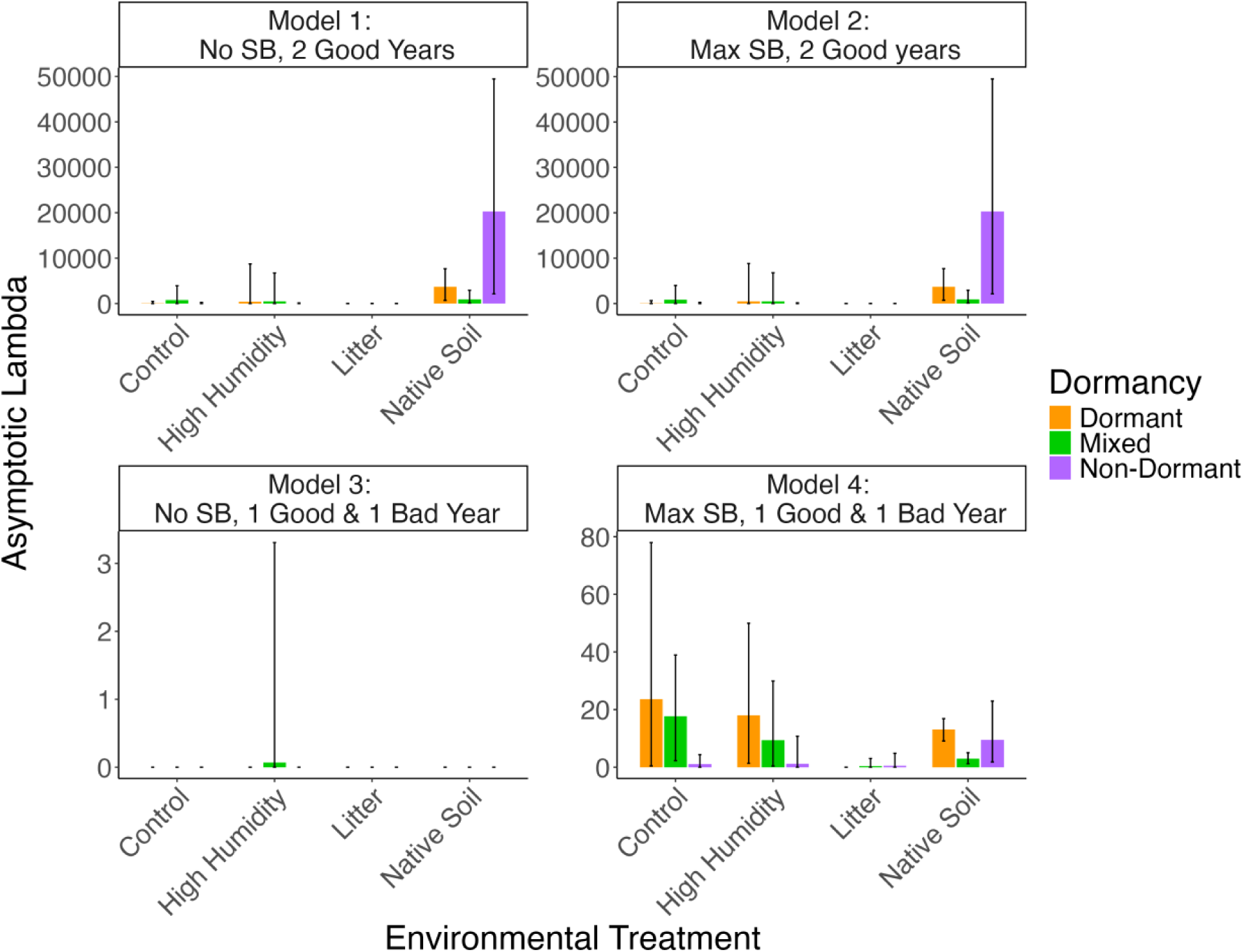
Asymptotic population growth rate from the population projection models. Each panel shows the results of one of the model scenarios. Error bars are bootstrapped 95% confidence intervals.

Seed banking increased population growth rates, especially when environments varied from year to year. The benefit of between-year dormancy is showcased by the 0 asymptotic lambdas found for most dormancy-by-environment treatments, for the scenario with environmental variation without between-year dormancy (model 3). In the scenarios with seed banking (models 2 and 4), the size of the below-ground seed population (lower matrix elements) had a stronger influence on population growth rate than the size of the above-ground population (upper matrix elements). In stable environments (model 2), the contribution of fresh seeds to the seed bank was the most influential, but in variable environments (model 4) below-ground seeds that came from plants that had a history of seed banking was most influential.

## Discussion

With climate change posing a threat to plant species persistence, it is important to investigate whether, and under what conditions, traits may facilitate population persistence. This study shows that seed dormancy can significantly increase population persistence, primarily by enhancing the performance of early life stages. However, environments that induce stress, especially early in life, may prevent dormancy from facilitating favorable population demography. The dormancy treatment affected seasonal seed dormancy, with Non-Dormant populations having more germination in the summer and Dormant populations having more germination in the fall. However, all dormancy treatments had low between-year dormancy. Therefore, the results discussed here primarily pertain to seasonal seed dormancy. Interestingly, the effects of dormancy also depend on genetic background (RIL sets). Furthermore, dormancy’s effects on early life stages may have resulted in dormancy evolving in the Mixed populations. By increasing population persistence and population size, dormancy may help stabilize population demography across years.

### Dormancy Positively Influenced Population Demography

As expected, dormancy increased population persistence, thereby reducing extinction risk (Willis *et al*. 2014). Dormant populations tended to have larger seedling classes and higher survival from seedling to rosette than Non-Dormant populations. These results align with the expectation for dormancy to increase survival by facilitating the tracking of seasonal environmental conditions (Donohue *et al*. 2010) and contribute to the growing empirical evidence of early life effects of dormancy (e.g. (González-Astorga and Núñez-Farfán 2000; Lu *et al*. 2016; Postma and Ågren 2022). However, some studies have also found a tradeoff between the early survival benefits conferred by dormancy and late life survival and/or fecundity (González-Astorga and Núñez-Farfán 2000; Postma and Ågren 2022). In our study, there was no apparent tradeoff between early life and late life traits. In fact, in the High Humidity Italy x Sweden populations and Germany x Tajikistan Control populations, Dormant populations had higher survival to reproduction than Non-Dormant populations. This result is consistent with the prediction that dormancy permits plants to germinate into more favorable conditions. The lack of more widespread dormancy differences in survival from rosette to reproduction and per-capita reproductive output could be due to the unavoidable caveat of reduced sample sizes at later life stages, and by the fact that the reproductive timing becomes synchronized later in life. More work is needed to assess how seed dormancy affects later life stages.

The strongest effect of dormancy was on total seedling number. The size of the seedling class is the consequence of both germination propensity and survival of germinants until the time of census. Thus, higher observed seedling numbers in Dormant populations in year 3 could be from higher germination in that treatment compared to the Non-Dormant treatment. This would be counter to expectations as non-dormant seeds should be able to germinate across a wider range of conditions and thus have higher germination proportions. A higher germination proportion conferred by the dormant treatment was also observed in the seed-bank experimental results. Note that due to the differences in when the experiments were planted, the first year of the seed-bank pot experiment is the third year of the population-cage experiment. Thus, the higher germination proportions of fresh Dormant seeds compared to Non-Dormant seeds in the pots align with the year 3 population cage results. However, it is also possible that cryptic episodes of germination and very early seedling mortality were missed between censuses, and that Non-Dormant populations tended to germinate more under unfavorable conditions that would cause mortality between censuses, leading to lower observed germination proportions. Consequently, lower seedling numbers likely indicates higher immediate mortality in Non-Dormant populations. This hypothesis is supported by a higher proportion of germination in the Non-Dormant populations occurring in the summer, a time when both soil temperature was high, and soil moisture was low.

### Dormancy Effects were Stronger in more Favorable Environments

Dormancy had a larger effect on population demography in the more permissive Control and High Humidity treatments. Overall, these results highlight the importance of measuring dormancy differences across multiple environments. Native Soil and Litter were the most stressful treatments as seedling number was low in both treatments and fitness estimates were low in the Litter treatment. In addition, the Native Soil treatment experienced the most extreme soil temperature and moisture conditions. It is possible that in stressful environments, the effects of dormancy are more suppressed. Notably, seedling number was higher in Dormant Native Soil populations than Non-Dormant populations in year 3. This suggests that dormancy was still being expressed in that environment and that the lack of dormancy effects at the later life stages was not due to that environment affecting the expression of dormancy, as has been found in previous experiments (Donohue *et al*. 2005b; Lu *et al*. 2016). Dormancy simply may not have been able to counter the adverse effects of the extreme conditions present in the Native Soil treatment. This suggests that dormancy enhances demographic performance not by allowing seeds to escape the most stressful conditions in overall poor environments, but by ensuring they germinate in the best conditions in more favorable environments.

### The Effects of Environmental Variation were more Pronounced in Dormant Populations

Theory suggests that dormancy should mitigate the effects of environmental variation (D’Aguillo *et al*. 2019). However, in this study we found that Dormant populations varied more across environmental treatments than Non-Dormant populations. This unexpected result could be due to the fact that Dormant populations were better able to take advantage of favorable conditions (Control and High Humidity treatments), which permitted a greater range of variation in fitness than in stressful environments. By germinating under the most favorable seasonal conditions in the permissive environmental treatments, Dormant populations tended to increase in population size while Non-Dormant populations tended to decrease, even under more permissive conditions.

### Dormancy may have Evolved in the Field

Natural selection is expected to favor seasonal dormancy in environments where hot, dry summers would reduce survival of non-dormant summer germinants. Our study contributes to previous studies by providing some evidence for positive selection on cued seed dormancy (Donohue *et al*. 2005b; Huang *et al*. 2010). First, Dormant populations had higher lifetime fitness than Non-Dormant populations in the Control and High Humidity treatments. In addition, across RIL sets, Mixed populations tended to resemble Dormant populations over time with respect to total seedling number, seedling establishment, and population persistence. This suggests that some selection on dormancy occurred in Mixed populations, favoring seed dormancy, and permitting them to attain higher demographic performance over time. This interpretation is consistent with lab observations (K Donohue, unpubl. res.) that the Mixed populations did evolve to become more dormant over the course of the experiment. Our results here demonstrate that, as dormancy of the Mixed populations became more like the Dormant populations, so too did their population demography.

### Dormancy Effects Differ across Genetic Backgrounds

It is also important to note that the demographic effects of dormancy differed across RIL sets. Importantly, the direction of the effect of dormancy did not differ across RIL sets, but the magnitude did. In general, Italy x Sweden populations performed better than the other RIL sets. This higher performance could be what allowed for the detection of significant differences in demography between dormancy treatments. Alternatively, since the genetic background and selected dormancy loci differed across RIL sets, seed dormancy may have not differed as strongly in the other RIL sets, reducing the magnitude of the effects of dormancy.

## Conclusion

Overall, our results show that populations of individuals with high seed dormancy would be expected to have more stable population demography, especially in more permissive environments. Less stable populations would be expected to have lower effective population sizes (Vucetich *et al*. 1997), which could increase the strength of genetic drift and the loss of genetic variation (Lande 1988; Lowe *et al*. 2017). By maintaining larger populations and decreasing drift, dormancy could potentially increase the opportunity for response to selection on other traits (Holt 1987). These effects on adaptation could then increase population growth and persistence. Future studies should directly investigate the relationship between dormancy, its effects on population demography, and any resulting indirect effects on adaptive dynamics. Seed dormancy may not only permit individuals to track changes in seasonal environmental conditions, but it may also facilitate a positive eco-evo feedback that increases population persistence or even establishment in novel environments.

## Supplementary Information

The supplementary information includes supplementary tables, figures, and methods. Supplemental methods provide additional details on the lab germination assays, calculation of seedling establishment, silique sampling methods, Seedbank Pots germination monitoring and analysis, and the population projection matrix. Supplementary tables and figures provide additional details related to the design of the study, statistics and figures for the other RIL sets (Germany x Tajikistan, UK x USA), and sub-models testing pairwise differences within treatments.

## Author Contributions

B.Q.C. conceived the ideas, designed the methodology, and collected and analyzed the data with assistance from K.D. Both authors contributed critically to the drafts and gave final approval for publication.

## Availability of Data and Materials

Data will be posted on Zenodo and GitHub upon publication.

## Acknowledgments

We thank the staff of the Duke Phytotron and Greenhouse facilities for the excellent care of the maternal plants for this experiment, their help with pasteurizing field soil, and help procuring all the supplies necessary for the experiment. Lindy Pittman, Nicholas Doak, Marley Kaplan, Jacob Carnes, Jonathan Pertile, Xavier Heidelberg, Yujin Kim, Caroline Fiore, Devan Wainright, Lester Sims, and Lesyia Smith-Sims helped substantially with field maintenance and data collection. We also thank the staff of the Duke Campus Farm who shared field maintenance equipment and storage space. William Morris provided invaluable feedback on the demographic model and early drafts of this manuscript.

## Literature Cited

1. Alonso-Blanco C, Bentsink L, Hanhart CJ, Vries HB, Koornneef M. 2003. Analysis of Natural Allelic Variation at Seed Dormancy Loci of *Arabidopsis thaliana*. Genetics 164: 711–729.

2. Baskin JM, Baskin CC. 1983. Seasonal Changes in the Germination Responses of Buried Seeds of *Arabidopsis thaliana* and Ecological Interpretation. Botanical Gazette.

3. Baskin JM, Baskin CC. 2004. A classification system for seed dormancy. Seed Science Research 14: 1–16.

4. Bentsink L, Hanson J, Hanhart CJ, et al. 2010. Natural variation for seed dormancy in Arabidopsis is regulated by additive genetic and molecular pathways. Proceedings of the National Academy of Sciences 107: 4264–4269.

5. Bertin RI. 2008. Plant phenology and distribution in relation to recent climate change. Journal of the Torrey Botanical Society 135: 126–146.

6. Burghardt LT, Metcalf CJE, Donohue K. 2016. A cline in seed dormancy helps conserve the environment experienced during reproduction across the range of *Arabidopsis thaliana*. American Journal of Botany 103: 47–59.

7. Campbell C. 2019. Why does my soil moisture sensor read negative? Environmental Biophysics.

8. Canty A, Ripley B, Brazzale AR. 2024. boot: Bootstrap Functions (Originally by Angelo Canty for S).

9. Carpenter J, Bithell J. 2000. Bootstrap confidence intervals: when, which, what? A practical guide for medical statisticians. Statistics in Medicine 19: 1141–1164.

10. Caswell H. 2001. Matrix population models: Construction, analysis, and interpretation. Sinauer Sunderland, MA.

11. Chaudhry S, Sidhu GPS. 2022. Climate change regulated abiotic stress mechanisms in plants: a comprehensive review. Plant Cell Reports 41: 1–31.

12. Cleland EE, Allen JM, Crimmins TM, et al. 2012. Phenological tracking enables positive species responses to climate change. Ecology 93: 1765–1771.

13. D’Aguillo M, Donohue K. 2023. Changes in phenology can alter patterns of natural selection: the joint evolution of germination time and postgermination traits. New Phytologist 238: 405–421.

14. D’Aguillo MC, Edwards BR, Donohue K. 2019. Can the Environment have a Genetic Basis? A Case Study of Seedling Establishment in *Arabidopsis thaliana*. Journal of Heredity 110: 467–478.

15. Debieu M, Tang C, Stich B, et al. 2013. Co-Variation between Seed Dormancy, Growth Rate and Flowering Time Changes with Latitude in *Arabidopsis thaliana*. PLOS ONE 8: e61075.

16. Donohue K. 2003. Setting the Stage: Phenotypic Plasticity as Habitat Selection. International Journal of Plant Sciences 164: S79–S92.

17. Donohue K, Dorn L, Griffith C, et al. 2005a. The Evolutionary Ecology of Seed Germination of *Arabidopsis thaliana*: Variable Natural Selection on Germination Timing. Evolution 59: 758–770.

18. Donohue K, Dorn L, Griffith C, et al. 2005b. Environmental and Genetic Influences on the Germination of *Arabidopsis thaliana* in the Field. Evolution 59: 740–757.

19. Donohue K, Rubio de Casas R, Burghardt L, Kovach K, Willis CG. 2010. Germination, Postgermination Adaptation, and Species Ecological Ranges. Annual Review of Ecology, Evolution, and Systematics 41: 293–319.

20. Evans MEK, Dennehy JJ. 2005. Germ Banking: Bet-Hedging and Variable Release From Egg and Seed Dormancy. The Quarterly Review of Biology 80: 431–451.

21. Falahati-Anbaran M, Lundemo S, Ågren J, Stenoien HK. 2011. Genetic consequences of seed banks in the perennial herb *Arabidopsis lyrata subsp. petraea* (Brassicaceae). American Journal of Botany 98: 1475–1485.

22. Finch-Savage WE, Leubner-Metzger G. 2006. Seed dormancy and the control of germination. New Phytologist 171: 501–523.

23. Furness AI, Lee K, Reznick DN. 2015. Adaptation in a variable environment: Phenotypic plasticity and bet-hedging during egg diapause and hatching in an annual killifish. Evolution 69: 1461–1475.

24. Gianella M, Bradford KJ, Guzzon F. 2021. Ecological, (epi)genetic and physiological aspects of bet-hedging in angiosperms. Plant Reproduction 34: 21–36.

25. González-Astorga J, Núñez-Farfán J. 2000. Variable demography in relation to germination time in the annual plant *Tagetes micrantha* Cav. (Asteraceae). Plant Ecology 151: 253–259.

26. Gremer JR, Kimball S, Venable DL. 2016. Within-and among-year germination in Sonoran Desert winter annuals: bet hedging and predictive germination in a variable environment. Ecology letters 19: 1209–1218.

27. Gremer JR, Venable DL. 2014. Bet hedging in desert winter annual plants: Optimal germination strategies in a variable environment (Y Buckley, Ed.). Ecology Letters 17: 380–387.

28. Holt RD. 1987. Population dynamics and evolutionary processes: the manifold roles of habitat selection. Evolutionary Ecology 1: 331–347.

29. Huang X, Schimitt J, Dorn L, et al. 2010. The earliest stages of adaptation in an experimental plant population: strong selection on QTLS for seed dormancy. Molecular Ecology 19: 1335–1351.

30. Iler AM, CaraDonna PJ, Forrest JRK, Post E. 2021. Demographic Consequences of Phenological Shifts in Response to Climate Change. Annual Review of Ecology, Evolution, and Systematics 52: 221–245.

31. Jagadish SVK, Bahuguna RN, Djanaguiraman M, Gamuyao R, Prasad PVV, Craufurd PQ. 2016. Implications of High Temperature and Elevated CO2 on Flowering Time in Plants. Frontiers in Plant Science 7.

32. Jessen M-T, Auge H, Harpole WS, Eskelinen A. 2023. Litter accumulation, not light limitation, drives early plant recruitment. Journal of Ecology 111: 1174–1187.

33. Kronholm I, Picó FX, Alonso-Blanco C, Goudet J, Meaux J de. 2012. Genetic Basis of Adaptation in *Arabidopsis thaliana*: Local Adaptation at the Seed Dormancy Qtl Dog1. Evolution 66: 2287–2302.

34. Lande R. 1988. Genetics and Demography in Biological Conservation. Science 241: 1455–1460.

35. Laserna MP, Sánchez RA, Botto JF. 2008. Light-related loci controlling seed germination in ler x Cvi and bay-0 x sha recombinant inbred-line populations of *Arabidopsis thaliana*. Annals of Botany 102: 631–642.

36. Leverett LD. 2017. Germination phenology determines the propensity for facilitation and competition. Ecology 98: 2437–2446.

37. Lowe WH, Kovach RP, Allendorf FW. 2017. Population Genetics and Demography Unite Ecology and Evolution. Trends in Ecology & Evolution 32: 141–152.

38. Lu JJ, Tan DY, Baskin CC, Baskin JM. 2016. Effects of germination season on life history traits and on transgenerational plasticity in seed dormancy in a cold desert annual. Scientific Reports 6: 25076.

39. Lundemo S, Falahati-Anbaran M, StenØien HK. 2009. Seed banks cause elevated generation times and effective population sizes of *Arabidopsis thaliana* in northern Europe. Molecular Ecology 18: 2798–2811.

40. Martínez-Berdeja A, Stitzer MC, Taylor MA, et al. 2020. Functional variants of DOG1 control seed chilling responses and variation in seasonal life-history strategies in *Arabidopsis thaliana*. Proceedings of the National Academy of Sciences of the United States of America 117: 2526–2534.

41. Mauricio R, Rausher MD. 1997. Experimental Manipulation of Putative Selective Agents Provides Evidence for the Role of Natural Enemies in the Evolution of Plant Defense. Evolution 51: 1435–1444.

42. Meng PH, MacQuet A, Loudet O, Marion-Poll A, North HM. 2008. Analysis of natural allelic variation controlling *Arabidopsis thaliana* seed germinability in response to cold and dark: Identification of three major quantitative trait loci. Molecular Plant 1: 145–154.

43. Miller ZR, Vasseur D, Hull PM. 2025. Stabilisation of Fluctuating Population Dynamics via the Evolution of Dormancy. Ecology Letters 28: e70141.

44. Montesinos-Navarro A, Picó FX, Tonsor SJ. 2012. Clinal Variation In Seed Traits Influencing Life Cycle Timing In *Arabidopsis thaliana*. Evolution 66: 3417–3431.

45. Penfield S. 2017. Seed dormancy and germination. Current Biology 27: R874–R878.

46. Philippi T, Seger J. 1989. Hedging one’s evolutionary bets, revisited. Trends in Ecology and Evolution 4: 41–44.

47. Picó FX. 2012. Demographic fate of *Arabidopsis thaliana* cohorts of autumn- and spring-germinated plants along an altitudinal gradient. Journal of Ecology 100: 1009–1018.

48. Postma FM, Ågren J. 2015. Maternal environment affects the genetic basis of seed dormancy in *Arabidopsis thaliana*. Molecular Ecology 24: 785–797.

49. Postma FM, Ågren J. 2022. Effects of primary seed dormancy on lifetime fitness of *Arabidopsis thaliana* in the field. Annals of Botany 129: 795–808.

50. Postma FM, Lundemo S, Ågren J. 2016. Seed dormancy cycling and mortality differ between two locally adapted populations of *Arabidopsis thaliana*. Annals of Botany 117: 249–256.

51. R Core Team. 2020. R: A Language and Environment for Statistical Computing. Vienna, Austria: R Foundation for Statistical Computing.

52. Seger J, Brockman HJ. 1987. What is bet-hedging? In: Harvey PH, Partridge L, eds. Oxford Surveys in Evolutionary Biology. Oxford: Oxford University Press, 182–211.

53. Sgrò CM, Terblanche JS, Hoffmann AA. 2016. What Can Plasticity Contribute to Insect Responses to Climate Change? Annual Review of Entomology 61: 433–451.

54. Stubben CJ, Milligan BG. 2007. Estimating and Analyzing Demographic Models Using the popbio Package in R. Journal of Statistical Software 22.

55. Taylor MA, Cooper MD, Sellamuthu R, et al. 2017. Interacting effects of genetic variation for seed dormancy and flowering time on phenology, life history, and fitness of experimental *Arabidopsis thaliana* populations over multiple generations in the field. New Phytologist 216: 291–302.

56. Thompson L. 1994. The Spatiotemporal Effects of Nitrogen and Litter on the Population Dynamics of *Arabidopsis thaliana*. Journal of Ecology 82: 63–68.

57. Tooke F, Battey NH. 2010. Temperate flowering phenology. Journal of Experimental Botany 61: 2853–2862.

58. Torres-Martínez L, Weldy P, Levy M, Emery NC. 2017. Spatiotemporal heterogeneity in precipitation patterns explain population-level germination strategies in an edaphic specialist. Annals of Botany 119: 253–265.

59. Tun W, Yoon J, Jeon J-S, An G. 2021. Influence of Climate Change on Flowering Time. Journal of Plant Biology 64: 193–203.

60. Venable DL. 2007. Bet hedging in a guild of desert annuals. Ecology 88: 1086–1090.

61. Vucetich JA, Waite TA, Nunney L. 1997. Fluctuating Population Size and the Ratio of Effective to Census Population Size. Evolution 51: 2017–2021.

62. Wickham H, Averick M, Bryan J, et al. 2019. Welcome to the tidyverse. Journal of Open Source Software 4: 1686.

63. Willis CG, Baskin CC, Baskin JM, et al. 2014. The evolution of seed dormancy: Environmental cues, evolutionary hubs, and diversification of the seed plants. New Phytologist 203: 300–309.

64. Wolkovich EM, Donahue MJ. 2021. How phenological tracking shapes species and communities in non-stationary environments. Biological Reviews 96: 2810–2827.

65. Zacchello G, Bomers S, Böhme C, Postma FM, Ågren J. 2022. Seed dormancy varies widely among *Arabidopsis thaliana* populations both between and within Fennoscandia and Italy. Ecology and Evolution 12: e8670.

66. Živković D, Tellier A. 2012. Germ banks affect the inference of past demographic events. Molecular Ecology 21: 5434–5446.

