## Supplemental methods, tables, and figures for "Seed dormancy increased population persistence in permissive environments, but not in stressful environments in an annual plant"

**Table S1. RIL sets and dormancy QTLs used to form Dormant and Non-Dormant populations. The names of the three RIL sets chosen, the locations the parents were sampled from, and the ABRC stock numbers are in the first two columns. The 2-3 dormancy QTLs, their additive effects on dormancy, and % variance in dormancy explained are shown alongside the source papers these data were derived from. The bottom half of the table lists the individual lines within each RIL set chosen for each dormancy treatment. For the Italy x Sweden RIL set, the dormant and non-dormant lines were split into two groups to facilitate the composition of the Mixed populations (see Table 2).**

| RIL Set | ABRC Stock # | Dormancy QTLs | Additive Effect | % Variance Explained | Source |
|---|---|---|---|---|---|
| Castelnuovo-12 x Rodasen-47 (Italy x Sweden) | CS98760 | D8 / DOG1 | 0.665 | 48.5 | Postma and Ågren, 2015 |
| | | D2 / CYP707A2 | 0.137 | 3.0 | |
| Bay-0 x Shahdara (Germany x Tajikistan) | CS57921 | DOG10 | -22.6 | 16 | Laserna et al. 2008 |
| | | DOG12 | -27.2 | 18 | |
| | | CDG-1 | -33.6 | 25 | Meng et al. 2008 |
| Cal-0 x Tac-1 (UK x USA) | CS97472 | QTL 1 / SNP223/Msat3.1 | -104.30 | 31.0 | Huang et al. 2010 |
| | | QTL 2 / SNP379 | -492.54 | 6.6 | |

| RIL Set | Dormancy | Individual Line Numbers |
|---|---|---|
| Italy x Sweden | Dormant | Group 1: CS98361, CS98394, CS98396, CS98421, CS98436, CS98441, CS98452, CS98467, CS98488, CS98522, CS98575, CS98593, CS98605, CS98606, CS98611, CS98612, CS98657, CS98658, CS98668, CS98686, CS98692, CS98719, CS98734, CS98740; Group 2: CS98354, CS98374, CS98378, CS98384, CS98398, CS98401, CS98415, CS98427, CS98492, CS98512, CS98514, CS98623, CS98625, CS98627, CS98640, CS98655, CS98661, CS98666, CS98671, CS98698, CS98706, CS98716, CS98739, CS98743 |
| | Non-Dorm | Group 1: CS98369, CS98376, CS98391, CS98416, CS98450, CS98462, CS98470, , CS98481, CS98486, CS98493, CS98504, CS98518, CS98571, CS98586, CS98600, CS98648, CS98652, CS98682, CS98708, CS98728, CS98732, CS98741, CS98744, CS98745; Group 2: CS98357, CS98360, CS98385, CS98392, CS98406, CS98407, CS98428, CS98476 CS98495, CS98496, CS98497, CS98503, CS98533, CS98539, CS98555, CS98572, CS98584, CS98591, CS98595, CS98618, CS98647, CS98660, CS98696, CS98731 |
| Germany x Tajikistan | Dormant | CS57521, CS57529, CS57562, CS57581, CS57613, CS57630, CS57633, CS57643, CS57649, CS57736, CS57768, CS57777, CS57788 CS57798, CS57800, CS57837, CS57860, CS57902, CS57917 |

| | | |
|---|---|---|
| | Non-Dorm | CS57524, CS57525, CS57556, CS57600, CS57611, CS57631, CS57640, CS57763, CS57771, CS57773, CS57812, CS57833, CS57836, CS57849, CS57871, CS57875, CS57887, CS57895, CS57914 |
| UK x USA | Dormant | CS97479, CS97485, CS97493, CS97497, CS97498, CS97501, CS97509, CS97514, CS97519, CS97520, CS97529, CS97530, CS97537, CS97558, CS97561, CS97573 |
| | Non-Dorm | CS97478, CS97484, CS97486, CS97488, CS97496, CS97499, CS97505, CS97510, CS97515, CS97518, CS97521, CS97524, CS97546, CS97547, CS97550, CS97555 |

**Table S2. Composition of founding population cages and seedbank pots. The number of available dormant and non-dormant homozygous recombinant inbred lines and the number of heterozygous lines produced from crosses within the Dormant/Non-Dormant populations are shown for each RIL set. The number of seeds per line planted within each population cage and seedbank pot for each RIL set are also shown, with the resulting N. For the RIL sets with heterozygous lines, each population/seedbank pot was composed so that there would be 15% heterozygosity. The homozygous and heterozygous lines, together, essentially represent the number of genetically distinct individuals, i.e., the Ne, in each Dormant or Non-Dormant sub-population. Note, the Mixed populations were composed using both the dormant and non-dormant lines, with all the dormant lines and all the non-dormant lines from the Dormant and Non-dormant populations. Half of the Mixed populations were composed of 5 seeds from Group 1 and 6 seeds from Group 2 in Table 1 while the other Mixed populations got 5 seeds from Group 2 and 6 seeds from Group 1 in Table 1. Only one of the three RIL sets (Italy x Sweden) was used for the seedbank pots. The number of seeds per line planted was reduced to account for the smaller area of growth, resulting in a smaller N.**

| | Homozyg. Lines | Seeds per Homo Line | Heterozyg. Lines | Seeds per Het Line | Percent Heterozyg. | Total # Seeds (N) |
|---|---|---|---|---|---|---|
| **Dormant and Non-Dormant Population Cages** | | | | | | |
| Italy x Sweden | 48 | 11 | 2 | 47 | 15.1% | 622 |
| Germany x Tajikistan | 19 | 28 | 1 | 95 | 15.4% | 617 |
| UK x USA | 16 | 39 | 0 | 0 | 0 | 624 |
| **Mixed Population Cages** | | | | | | |
| Italy x Sweden | 96 | 5-6 | 4 | 24 | 15.4% | 624 |
| Germany x Tajikistan | 38 | 14 | 2 | 47 | 15.3% | 616 |
| UK x USA | 32 | 20 | 0 | 0 | 0 | 640 |
| **Dormant and Non-Dormant Seedbank Pots** | | | | | | |
| Italy x Sweden | 48 | 2 | 2 | 9 | 15.8% | 114 |

A)

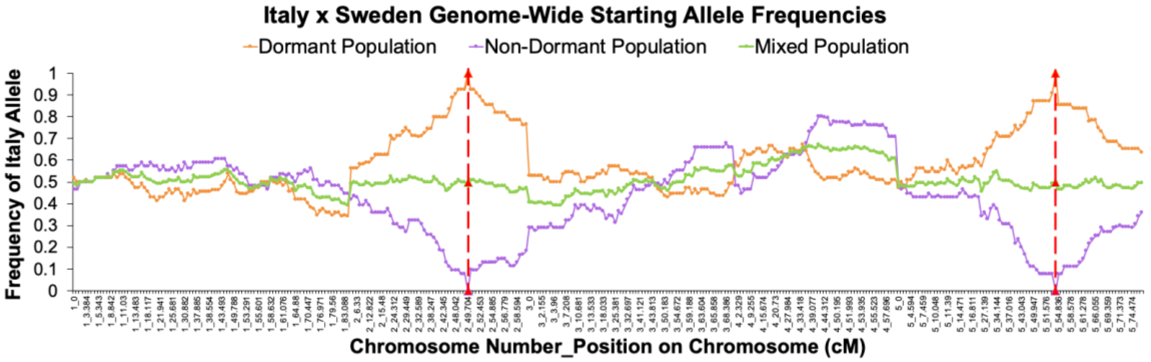

B)

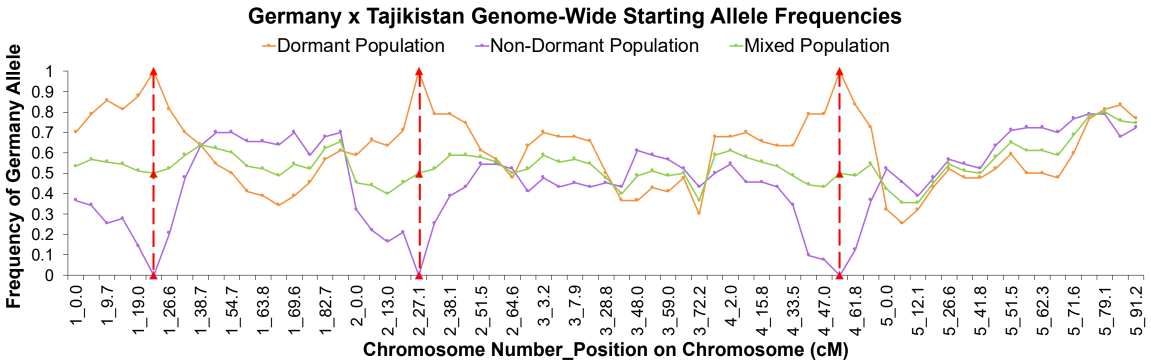

C)

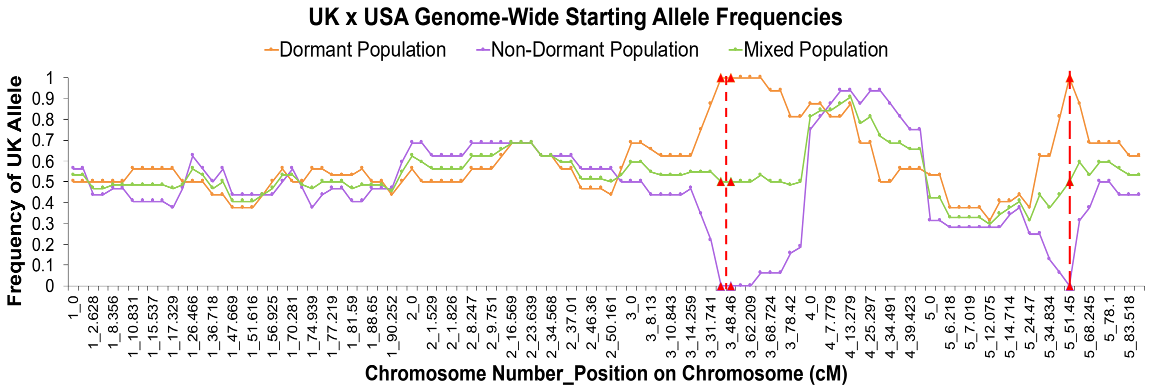

**Figure S1. Genome-wide starting allele frequencies for Italy x Sweden, Germany x Tajikistan, and UK x USA populations. Allele frequencies are based on the published SNP genotypes for each RIL that was used to compose the populations and based on the number of seeds per RIL shown in Table 2. Homozygous and heterozygous genotypes were accounted for appropriately in the allele frequency calculations. Dormant populations are in orange, Non-Dormant populations are in purple, and Mixed populations are in green. Red triangles and dotted lines represent a dormancy QTL of major effect that was selected to compose the experimental populations. A) Frequency of the Italian allele across the genome in the Italy x Sweden RIL set, B) frequency of the German allele across the genome in the Germany x Tajikistan RIL set, and C) frequency of the UK allele across the genome in the UK x USA RIL set. In all RIL sets, allele frequencies are close to 50% in all 3**

**population types, except near a dormancy locus. Allele frequencies at dormancy loci were made to be 100%, 50%, or 0% for the Dormant, Mixed, and Non-Dormant populations, respectively.**

### Lab Germination Assay Methods

Seeds were assayed at three temperatures, Optimal (18°C), Cold (10°C), Above Optimal (26°C). Each line within the Dormant and Non-Dormant population pools provided 20 seeds that were placed on a 35-mm petri plate containing 0.7% agar. There were three replicate plates per genotype per temperature treatment (9 plates per genotype). Plates were placed in Percival Model GR41LX incubation chambers (Percival Scientific Inc., Perry, IA, USA) with 12-hour days. Seeds were monitored for germination 4 days, 2 weeks, and 4 weeks after seeding. Germination was scored as radicle protrusion, and seed viability evaluated by assessing firmness to touch. Germination proportion was calculated as the total number of germinants divided by the number of viable seeds, with petri plate as the unit of replication. A logistic regression with a logit-link was used to test for differences in germination proportion between treatments within each RIL set (PROC LOGISTIC, SAS 9.4 SAS Institute, Inc. 2015).

### Methods to Calculate Seedling Establishment

During the germination season, censuses were conducted either weekly or bi-weekly, so the method with which seedling establishment was calculated depended on the time between censuses (Figure S2). Both methods of calculation estimated seedling establishment as the probability of survival for two-weeks from the time of germination, or survival from germination to the rosette stage, whichever came first. This probability was based on the number of new germinants that were observed two weeks prior that had either remained seedlings two weeks later or that had transitioned into rosettes; in addition, we considered the number of new rosettes that appeared between census that could not be attributed to a seedling from the prior census ("rapid rosettes"; Fig. S2) and added these to the number of germinants that survived to establishment.

**When there were two weeks between censuses**, a set of if/else statements in R was used to calculate the number of survivors (Fig. S2a). They are as follows: 1) IF the previous census's (Census(t-1)) new seedling number was greater than the number of new rosettes at Census(t) AND less than or equal to the sum of seedlings and new rosettes at Census(t), then all new seedlings at Census(t-1) survived and the proportion of seedling establishment is equal to 1 (Scenario 1 in Fig. S2a); ELSE 2) IF the number of new seedlings at Census(t-1) was greater than the number of new rosettes at Census(t) AND greater than the number of seedlings and new rosettes at Census(t), then some seedlings from Census(t-1) died, the survivors are the sum of the number of the seedlings and new rosettes at Census(t), and the proportion of seedling establishment is less than 1 (Scenario 2 in Fig. S2a); ELSE 3) IF the number of new seedlings at Census (t-1) was less than the number of new rosettes at Census(t), then the number of survivors is equal to the number of new seedlings at Census(t-1) plus the number of rapid rosettes and the proportion of seedling establishment is equal to 1 (Scenario 3 in Fig. S2a).

**When there was only one week between censuses**, the total number of survivors was calculated as the sum of the number of new seedlings at Census(t-2) that survived to Census(t-1) as rosettes, and those that remained seedlings at Census(t-1) and survived to Census(t) (Fig. S2b). In the case of weekly censuses, it is possible that some new rosettes at Census(t-1) could have come from slow growing germinants from Census(t-3). Thus, the contribution of "old seedlings," or germinants from Census(t-3) had to be distinguished from the contribution of "new seedlings," or germinants from Census(t-2). Thus, the number of new seedlings that transitioned to rosettes at Census(t-1) was the difference of the number of new rosettes at Census(t-1) and old seedlings still alive at Census(t-2); if that was negative, then the number of new seedlings that transitioned to rosettes at Census(t-1) was equal to 0 (Scenario 1 in Fig. S2b). Note that this scenario is unlikely for bi-weekly censuses since a seedling from Census(t-2) would be 4 weeks old by Census(t). The number of new seedlings at Census(t-2) that remained seedlings over 1 week (Census (t-1)) and then survived to the following week (Census(t)) was estimated using two scenarios: 1) IF the number of new rosettes at Census(t) is less than the number of seedlings that remained seedlings between Census(t-2) and Census(t-1), then some new seedlings died and the number of survivors is equal to the number of new rosettes at Census(t) (Scenario 2 in Fig. S2b); 2) ELSE, all the new seedlings that remained seedlings between Census(t-2) and Census(t-1) survived to Census(t) (Scenario 3 in Fig. S2b).

The total seedling establishment for summer (early June-early September) and autumn (late September-early December) for each population was calculated as the total number of survivors within that season divided by the total number of seedlings and rapid rosettes for each year.

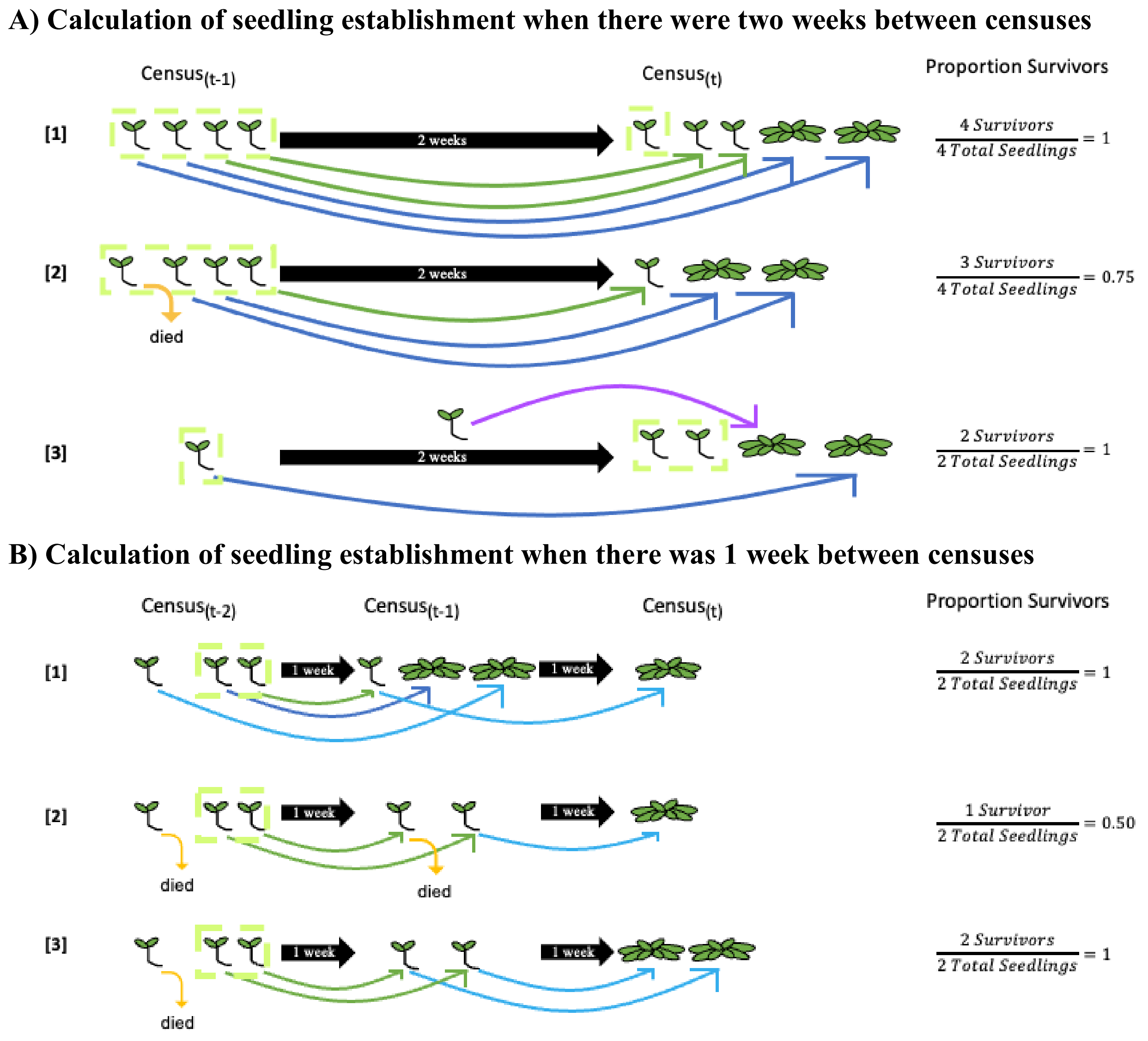

**Figure S2. Depiction of how seedling establishment was calculated from the population census data. A) Examples of how seedling establishment was calculated when there were two weeks between censuses, and B) examples of how seedling establishment was calculated when there was one week between censuses. The numbers on the left side of each row correspond to the scenarios described in the text. "Proportion Survivors" shows the number of survivors at Census(t) in the numerator and the number of new seedlings at Census(t-1) in A) and Census(t-2) in B). Note that for the 2-week censuses, survival is tracked for new seedlings from Census(t-1) and for the 1-week censuses, survival is tracked for new seedlings from Census(t-2). Light green boxes indicate new germinants observed during that census. Dark blue arrows represent seedlings that transitioned to rosettes between censuses; green arrows represent seedlings that remained seedlings between censuses; yellow arrows indicate seedlings that died; purple arrows represent seedlings that germinated and**

**transitioned to rosettes between censuses (rapid rosettes) which only occurred in two-week censuses; and light blue arrows represent seedlings that transitioned to rosettes at least two weeks after germinating. <u>Only new rosettes are shown.</u> A seedling was considered "established" if it survived as a seedling for two weeks or transitioned to rosette at any time within that 2-week window.**

### Silique Sampling Methods for Counting Seed Number

We collected siliques from up to six individuals in one Dormant and Non-Dormant population in each of 4 of the blocks and counted seeds in up to 5 siliques per individual (4 populations per dormancy treatment x 6 individuals per population x 5 siliques per individual = 120 siliques per dormancy treatment). All Dormant and Non-Dormant population pairs (from the same block) were sampled from the same environmental treatment. Native Soil populations were sampled in 3 out of 4 of the blocks and High Humidity populations were sampled in the $4^{th}$ block.

### Seedbank Pots Germination Monitoring and Analysis Methods

During each germination census in the Seedbank pots, the number of confirmed new seedlings (seedlings that were confirmed to be *A. thaliana*) was recorded, and those seedlings were removed from the pot. For seedlings for which we were not 100% confident were *A. thaliana,* we marked them with a toothpick and small bird band for future follow-up. Once those seedlings were big enough to be confirmed *A. thaliana,* or some other species, they were recorded and removed from the pot. This consistent monitoring and removal of young plants, along with the fabric around the cages, reduced the chances that new seeds came in from outside the pots. All seedlings' germination dates were recorded as the day they were first found in the field (regardless of when they were confirmed to be *A. thaliana)*. In contrast, when rosettes (4 true leaves) were found in the pots their germination date was recorded as the week prior to when they were first found, as their germination was likely more than a week earlier. Only 47 rosettes were found over the course of the two years.

To account for seedlings that were missing (died) after they were marked but not confirmed as *A. thaliana*, we analyzed the data in two ways, 1) a confirmed total number of seedlings, and 2) a maximum number of seedlings. The confirmed number of seedlings only included seedlings that were confirmed to be *A. thaliana*, either when they were first found or after being marked for follow-up. The maximum number of seedlings included any seedlings that were marked for follow-up and not found the following week (due to rapid mortality or soil shifting).

For the analysis, seeds could have three fates: germinate in year 1, germinate in year 2, or don't germinate in either year. Seeds that did not germinate after year 2 were assumed to have died. We calculated the proportion of each seed category out of the total number of seeds planted for each seed-bank pot. Note, that we cannot distinguish between seeds that died during the first year from those died during the second year. The proportions of the seed fates were calculated using both the confirmed and maximum estimates of seedlings that germinated in each pot. Year 1 was classified as May 9, 2021, through May 9, 2022, year 2 was then May 13, 2022, through December 9, 2022. These classifications match the lifecycle expressed by *A. thaliana* in our field site.

The effect of dormancy, environmental treatment, dormancy*environment, and block on the proportions of seeds that germinated in year 1 or 2, or died, in the seedbank pots was evaluated using a cumulative logit logistic regression (PROC LOGISTIC). Sub-models were used to assess pairwise differences between the Dormant and Non-Dormant treatments within each environmental treatment.

### Population Projection Matrix Details

We constructed 2 x 2 matrices to analyze the demographic dynamics of populations with different treatments and scenarios of seed dormancy (Figure 3). Matrix element [1,1] represents how fresh seeds at the start of year 1 contribute to the number of fresh seeds in year 3 by either germinating, surviving, and reproducing in years 1 and 2 or not germinating in year 1, surviving in the seedbank, germinating out of the seedbank, and surviving and reproducing in year 2 (Fig. 3B). Matrix element [1,2] represents how seeds in the seed bank at the start of year 1 contribute to the number of fresh seeds in year 3 by germinating out of the seed bank, surviving, and reproducing in year 1, and germinating as fresh seeds, surviving, and reproducing in year 2. Matrix element [2,1] represents how fresh seeds at the start of year 1 contribute to the number of seed bank seeds in year 3 by germinating, surviving, and reproducing in year 1, but not germinating in year 2 and surviving as a seed in the seed bank. Finally, matrix element [2,2] represents how seeds in the seed bank at the start of year 1 can contribute to the number of seed-bank seeds in year 3 by germinating, surviving, and reproducing in year 1, but not germinating in year 2 and surviving in the seed bank.

The vital rates were estimated from a combination of data from the population-cage field experiment and the seedbank pot-cages. The germination rate of fresh seeds ($G_f$) was estimated as the maximum number of seedlings inside the circle of the population cages in year 1 divided by the estimated number of seeds dispersed into the circle of the population cages ($Seeds_{Circle}$). The maximum number of seedlings estimate assumes that all seedlings died between censuses, and that any seedlings observed during a census represents a new germinant; therefore, every seedling observed during a census was counted as new germinants at each census. Given the potential for fast-growing plants that germinated in between censuses, all new rosettes were also counted as new germinants for this maximal estimate. The maximum number of seedlings was calculated as the sum of new germinants and new rosettes (assuming mortality between censuses) across all censuses within a year. Since seedling number was monitored only inside the circle for year 1, we estimated the number of seeds dispersed into the circle ($Seeds_{Circle}$) using the following formula: $Seeds_{Circle} = Seeds_{Total} * \frac{Rosettes_{Circle}}{Rosettes_{Total}}$. $Seeds_{Total}$ represents the total number of seeds dispersed into the population (inside and outside the circle), $Rosettes_{Circle}$ represents the total number of rosettes inside the circle in year 1, and $Rosettes_{Total}$ represents the total number of rosettes inside and outside the circle in year 1. Survival to reproduction ($S_{rep}$) was estimated as the total number of reproductive individuals inside the circle of the population cages divided by the maximum number of seedlings inside the circle of the population cages. Fecundity (F), or the number of seeds produced by each individual, was estimated from the individual reproductive output data in the population cages. The average number of fruits for each population was multiplied by 29, the average number of seeds per silique found in the silique samples taken in year 1 of the field experiment. $S_{rep}$*F thus represents fitness (survival times mean number of fruits of those that survived; w). Since reproductive output was measured only in the Italy x Sweden populations (five blocks only), the other RIL sets were not included in the models. $G_f$ was a constant across years in the model, while favorable years in the model consisted of the $S_{rep}$ and F estimates from year 1 of the population cages and unfavorable years in the model consisted of the $S_{rep}$ and F from year 2 of the population cages.

$G_{sb}$, germination from the seed bank, was estimated as the proportion of germinants in year 2 of the seedbank pots experiment out of the remaining seeds that did not germinate in year 1. The maximum $G_{sb}$ calculated from across the seed bank pots (0.1233) was used for all dormancy*environment treatments in the model. Germination from the seed bank was low across all dormancy and environmental treatments (range: $8.2x10^{-5}$-0.1233) so we used the maximum estimate to test for the effects of having seed banking versus not, in general. Survival of seeds in the seedbank was estimated differently depending on the age of the seed. For seeds in the seed bank for their first year, $S_{seed}$ was estimated as the maximum number of seedlings that germinated in the seed bank pots divided by the total number of seeds planted in each pot. We assumed that seeds that did not germinate in their second Autumn died or were no longer viable, so $S_{seed}$ is equal to 0 for seeds in the seed bank for their second year. The average $S_{seed}$ was calculated for each dormancy*environment treatment and used in the model. Since we did not have a Litter treatment in the seedbank pots experiment, we estimated $S_{seed}$ for the Dormant and Non-Dormant Litter populations to be the average $S_{seed}$ for each dormancy treatment, across all the other environmental treatments. To estimate $S_{seed}$ for the Mixed populations of each environmental treatment, we used the average $S_{seed}$ for each environmental treatment, across the other dormancy treatments. $G_{sb}$ and $S_{seed}$ were constants across years in the model.

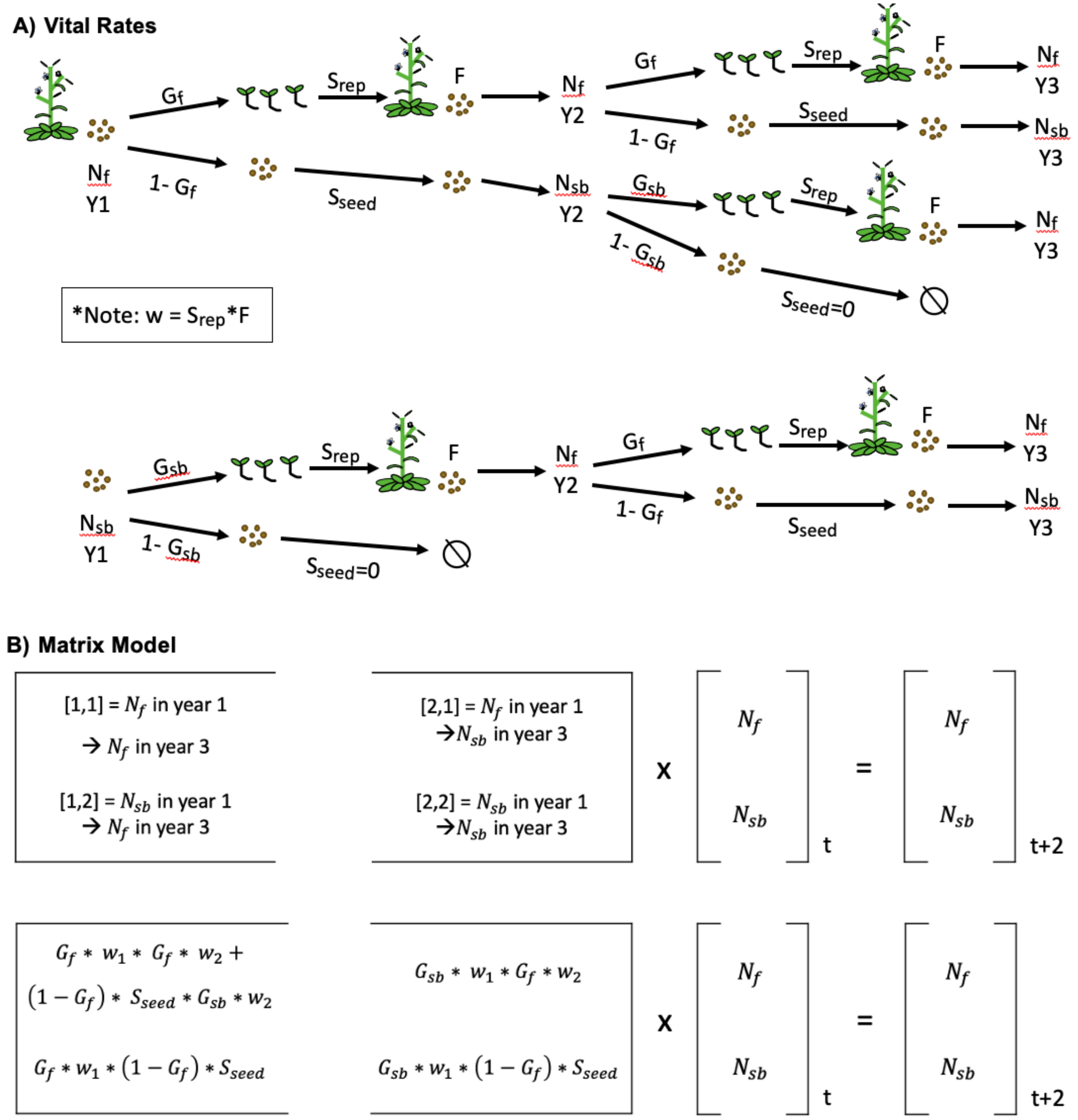

**Figure S3. Structure of the population projection matrix model. A) The vital rates included in the model. B) The 2x2 matrix that was created for each population to calculate lambda. The first row in B shows what each matrix element represents, and the second row shows how the vital rates were combined to represent the transition shown in the first row. The sum of $N_f$ and $N_{sb}$ is equal to the total population size. Individual matrices for each dormancy*environment combination were constructed by taking the average of the vital rates across population replicates for each treatment; the asymptotic population growth rate and sensitivities of the matrix elements were calculated for those matrices.**

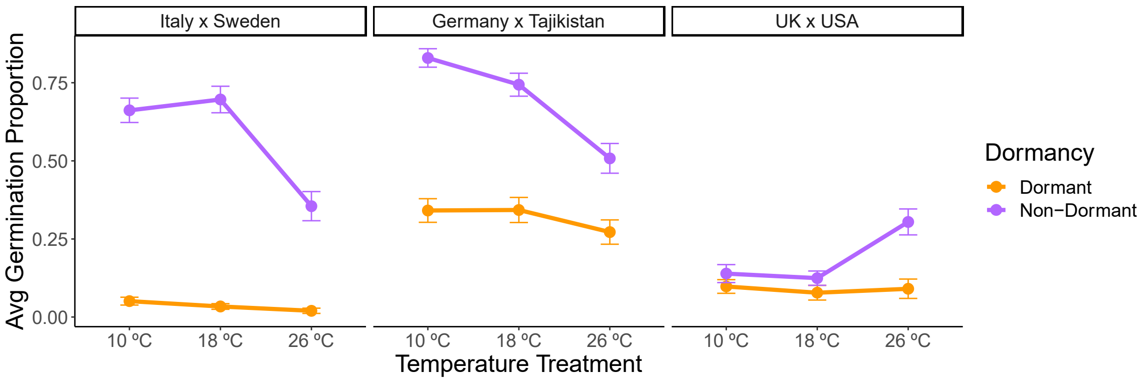

**Figure S4. Initial differences in germination between Dormant and Non-Dormant lines in the lab. Mean germination proportion after 4 weeks in all the Dormant and Non-Dormant lines/genotypes at the time of seed dispersal. Germination assays were conducted at 3 temperatures. For all RIL sets, Non-Dormant genotypes germinated more than Dormant genotypes (It x Sw $X^2$ =178.85, P < 0.001; Germany x Tajikistan $X^2$ = 101.55, P < 0.001; UK x USA $X^2$ =27.37, P < 0.001). Compared to the other RIL sets, the differences between dormancy treatments were smaller in UK x USA. The only significant differences in germination proportion were in the high temperature treatment for UK x USA, resulting in a dormancy*germination treatment effect ($X^2$=13.23, P=0.0013). In contrast, Germany x Tajikistan had smaller differences in germination in the high-temperature treatment compared to the other temperatures, resulting in a dormancy*temperature treatment effect ($X^2$=7.83, P=0.0199). Interestingly, dormancy effects were largest in the higher temperature for UK x USA compared to the other RIL sets where the differences were greatest in the lower temperature.**

**Table S3. Effects of dormancy and environmental treatments on seasonal germination in the population cages. A) The effect of dormancy, RIL set or environment, and the interaction between those effects on the season of germination. B) P-values for the pairwise differences between dormancy treatments within each RIL set and environmental treatment. Seasons were classified according to the germination peaks across time. P-values are represented as ***P < 0.001, **P < 0.01, *P < 0.05, $^{\Psi}$P < 0.08. The control treatment model was only run for years 1 and 2 due to low variation in seasonal germination in year 3. PROC CATMOD classified some effects as redundant due to zeros in the contingency table; chi-square statistics were not calculated for those effects (marked with "--").**

**A) Full Models**

| | Control Treatment All RIL Sets | | | | Italy x Sweden All Environmental Treatments | | | | | |
|---|---|---|---|---|---|---|---|---|---|---|
| | Year 1 | | Year 2 | | Year 1 | | Year 2 | | Year 3 | |
| | df | $X^2$ | df | $X^2$ | df | $X^2$ | df | $X^2$ | df | $X^2$ |
| Dormancy | 4 | 10.3* | 4 | 40.4*** | 4 | 48.9*** | 4 | 219.2*** | 2 | 15.0*** |
| RIL/Envt | -- | -- | 4 | 32.7*** | 6 | 176.6*** | -- | -- | 3 | 65.0*** |
| Dorm* RIL/Envt | 8 | 42.2*** | -- | -- | 12 | 98.7*** | -- | -- | 6 | 23.8*** |

**B) Pairwise Comparisons**

| | Control Treatment Within Each RIL Set | | Italy x Sweden Sweden Within Each Environmental Treatment | |
|---|---|---|---|---|
| | Italy x Sweden | | High Humidity | |
| | Year 1 | Year 2 | Year 1 | Year 2 |
| Dorm vs. NonDorm | <0.0001*** | 0.4358 | <0.0001*** | <0.0001*** |
| Dorm vs. Mixed | 0.0021** | -- | 0.0005*** | <0.0001*** |
| NonDorm vs. Mixed | 0.0272* | -- | <0.0001*** | <0.0001*** |
| | Germany x Tajikistan | | Litter | |
| | Year 1 | Year 2 | Year 1 | Year 2 |
| Dorm vs. NonDorm | 0.4555 | 0.7701 | <0.0001*** | -- |
| Dorm vs. Mixed | 0.0240* | 0.0040** | 0.0035** | -- |
| NonDorm vs. Mixed | 0.1541 | 0.0027** | <0.0001*** | <0.0001*** |
| | UK x USA | | Native Soil | |
| | Year 1 | Year 2 | Year 1 | Year 2 |
| Dorm vs. NonDorm | 0.0380* | 0.3364 | <0.0001*** | <0.0001*** |
| Dorm vs. Mixed | 0.0007*** | 0.6962 | <0.0001*** | <0.0001*** |
| NonDorm vs. Mixed | 0.4046 | 0.7741 | 0.0940 | <0.0001*** |

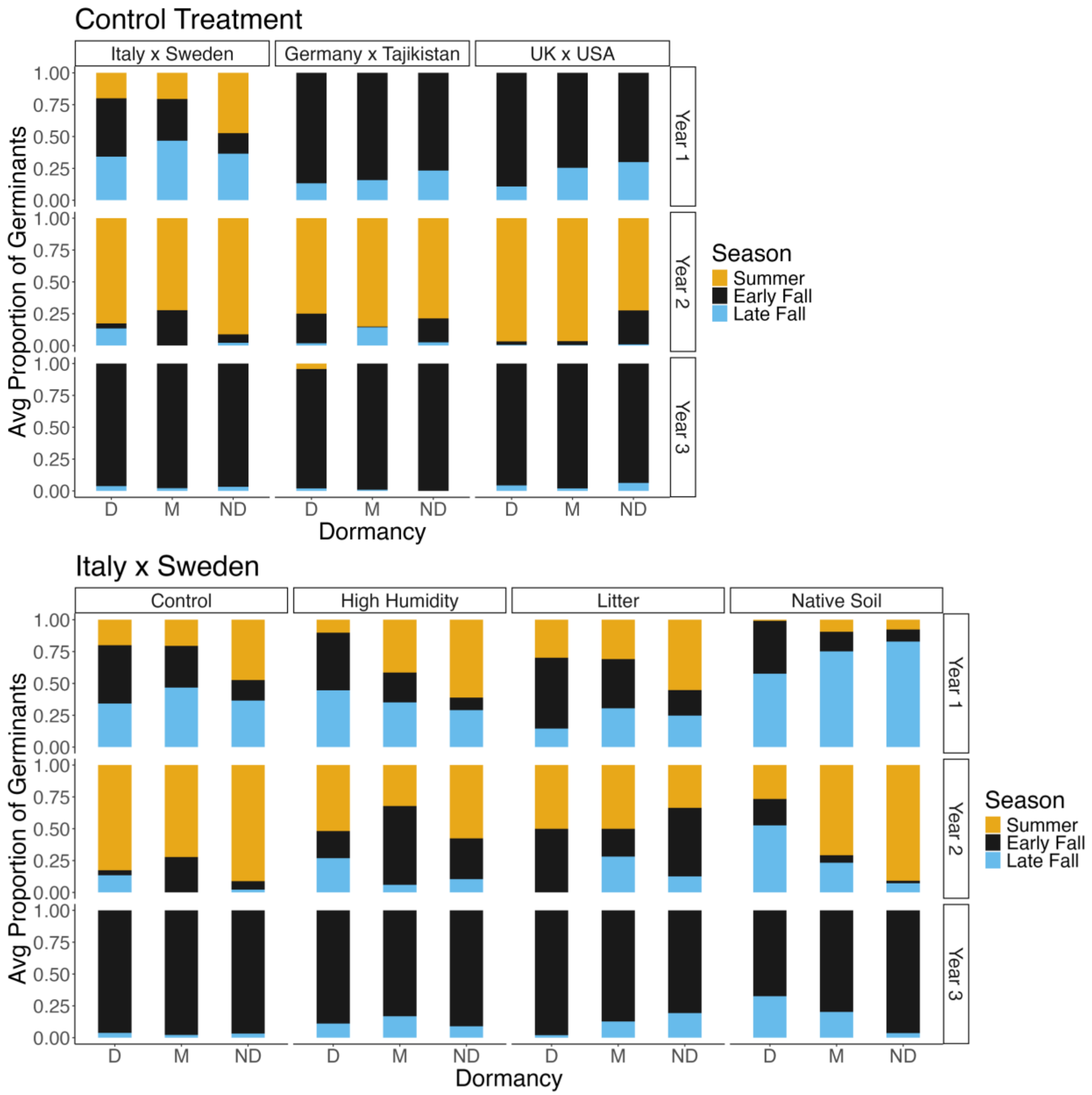

**Figure S5. Seasonal germination proportions in the population cages. The average proportion of germinants in each season is shown for the Control treatment in each RIL set and for each dormancy*environment treatment in the Italy x Sweden RIL set. The colors denote the season. The 'D' bars are for Dormant populations, 'M' bars for Mixed populations and 'ND' bars are for Non-Dormant populations.**

**Table S4. Effects of dormancy treatment (Dormant and Non-Dormant) and environmental treatment on seasonal germination in the SeedBank pots. A) The effect of dormancy, environment, and the interaction between those effects on the season of germination. B) P-values for the pairwise differences between Dormant and Non-dormant treatments within each environmental treatment. "Confirmed total" refers to the number of seedlings that were confirmed to be *A. thaliana* germinants; "Max total" includes the number of seedlings that were suspected of being *A. thaliana,* but which died too soon for definitive identification. P-values are represented as ***P < 0.001, **P < 0.01, *P < 0.05, $^{\Psi}$P < 0.08. The models were only run for year 1 since there was both low germination overall and low variation in seasonal germination in year 2.**

| **A) Full Models** | | | | |
|---|---|---|---|---|
| | Confirmed Total | | Max Total | |
| | df | $X^2$ | df | $X^2$ |
| Dormancy | 2 | 14.2*** | 2 | 16.4*** |
| Envt | 4 | 49.2*** | 4 | 69.5*** |
| Dorm*Envt | 4 | $8.7^{\Psi}$ | 4 | 17.6** |

| **B) Pairwise Comparisons Between Dormancy Treatments** | | |
|---|---|---|
| | Confirmed Total | Max Total |
| Control | **<0.0001*** ** | **<0.0001*** ** |
| High Humidity | **<0.0001*** ** | **<0.0001*** ** |
| Native Soil | 0.6642 | **0.0457*** |

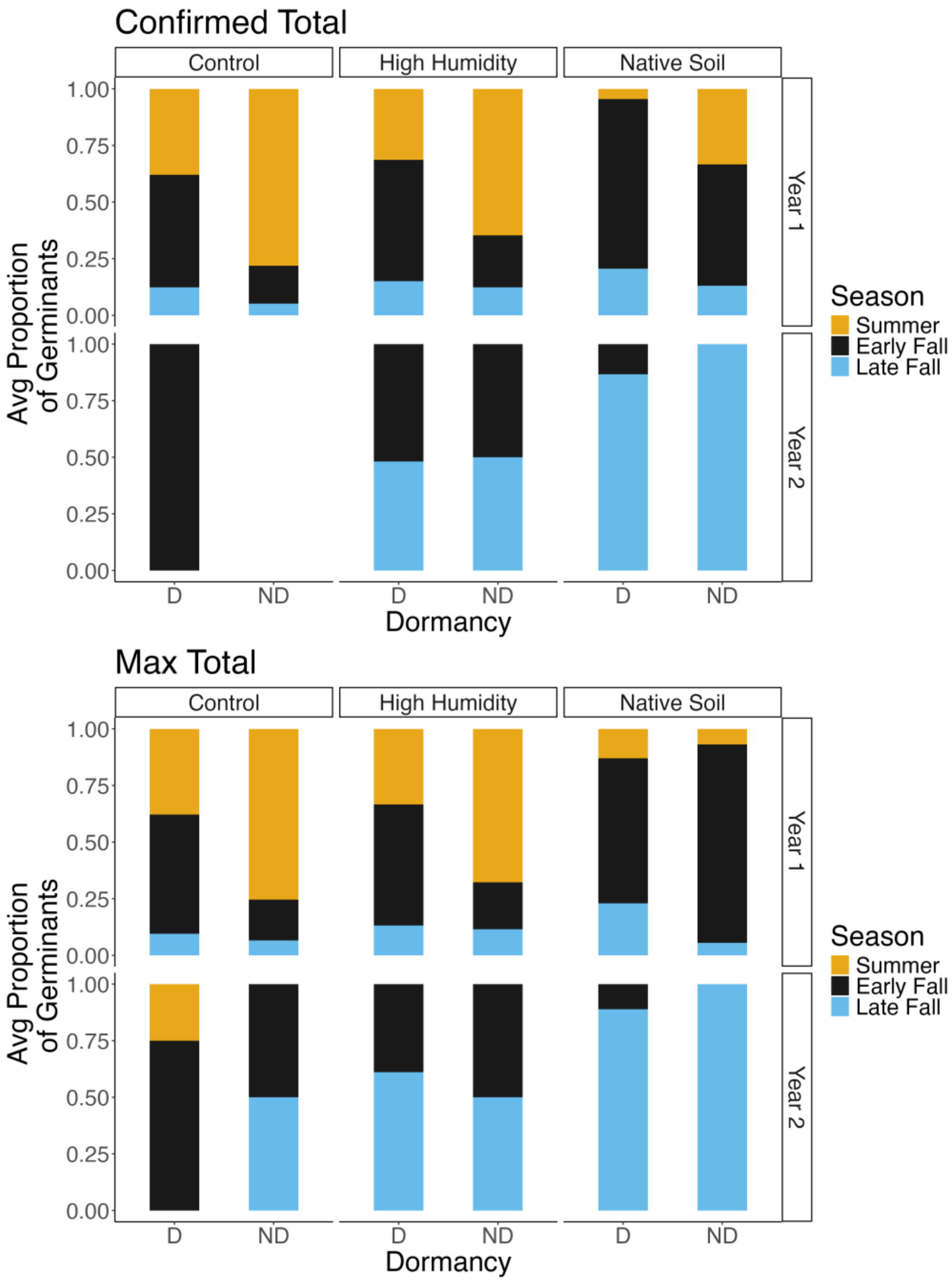

Figure S6. Seasonal germination proportions in the SeedBank pots. The average proportion of germinants in each season is shown for each dormancy*environment treatment. Seasonal germination proportions were calculated from both the Confirmed and Maximum total number of seedlings (see the methods for SeedBank pots germination monitoring in the Supplementary Information for more details). The colors denote the season. The D bars are for Dormant pots and ND bars are for Non-Dormant pots. In Year 2 in the Control treatment, ND seeds had no germination and only 1 Dormant seed germinated.

**Table S5. Test for the effects of experimental environmental treatments on environmental conditions experienced within the cages. A) The effect of environmental treatment, month-year, and the interaction between those effects on daily average, minimum, and maximum soil temperature, and moisture. The effect of environmental treatment (Control versus High Humidity) on PPFD is also shown. For all analysis, a Type III ANOVA was used, with block as a random factor. B) Significant pairwise differences in soil temperature and moisture between environmental treatments within specific Month-Years. The P-values are represented as ***P < 0.001, **P < 0.01, *P < 0.05.**

A) ANOVA Table of Environmental Differences

| | Environmental Treatment | | Month-Year | | Envt*Mon-Year | |
|---|---|---|---|---|---|---|
| | df | F | df | F | df | F |
| Daily Avg Soil Temp | 3 | 4.77** | 30 | 3600.61*** | 88 | 1.78*** |
| Daily Max Soil Temp | 3 | 56.49*** | 30 | 2288.13*** | 88 | 4.89*** |
| Daily Min Soil Temp | 3 | 15.93*** | 30 | 2822.42*** | 88 | 1.76*** |
| Daily Avg SWC | 3 | 423.58*** | 28 | 258.63*** | 84 | 12.44*** |
| Daily Max SWC | 3 | 342.60*** | 28 | 223.39*** | 84 | 10.50*** |
| Daily Min SWC | 3 | 446.54*** | 28 | 236.53*** | 84 | 12.70*** |
| PPFD | 1 | 0.01 | -- | -- | -- | -- |

B) Tukey's Tests of Pairwise Differences within Month-Years

| | Ctrl vs. HH | Ctrl vs. Soil | Ctrl vs. Litter | HH vs. Soil | HH vs. Litter | Soil vs. Litter |
|---|---|---|---|---|---|---|
| Avg Soil Temp | | | | | | |
| May-19 | <0.0001*** | 0.1151 | NA | 1.0000 | NA | NA |
| Max Soil Temp | | | | | | |
| May-19 | <0.0001*** | <0.0001*** | NA | 0.0897 | NA | NA |
| May-21 | 0.8990 | 0.1247 | 1.0000 | 1.0000 | 0.0003*** | <0.0001*** |
| June-21 | 1.0000 | 0.8979 | 1.0000 | 0.1149 | 1.0000 | 0.0010** |
| July-19 | 1.0000 | 0.0471* | 0.0058** | 0.0138* | NA | NA |
| July-21 | 1.0000 | 0.2854 | 1.0000 | 0.0997 | 1.0000 | 0.0024** |
| Aug-19 | 0.0931 | <0.0001*** | 1.0000 | 1.0000 | 0.9998 | 0.1323 |
| Aug-21 | 1.0000 | 0.9443 | 1.0000 | 0.5027 | 1.0000 | 0.0017** |
| Sept-19 | <0.0001*** | <0.0001*** | 0.3692 | 1.0000 | 0.1161 | <0.0001*** |
| Sept-21 | 1.0000 | 0.8208 | 1.0000 | 0.1609 | 1.0000 | 0.0053** |
| Oct-19 | <0.0001*** | 0.0138* | 0.9177 | 1.0000 | 0.7299 | 1.0000 |
| Feb-20 | 0.0712 | 0.9967 | 1.0000 | 1.0000 | 0.0299* | 0.9743 |
| March-20 | 0.0097** | 1.0000 | 1.0000 | 0.9950 | 0.0008*** | 0.9999 |
| Min Soil Temp | | | | | | |
| Nov-19 | 0.4605 | 0.0014** | 0.9565 | 1.0000 | 1.0000 | 1.0000 |

| | Ctrl vs. HH | Ctrl vs. Soil | Ctrl vs. Litter | HH vs. Soil | HH vs. Litter | Soil vs. Litter |
|---|---|---|---|---|---|---|
| Avg SWC | | | | | | |
| May-21 | 0.5580 | 1.0000 | 0.7514 | 0.8148 | <0.0001*** | 0.5484 |
| June-20 | 0.3337 | 0.9803 | 0.9995 | <0.0001*** | <0.0001*** | 1.0000 |
| June-21 | 0.5290 | 0.6998 | 1.0000 | <0.0001*** | 0.0222* | 1.0000 |
| July-20 | <0.0001*** | 1.0000 | 1.0000 | <0.0001*** | <0.0001*** | 1.0000 |
| July-21 | 0.9542 | 0.0002*** | 1.0000 | <0.0001*** | 0.4621 | 0.2020 |
| Aug-20 | 0.7799 | 0.9999 | 1.0000 | 0.0004*** | 0.0028** | 1.0000 |
| Aug-21 | 0.9272 | 1.0000 | 0.9912 | 0.0033** | 0.0008*** | 1.0000 |
| Sept-20 | <0.0001*** | 0.7995 | 1.0000 | <0.0001*** | <0.0001*** | 0.9995 |
| Sept-21 | 0.9943 | 1.0000 | 0.9754 | 0.0983 | 0.0019** | 1.0000 |
| Oct-20 | <0.0001*** | <0.0001*** | 1.0000 | <0.0001*** | <0.0001*** | 0.1124 |
| Oct-21 | 0.2245 | 1.0000 | 1.0000 | 0.0012** | 0.0318* | 1.0000 |
| Nov-20 | <0.0001*** | <0.0001*** | 1.0000 | <0.0001*** | <0.0001*** | <0.0001*** |
| Nov-21 | <0.0001*** | 1.0000 | 1.0000 | 0.0015** | <0.0001*** | 1.0000 |
| Dec-20 | 0.0079** | <0.0001*** | 1.0000 | <0.0001*** | <0.0001*** | <0.0001*** |
| Dec-21 | <0.0001*** | 1.0000 | 1.0000 | 0.3261 | 0.0075** | 1.0000 |
| Jan-20 | 1.0000 | 0.5924 | 1.0000 | 0.0011** | 1.0000 | 0.9223 |
| Jan-21 | <0.0001*** | <0.0001*** | 1.0000 | <0.0001*** | <0.0001*** | <0.0001*** |
| Feb-20 | 1.0000 | 0.0124* | 1.0000 | <0.0001*** | 1.0000 | <0.0001*** |
| Feb-21 | 1.0000 | <0.0001*** | 1.0000 | <0.0001*** | 0.9947 | <0.0001*** |
| March-21 | <0.0001*** | <0.0001*** | 1.0000 | <0.0001*** | <0.0001*** | 0.0127* |
| April-20 | 0.4328 | 0.9672 | 1.0000 | <0.0001*** | 0.9557 | 0.6360 |
| April-21 | 0.5542 | <0.0001*** | 0.0041** | <0.0001*** | <0.0001*** | 1.0000 |
| Max SWC | | | | | | |
| May-21 | 0.9546 | 1.0000 | 0.9652 | 0.9912 | 0.0003*** | 0.9134 |
| June-20 | <0.0001*** | 0.9064 | 1.0000 | <0.0001*** | <0.0001*** | 1.0000 |
| June-21 | 0.9770 | 0.2347 | 1.0000 | <0.0001*** | 0.3725 | 0.9983 |
| July-20 | <0.0001*** | 1.0000 | 1.0000 | <0.0001*** | <0.0001*** | 1.0000 |
| July-21 | 1.0000 | 0.0003*** | 1.0000 | <0.0001*** | 0.9993 | 0.0388* |
| Aug-20 | 0.9955 | 0.9796 | 0.9967 | 0.0008*** | 0.0029** | 1.0000 |
| Aug-21 | 0.9998 | 0.9998 | 1.0000 | 0.0170* | 0.1373 | 1.0000 |
| Sept-20 | 0.3531 | 0.0985 | 1.0000 | <0.0001*** | 0.1127 | 0.7896 |
| Oct-20 | 0.0274* | 0.0039** | 1.0000 | <0.0001*** | <0.0001*** | 0.9544 |
| Oct-21 | 0.7559 | 1.0000 | 1.0000 | 0.0341* | 0.4652 | 1.0000 |
| Nov-20 | 0.0070** | <0.0001*** | 1.0000 | <0.0001*** | <0.0001*** | <0.0001*** |
| Nov-21 | <0.0001*** | 1.0000 | 1.0000 | 0.0346* | 0.0041** | 1.0000 |
| Dec-20 | 1.0000 | <0.0001*** | 1.0000 | <0.0001*** | 0.5431 | <0.0001*** |
| Dec-21 | 0.0083** | 1.0000 | 1.0000 | 0.9359 | 0.2070 | 1.0000 |

| | Ctrl vs. HH | Ctrl vs. Soil | Ctrl vs. Litter | HH vs. Soil | HH vs. Litter | Soil vs. Litter |
|---|---|---|---|---|---|---|
| Max SWC Continued… | | | | | | |
| Jan-20 | 1.0000 | 0.3681 | 1.0000 | 0.0153* | 1.0000 | 0.8140 |
| Jan-21 | 0.2244 | <0.0001*** | 1.0000 | <0.0001*** | 0.1179 | <0.0001*** |
| Feb-20 | 1.0000 | 0.0017** | 1.0000 | <0.0001*** | 1.0000 | <0.0001*** |
| Feb-21 | 1.0000 | <0.0001*** | 1.0000 | <0.0001*** | 1.0000 | <0.0001*** |
| March-21 | 0.0077** | <0.0001*** | 1.0000 | <0.0001*** | 0.0002*** | 0.0024** |
| April-20 | 0.9998 | 0.9843 | 1.0000 | 0.0132* | 1.0000 | 0.9309 |
| April-21 | 0.9578 | <0.0001*** | 0.2182 | <0.0001*** | <0.0001*** | 1.0000 |
| Min SWC | | | | | | |
| May-21 | 0.2314 | 1.0000 | 0.6643 | 0.7166 | <0.0001*** | 0.2330 |
| June-21 | 0.0959 | 0.9989 | 1.0000 | <0.0001*** | 0.0006*** | 1.0000 |
| July-20 | <0.0001*** | 0.9968 | 1.0000 | 0.3055 | 0.0001*** | 1.0000 |
| July-21 | 0.3149 | 0.0132* | 1.0000 | <0.0001*** | 0.0029** | 0.9993 |
| Aug-20 | 0.4177 | 1.0000 | 1.0000 | 0.0002*** | 0.0055** | 1.0000 |
| Aug-21 | 0.5593 | 1.0000 | 0.7820 | 0.0027** | <0.0001*** | 1.0000 |
| Sept-20 | <0.0001*** | 1.0000 | 1.0000 | <0.0001*** | <0.0001*** | 1.0000 |
| Sept-21 | 0.9471 | 1.0000 | 0.7953 | 0.0729 | <0.0001*** | 1.0000 |
| Oct-20 | <0.0001*** | <0.0001*** | 1.0000 | <0.0001*** | <0.0001*** | 0.0228* |
| Oct-21 | 0.1202 | 1.0000 | 1.0000 | 0.0004*** | 0.0026** | 1.0000 |
| Nov-20 | <0.0001*** | <0.0001*** | 1.0000 | <0.0001*** | <0.0001*** | <0.0001*** |
| Nov-21 | <0.0001*** | 1.0000 | 1.0000 | 0.0006*** | <0.0001*** | 1.0000 |
| Dec-20 | <0.0001*** | <0.0001*** | 1.0000 | <0.0001*** | <0.0001*** | <0.0001*** |
| Dec-21 | <0.0001*** | 1.0000 | 1.0000 | 0.1406 | 0.0012** | 1.0000 |
| Jan-20 | 1.0000 | 0.8928 | 1.0000 | 0.0003*** | 0.9997 | 0.9617 |
| Jan-21 | <0.0001*** | <0.0001*** | 1.0000 | <0.0001*** | <0.0001*** | <0.0001*** |
| Feb-20 | 0.7267 | 0.8674 | 1.0000 | <0.0001*** | 1.0000 | 0.0009*** |
| Feb-21 | 0.2070 | <0.0001*** | 1.0000 | <0.0001*** | 0.0948 | <0.0001*** |
| March-21 | <0.0001*** | <0.0001*** | 1.0000 | <0.0001*** | <0.0001*** | 0.1297 |
| April-20 | 0.0296* | 0.9872 | 1.0000 | <0.0001*** | 0.8744 | 0.2900 |
| April-21 | 0.1618 | <0.0001*** | 0.0003*** | <0.0001*** | <0.0001*** | 1.0000 |

### A) Soil Temperature

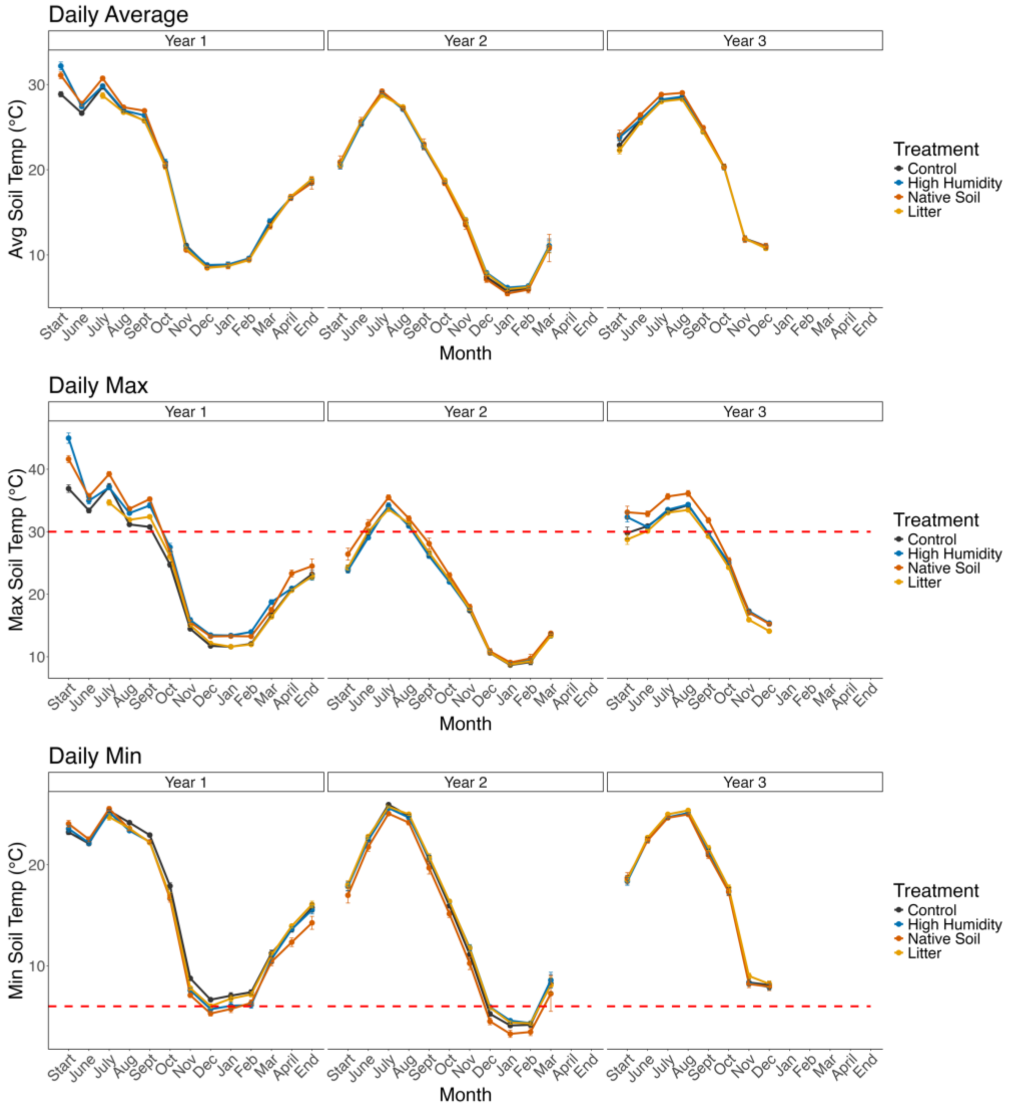

### B) Soil Water Content

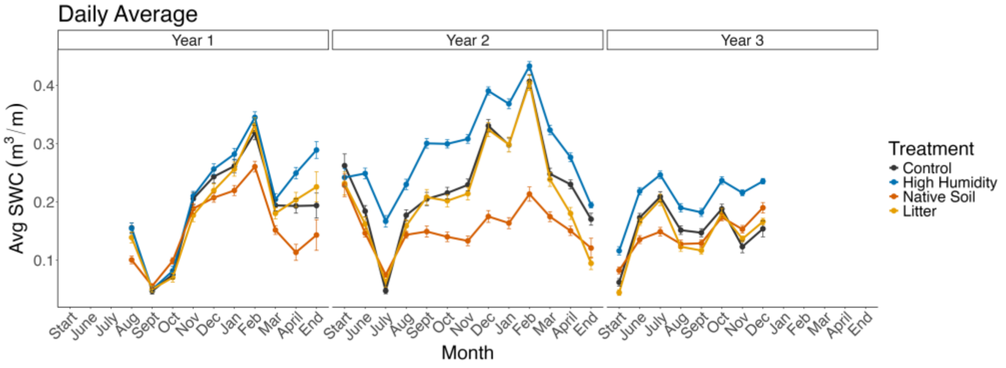

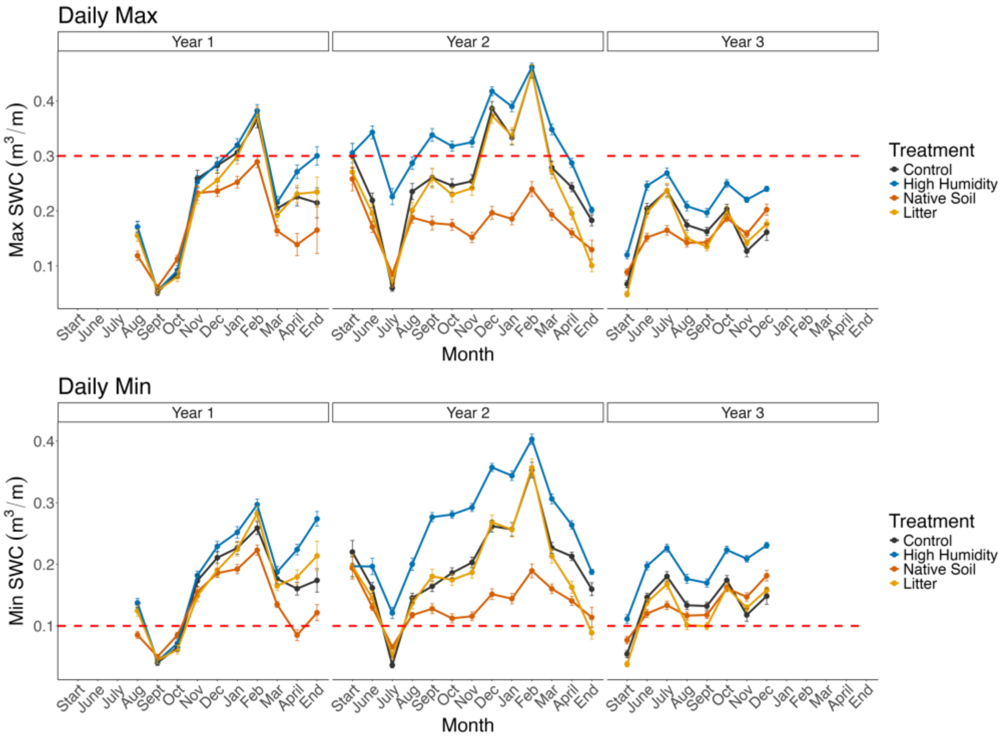

**C) Photosynthetic Photon Flux Density**

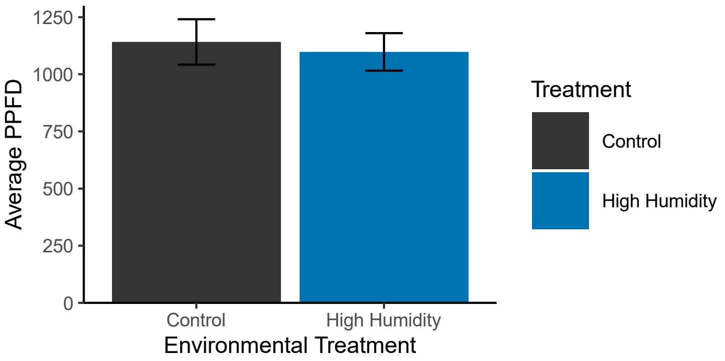

**Figure S7. Effects of the experimental environmental treatments on conditions within cages. Daily average, maximum, and minimum A) Soil temperature (°C) and B) Soil water content ($m^3$ water/$m^3$ soil). Census years are shown such that "Start" represents mid-May (after the dispersal season ends) and "End" represents early-May (the end of the dispersal season). Threshold temperatures (30°C and 6°C) and SWC (0.3 and 0.1) are marked with a red dotted line on the maximum and minimum figures, respectively, to indicate transitions between warm/cool and wet/dry conditions. SWC values 0.0-0.1 represent oven dry to dry soil and 0.3 or higher represents wet to saturated soil. The High Humidity treatment was warmer than the Control treatment (daily average: $P=0.0182$; daily max: $P<0.0001$) and wetter than all the other treatments ($P<0.0001$ for all pairwise tests across average, min, and max). The Native Soil treatment had hotter daily maximum temperatures than the Control treatment ($P<0.0001$) and drier daily average, max, and min than the other treatments ($P<0.0001$ for all). A comparison of C) PPFD between the Control and High Humidity treatment (which had a partial canopy) is also shown but is not significant.**

**Table S6. Effects of dormancy treatment (Dormant and Non-Dormant) and environmental treatments on the proportion of seeds that germinated in the SeedBank Pots. A) The effect of dormancy, environment, the interaction between those effects, and block on the proportion of seeds that germinated in year 1 and in year 2, and seed mortality (proportion of seeds that did not germinate within the 2-year experiment). B) P-values for the pairwise differences between Dormant and Non-dormant treatments within each environmental treatment. "Confirmed total" refers to the number of seedlings that were confirmed to be *A. thaliana* germinants; "Max total" includes the number of seedlings that were suspected of being *A. thaliana,* but which died too soon for definitive identification. P-values are represented as ***P < 0.001, **P < 0.01, *P < 0.05, $^{\Psi}$P < 0.08. Significant main effects that were observed in the main-effect-only logistic regression models are represented as "m".**

**A) Full Models**

| | Confirmed Total | | | | | | Max Total | | | | | |
|---|---|---|---|---|---|---|---|---|---|---|---|---|
| | Year 1 Germination | | Year 2 Germination | | Seed Mortality | | Year 1 Germination | | Year 2 Germination | | Seed Mortality | |
| | df | $X^2$ | df | $X^2$ | df | $X^2$ | df | $X^2$ | df | $X^2$ | df | $X^2$ |
| Dormancy | 1 | 11.65***$^{,m}$ | 1 | 0.01 | 1 | 12.31***$^{,m}$ | 1 | 14.46***$^{,m}$ | 1 | 4.06*$^{,m}$ | 1 | 14.87***$^{,m}$ |
| Envt | 2 | 33.73***$^{,m}$ | 2 | 0.57 | 2 | 33.73***$^{,m}$ | 2 | 34.53***$^{,m}$ | 2 | 0.83 | 2 | 34.91***$^{,m}$ |
| Dorm*Envt | 2 | 5.70$^{\Psi}$ | 2 | 0.07 | 2 | 6.28* | 2 | 3.98 | 2 | 0.31 | 2 | 4.56 |
| Block | 7 | 12.28 | 7 | 5.77 | 7 | 10.95 | 7 | 13.34$^{\Psi}$ | 7 | 9.11 | 7 | 14.40* |

**B) Pairwise Comparisons Between Dormancy Treatments**

| | Confirmed Total | | | Max Total | | |
|---|---|---|---|---|---|---|
| | Year 1 Germination | Year 2 Germination | Seed Mortality | Year 1 Germination | Year 2 Germination | Seed Mortality |
| Control | 0.9416 | 0.5530 | 0.9500 | 0.3368 | 0.2503 | 0.3721 |
| High Humidity | **0.0010**** | 0.1979 | **0.0014**** | **0.0007***** | 0.1041 | **0.0007***** |
| Native Soil | 0.8710 | 0.1784 | 0.8971 | **0.0009***** | 0.1784 | **0.0004***** |

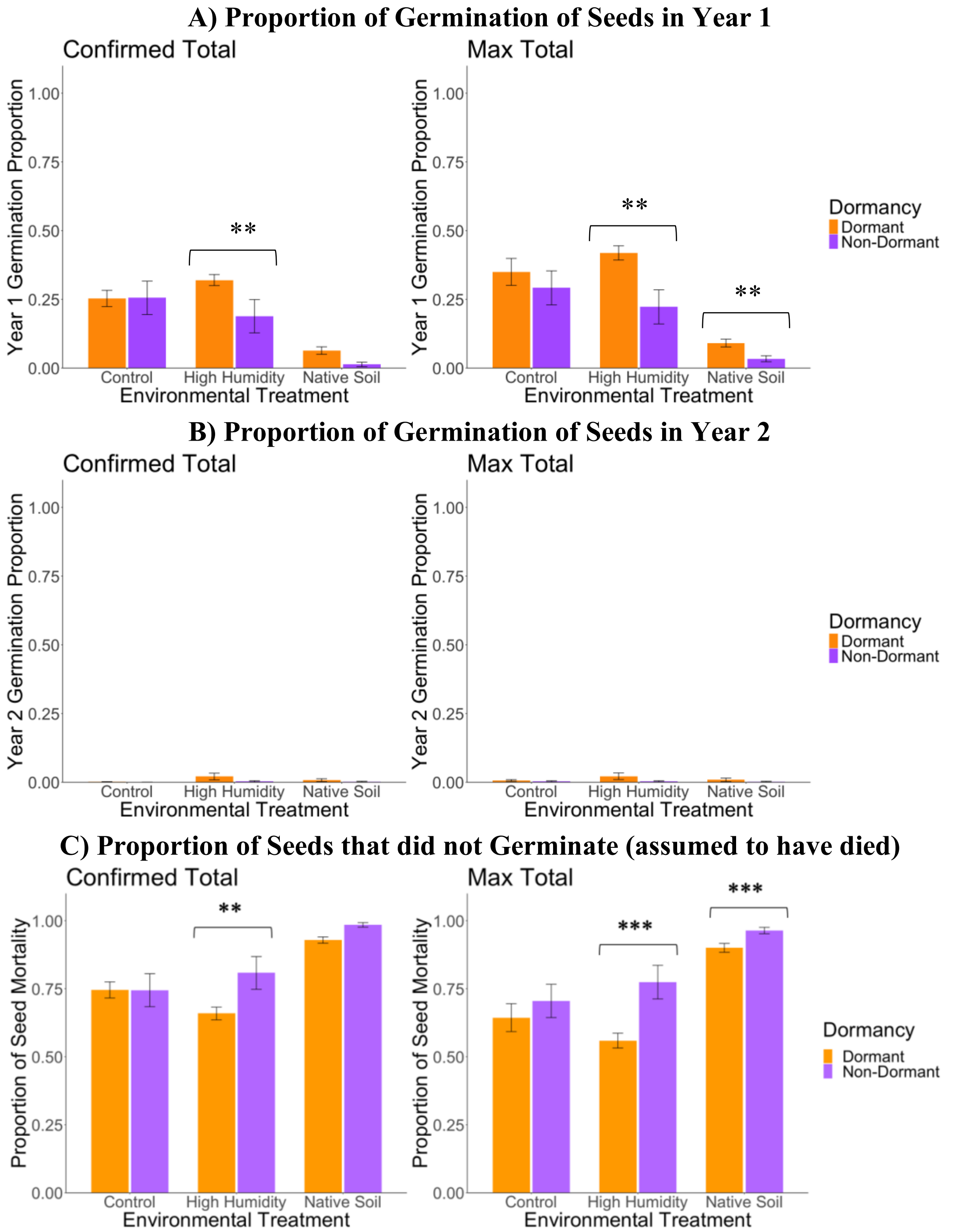

**Figure S8. Proportion of germination in the SeedBank Pots. A) The number of seedlings that germinated in year 1 divided by the 114 total seeds planted in each pot. B) The number of seedlings that germinated in year 2 divided by the 114 total seeds planted in each pot. C) The number of seeds that did not germinate in either year of the experiment divided by the total number of seeds planted in each pot. The data were calculated from the confirmed totals are on the left and the maximum totals are on the right. P-values are represented as ***P < 0.001, **P < 0.01, *P < 0.05.**

A) Seedling Number

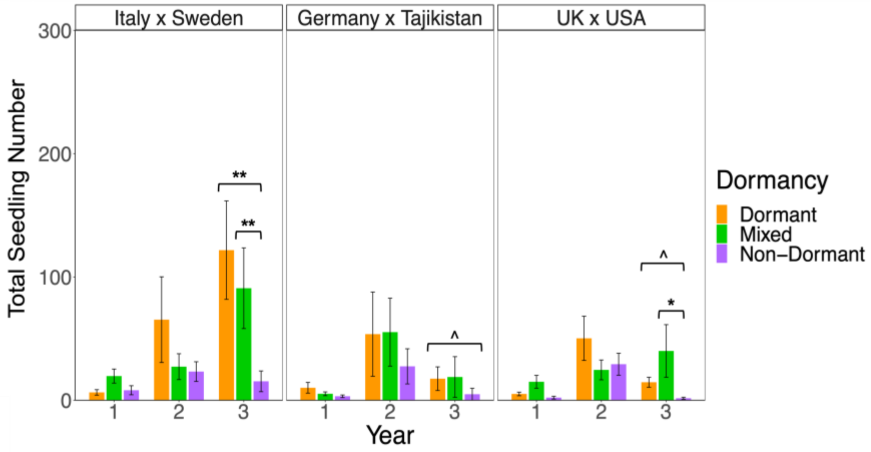

B) Seedling Establishment

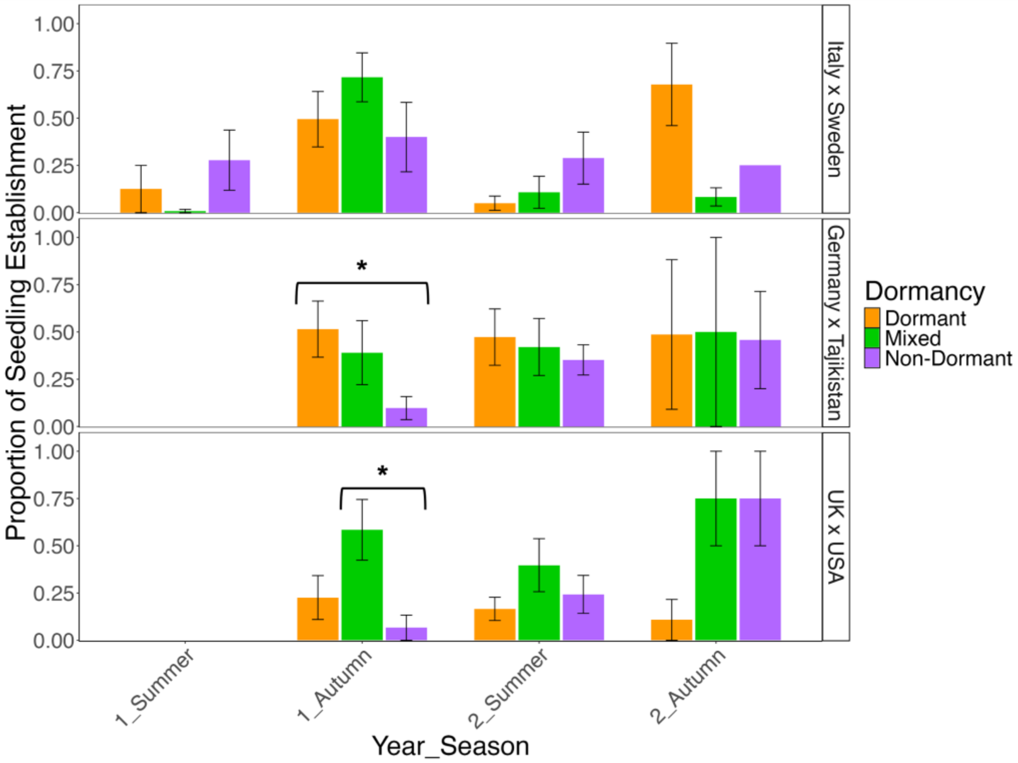

C) Survival to Rosette

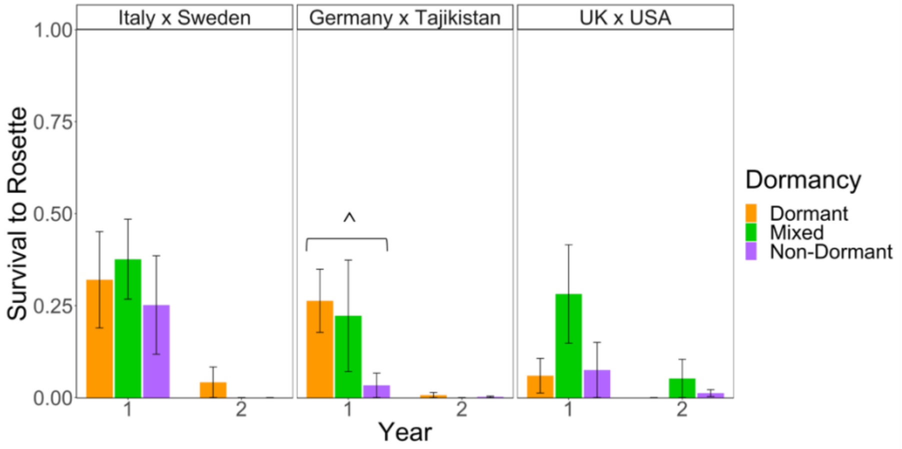

D) Survival to Reproduction

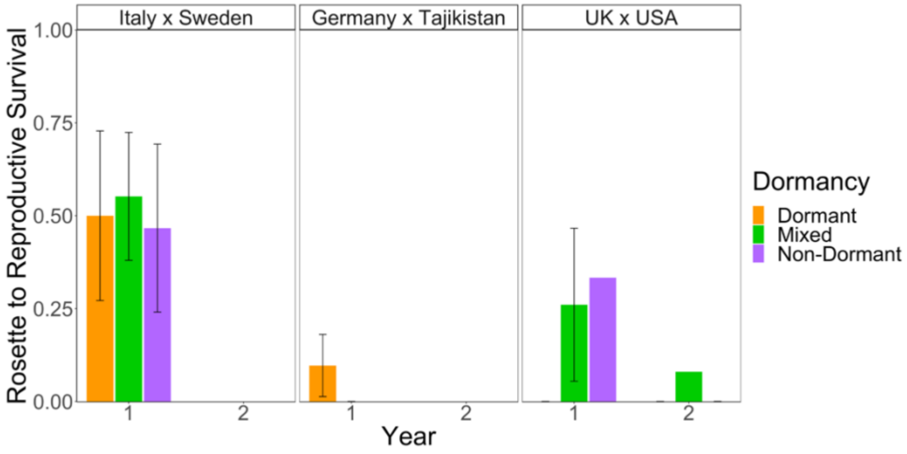

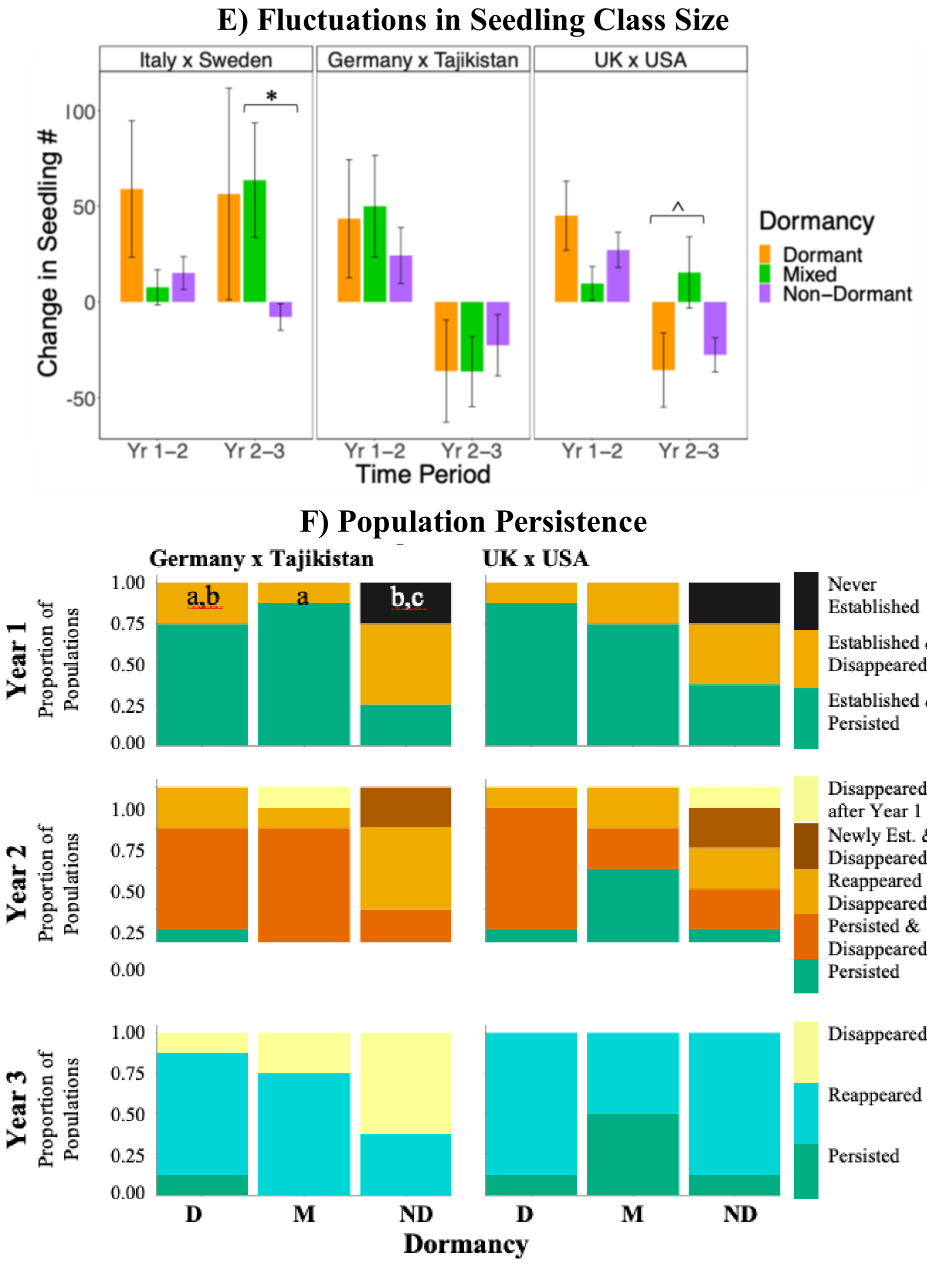

**Figure S9. Comparisons across dormancy treatments for each RIL set in the Control environmental treatment. Dormancy differences in A) the size of the seedling class, B) seedling establishment, C) proportion of survival to rosette, D) proportion of survival to reproduction, E) the change in total seedling number between years, F) for each Germany x Tajikistan and UK x USA in the Control treatment. Dormant populations are in orange, Mixed in green, and Non-Dormant in purple. Raw means with standard errors are shown for A-E. The proportion of populations that persisted (green shading) or went extinct (warm shading) is shown for F (See Fig. 2 for more details). Brackets with asterisks and letters indicate significant differences among dormancy treatments. P-values are represented as ***P < 0.001, **P < 0.01, *P < 0.05, ^P<0.05 prior to correction test.**

**Table S7. Test for differences in demographic performance among dormancy treatments within each RIL set in the Control environmental treatment. See Table 1 for results from the full model. P-values from sub-models used to test for dormancy effects within each RIL set and year. Tukey's test was used to adjust the p-values for each trait analyzed with an ANOVA, while Hochberg tests were used for all traits analyzed with a logistic regression. Note that no adjustment tests were used for survival to reproduction or the chi-square contingency analysis of population persistence with all categories included. "New block" was included in all the sub models for survival to rosette and only in the sub models for Italy x Sweden year 1 for survival to reproduction. Within each RIL set, results for pairwise comparisons between dormancy treatments with all years included ("Dorm vs. Non-Dorm," "Dorm vs. Mixed," and "Non-Dorm vs. Mixed" rows) and models within a single year for each pair-wise combination of dormancy treatments are shown. Note that year 3 data are not available for all survival data. The seedling number fluctuation data were analyzed in two ways (ANOVA and logistic regression); transition 1 represents the change from year 1 to 2 and transition 2 represents the change from year 2 to 3. P-values are represented as ***P < 0.001, **P < 0.01, *P < 0.05, ^P<0.05 prior to correction test.**

| | Seedling Number | Survival to Rosette | Survival to Reproduction | Seedling Class Flucts (ANOVA) | Seedling Class Flucts (LOG REG) | Population Persistence (Binomial) | Population Persistence (CATGs) |
|---|---|---|---|---|---|---|---|
| Italy x Sweden | | | | | | | |
| Dorm vs. Non-Dorm | **0.0137*** | 0.9664 | -- | 0.1197 | 0.3641 | **0.0250*** | -- |
| Year 1 / Trans. 1 | 1.0000 | 0.7986 | 0.9099 | 0.7887 | 1.0000 | 0.3170 | 0.0695$^{\Psi}$ |
| Year 2 / Trans. 2 | 0.9658 | 0.7986 | -- | 0.5349 | 0.8910 | 0.3170 | 0.0966 |
| Year 3 | **0.0067**** | -- | -- | -- | -- | -- | **0.0389*** |
| Dorm vs. Mixed | 0.5421 | 0.9769 | -- | 0.5508 | 1.0000 | 0.6425 | -- |
| Year 1 / Trans. 1 | 0.4883 | 0.7986 | 0.8986 | 0.7520 | 1.0000 | -- | -- |
| Year 2 / Trans. 2 | 0.9926 | 0.7986 | -- | 0.9990 | 1.0000 | 0.6963 | 0.5896 |
| Year 3 | 1.0000 | -- | -- | -- | -- | -- | 0.5896 |
| Non-Dorm vs. Mixed | **0.0012**** | 0.9975 | -- | 0.0665 | 0.3641 | **0.0095**** | -- |
| Year 1 / Trans. 1 | 0.5992 | 0.7986 | 0.4386 | 0.9882 | 1.0000 | 0.3170 | 0.0695$^{\Psi}$ |
| Year 2 / Trans. 2 | 0.9992 | -- | -- | **0.0282*** | 1.0000 | 0.3170 | 0.0728$^{\Psi}$ |
| Year 3 | **0.0026**** | -- | -- | -- | -- | -- | **0.0117*** |
| Germany x Tajikistan | | | | | | | |
| Dorm vs. Non-Dorm | **0.0140*** | 0.1121 | -- | 0.9022 | 0.7314 | **0.0105*** | -- |
| Year 1 / Trans. 1 | 0.8320 | 0.2460^ | 0.9695 | 0.9344 | 1.0000 | 0.6405 | 0.0970 |

| | Seedling Number | Survival to Rosette | Survival to Reproduction | Seedling Class Flucts (ANOVA) | Seedling Class Flucts (LOG REG) | Population Persistence (Binomial) | Population Persistence (CATGs) |
|---|---|---|---|---|---|---|---|
| Year 2 / Trans. 2 | 0.9733 | 0.6246 | -- | 0.9757 | 1.0000 | 0.6405 | 0.1753 |
| Year 3 | 0.1445^ | -- | -- | -- | -- | 0.6405 | 0.0970 |
| Dorm vs. Mixed | 0.2908 | 0.9652 | -- | 0.9055 | 0.2999 | 0.6802 | -- |
| Year 1 / Trans. 1 | 0.9976 | 0.6246 | 0.9590 | 0.9980 | 1.0000 | 0.6405 | 0.5218 |
| Year 2 / Trans. 2 | 0.9985 | 0.6246 | -- | 1.0000 | 1.0000 | 0.6405 | 0.4891 |
| Year 3 | 0.9117 | -- | -- | -- | -- | 0.6405 | 0.5134 |
| Non-Dorm vs. Mixed | 0.2015 | 0.9773 | -- | 0.7601 | 0.5530 | **0.0272*** | -- |
| Year 1 / Trans. 1 | 0.9853 | 0.6246 | -- | 0.7846 | 1.0000 | 0.6405 | **0.0373*** |
| Year 2 / Trans. 2 | 0.9997 | 0.6246 | -- | 0.9579 | 1.0000 | -- | 0.0786$^{\Psi}$ |
| Year 3 | 0.7792 | -- | -- | -- | -- | 0.6405 | 0.1306 |
| UK x USA | | | | | | | |
| Dorm vs. Non-Dorm | **0.0070**** | 0.9718 | -- | 0.7371 | 1.0000 | 0.3491 | -- |
| Year 1 / Trans. 1 | 0.7691 | 1.0000 | 0.3676 | 0.8221 | 0.6405 | 0.8154 | 0.1003 |
| Year 2 / Trans. 2 | 0.9564 | 1.0000 | -- | 0.9801 | 0.6405 | 1.0000 | 0.2548 |
| Year 3 | 0.0822^ | -- | -- | -- | -- | -- | 1.0000 |
| Dorm vs. Mixed | 0.6515 | 0.9670 | -- | 0.6489 | 0.5921 | 0.3687 | -- |
| Year 1 / Trans. 1 | 0.9324 | 1.0000 | 0.9495 | 0.4574 | 0.6405 | 1.0000 | 0.5218 |
| Year 2 / Trans. 2 | 0.9416 | 1.0000 | -- | 0.1704^ | 0.6405 | 0.8154 | 0.1266 |
| Year 3 | 0.9731 | -- | -- | -- | -- | -- | 0.1056 |
| Non-Dorm vs. Mixed | **0.0033*** | 0.4180 | -- | 0.3051 | 0.6178 | 0.0715$^{\Psi}$ | -- |
| Year 1 / Trans. 1 | 0.2441^ | 1.0000 | 0.6573 | 0.7396 | 0.6405 | 0.8154 | 0.2019 |
| Year 2 / Trans. 2 | 1.0000 | 1.0000 | -- | 0.0879^ | 0.6405 | 0.8154 | 0.3084 |
| Year 3 | **0.0192*** | -- | -- | -- | -- | -- | 0.1056 |

**Table S8. Test for differences among dormancy treatments within each RIL set for seasonal seedling establishment. Within each RIL set, p-values for pairwise comparisons between dormancy treatments with both years included ("Dorm vs. Non-Dorm," "Dorm vs. Mixed," and "Non-Dorm vs. Mixed" rows) and models within a single year for each pair-wise combination of dormancy treatments are shown. Note that year 3 data are not available. Block as the two halves of the field was included in the sub-models when possible; [B] represents comparisons where block was included. Significant p-values are bolded.**

| | Summer | Autumn |
|---|---|---|
| Italy x Sweden | | |
| Dorm vs. Non-Dorm | 0.1365 | 0.1683 |
| Year 1 | 0.5191 | 0.4750[B] |
| Year 2 | 0.1532[B] | 0.8922 |
| Dorm vs. Mixed | 0.6243 | 0.7409[B] |
| Year 1 | 0.6212 | 0.2763[B] |
| Year 2 | 0.9206[B] | 0.9972[B] |
| Non-Dorm vs. Mixed | 0.0786[B] | 0.8084 |
| Year 1 | 0.1583[B] | 0.1905[B] |
| Year 2 | 0.3930[B] | 0.8922 |
| Germany x Tajikistan | | |
| Dorm vs. Non-Dorm | -- | 0.1052 |
| Year 1 | -- | **0.0397[B]** |
| Year 2 | 0.8354[B] | 1.0000 |
| Dorm vs. Mixed | -- | 0.5329 |
| Year 1 | -- | 0.0905[B] |
| Year 2 | 0.1875[B] | 1.0000 |
| Non-Dorm vs. Mixed | -- | 0.3924 |
| Year 1 | -- | 0.1894 |
| Year 2 | 0.9545[B] | 1.0000 |
| UK x USA | | |
| Dorm vs. Non-Dorm | -- | 0.9505 |
| Year 1 | -- | 0.2288 |
| Year 2 | 0.2537[B] | 0.9980 |
| Dorm vs. Mixed | -- | **0.0257** |
| Year 1 | -- | 0.1169 |
| Year 2 | 0.2387 | 0.9980 |

| | Summer | Autumn |
|---|---|---|
| Non-Dorm vs. Mixed | -- | 0.0753 |
| Year 1 | -- | **0.0369** |
| Year 2 | 0.8196[B] | 1.0000 |

### A) Individual Survival to Reproduction

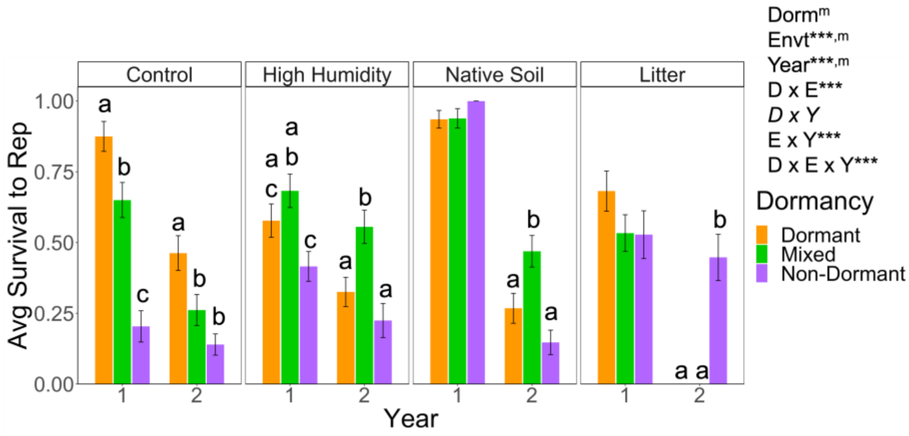

### B) Population-level Survival to Reproduction

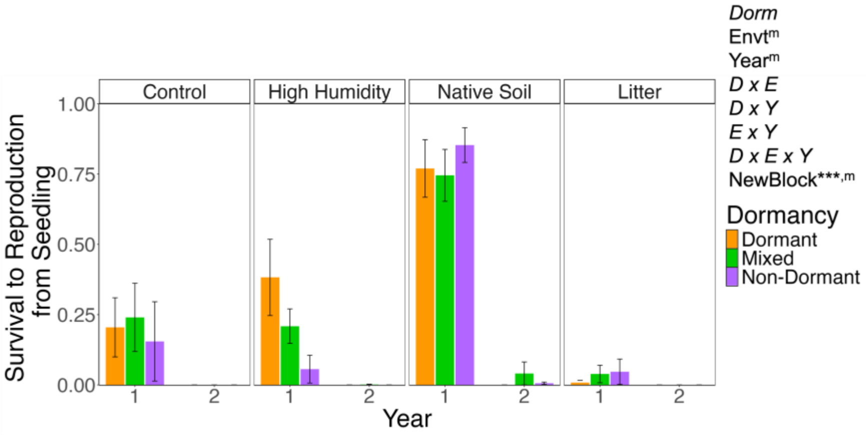

**Figure S10. A) Individual survival to reproduction versus B) population-level survival from seedling to reproduction. The results from the full models are indicated on the right. P-values are represented as ***P < 0.001, **P < 0.01, *P < 0.05, $^{\Psi}$P <0.08. Significant main effects that were observed in the main-effect-only logistic regression models are represented as $^{m}$. Italics indicate non-significant effects. Note that quasi-complete separation of data was detected for the full population-level model. Significant pairwise differences between dormancy treatments, within an environmental treatment, are indicated by different letters. The figure in A) is the same as in Figure 5 in the main text.**

**Table S9. Test for demographic differences among dormancy treatments within each environmental treatment. See Table 2 in the main text for results of the full model. P-values are from sub-models used to test for dormancy effects within each environmental treatment. Tukey's test was used to adjust the p-values for each trait analyzed with an ANOVA, while Hochberg tests were used for all traits analyzed with a logistic regression or Friedman's test. "New block" was included in all the sub models for survival to rosette and only in the sub models for year 1 for all treatments and year 2 for native soil comparisons without the non-dormant treatment for survival to reproduction. Note that year 3 data are not available for survival to reproduction. The seedling class fluctuation data were analyzed in two ways (ANOVA and logistic regression); transition 1 represents the change from year 1 to 2 and transition 2 represents the change from year 2 to 3. P-values are represented as ***P < 0.001, **P < 0.01, *P < 0.05, ^P<0.05 prior to correction test, $^{\Psi}$P <0.08 prior to correction test.**

| | Seedling Number | Survival to Rosette | Survival to Rep. | Seedling Class Flucts (ANOVA) | Seedling Class Flucts (LOG REG) | Population Persistence (Binomial) | Population Persistence (CATGs) |
|---|---|---|---|---|---|---|---|
| Control | | | | | | | |
| Dorm vs. Non-Dorm | **0.0137*** | 0.9664 | -- | 0.1197 | 0.3641 | **0.0250*** | -- |
| Year 1 / Transition 1 | 1.0000 | 0.7986 | 0.9099 | 0.7887 | 1.0000 | 0.3170 | 0.0695$^{\Psi}$ |
| Year 2 / Transition 2 | 0.9658 | 0.7986 | -- | 0.5349 | 0.8910 | 0.3170 | 0.0966 |
| Year 3 | **0.0067**** | -- | -- | -- | -- | -- | **0.0389*** |
| Dorm vs. Mixed | 0.5421 | 0.9769 | -- | 0.5508 | 1.0000 | 0.6425 | -- |
| Year 1 / Transition 1 | 0.4883 | 0.7986 | 0.8986 | 0.7520 | 1.0000 | -- | -- |
| Year 2 / Transition 2 | 0.9926 | 0.7986 | -- | 0.9990 | 1.0000 | 0.6963 | 0.5896 |
| Year 3 | 1.0000 | -- | -- | -- | -- | -- | 0.5896 |
| Non-Dorm vs. Mixed | **0.0012**** | 0.9975 | -- | 0.0665 | 0.3641 | **0.0095**** | -- |
| Year 1 / Transition 1 | 0.5992 | 0.7986 | 0.4386 | 0.9882 | 1.0000 | 0.3170 | 0.0695$^{\Psi}$ |
| Year 2 / Transition 2 | 0.9992 | -- | -- | **0.0282*** | 1.0000 | 0.3170 | 0.0728$^{\Psi}$ |
| Year 3 | **0.0026**** | -- | -- | -- | -- | -- | **0.0117*** |
| High Humidity | | | | | | | |
| Dorm vs. Non-Dorm | **0.0052**** | **0.0423*** | 0.9965 | 0.0555 | 0.0711 | **0.0133*** | -- |
| Year 1 / Transition 1 | 0.9985 | 0.1362^ | 0.5160 | 0.7180 | 0.6454 | 0.4410 | **0.0455*** |
| Year 2 / Transition 2 | 0.9483 | 0.8364 | -- | 0.2968 | 0.3804$^{\Psi}$ | 0.4410 | 0.2060 |
| Year 3 | **0.0056**** | -- | -- | -- | -- | -- | 0.1306 |

| | Seedling Number | Survival to Rosette | Survival to Rep. | Seedling Class Flucts (ANOVA) | Seedling Class Flucts (LOG REG) | Population Persistence (Binomial) | Population Persistence (CATGs) |
|---|---|---|---|---|---|---|---|
| Dorm vs. Mixed | 0.8354 | 0.8091 | 0.9722 | 0.5797 | 0.6900 | 1.0000 | -- |
| Year 1 / Transition 1 | 0.9918 | 0.8364 | 0.9495 | 0.7423 | 0.6454 | 1.0000 | 1.0000 |
| Year 2 / Transition 2 | 0.9838 | 0.8364 | 0.9495 | 0.2961 | 0.6454 | 1.0000 | 1.0000 |
| Year 3 | 0.9466 | -- | -- | -- | -- | -- | 1.0000 |
| Non-Dorm vs. Mixed | **0.0057**** | **0.0325*** | 0.9726 | 0.1728 | 0.1546 | **0.0133*** | -- |
| Year 1 / Transition 1 | 0.9488 | $0.3595^{\Psi}$ | 0.6344 | $0.1950^{\Psi}$ | 0.6454 | 0.4410 | **0.0455*** |
| Year 2 / Transition 2 | 0.7235 | 0.8364 | 0.9495 | 0.9998 | 0.6454 | 0.4410 | 0.2060 |
| Year 3 | 0.0708^ | -- | -- | -- | -- | -- | 0.1306 |
| Native Soil | | | | | | | |
| Dorm vs. Non-Dorm | 0.1748 | **0.0460*** | 1.0000 | 0.1064 | **0.0481*** | 0.6655 | -- |
| Year 1 / Transition 1 | 0.9990 | 0.6963 | 0.9786 | 0.9750 | 1.0000 | -- | -- |
| Year 2 / Transition 2 | 0.9975 | 0.4888 | -- | $0.2309^{\Psi}$ | 1.0000 | 1.0000 | 0.5488 |
| Year 3 | **0.0306*** | -- | -- | -- | -- | 1.0000 | 0.5738 |
| Dorm vs. Mixed | 0.7625 | 0.2328 | 0.9678 | 0.7623 | 0.2262 | 0.6425 | -- |
| Year 1 / Transition 1 | 0.9722 | 0.4888 | 0.9786 | 0.8961 | 1.0000 | -- | -- |
| Year 2 / Transition 2 | 0.9735 | 0.6963 | 0.9786 | 0.6740 | 1.0000 | 1.0000 | 0.5796 |
| Year 3 | 0.9190 | -- | -- | -- | -- | -- | 0.5896 |
| Non-Dorm vs. Mixed | 0.1455 | **0.0500*** | 0.9773 | 0.3468 | 0.3170 | 0.3940 | -- |
| Year 1 / Transition 1 | 0.9983 | 0.6963 | 0.9786 | 0.7340 | 1.0000 | -- | -- |
| Year 2 / Transition 2 | 0.9993 | $0.3384^{\Psi}$ | 0.9786 | 0.9871 | 1.0000 | 1.0000 | 0.5896 |
| Year 3 | 0.4654 | -- | -- | -- | -- | 1.0000 | 0.4493 |
| Litter | | | | | | | |
| Dorm vs. Non-Dorm | 0.4503 | 0.9785 | -- | 0.5924 | 0.6234 | 0.6188 | -- |
| Year 1 / Transition 1 | 0.9992 | 1.0000 | 0.4548 | 0.7679 | 1.0000 | 1.0000 | 0.5738 |
| Year 2 / Transition 2 | 0.9922 | 1.0000 | -- | 0.9970 | 1.0000 | 1.0000 | 0.5249 |
| Year 3 | 0.8748 | -- | -- | -- | -- | -- | 0.3017 |
| Dorm vs. Mixed | 0.4333 | 0.9675 | -- | 0.3812 | 0.8012 | 0.4745 | -- |

| | Seedling Number | Survival to Rosette | Survival to Rep. | Seedling Class Flucts (ANOVA) | Seedling Class Flucts (LOG REG) | Population Persistence (Binomial) | Population Persistence (CATGs) |
|---|---|---|---|---|---|---|---|
| Year 1 / Transition 1 | 0.8539 | 1.0000 | 0.6059 | 1.0000 | 1.0000 | 1.0000 | 0.3515 |
| Year 2 / Transition 2 | 0.9990 | 1.0000 | -- | 0.5826 | 1.0000 | -- | 0.5497 |
| Year 3 | 0.9912 | -- | -- | -- | -- | -- | -- |
| Non-Dorm vs. Mixed | 0.1842 | 0.8028 | -- | 0.2863 | 0.8175 | 0.6092 | -- |
| Year 1 / Transition 1 | 0.9650 | 1.0000 | 0.4923 | 0.8895 | 1.0000 | 1.0000 | 0.4966 |
| Year 2 / Transition 2 | 1.0000 | 1.0000 | -- | 0.8392 | 1.0000 | 1.0000 | 0.5698 |
| Year 3 | 0.7056 | -- | -- | -- | -- | -- | 0.3017 |

**Table S10. Test for effects of environmental treatment within each dormancy treatment, for the Italy x Sweden RIL set. See Table 2 for results of the full model. P-values from sub-models were used to test for the effects of environmental treatment within each dormancy treatment. "Year" represents "Transition" (ex: year 1 to 2) for seedling class fluctuations and "Year-Season" for seedling establishment; 1-Summer was removed from the analysis; block was excluded as well. Note that year 3 data are not available for the survival data and per-capita reproductive output. Only year 1 was analyzed for survival to reproduction, and per-capita reproductive output. The fluctuations in seedling number data were analyzed in two ways (ANOVA and logistic regression). The P-values for the interaction between dormancy and the environment are shown in the last row; these are from the full model results shown in Table 2. P-values are represented as ***P < 0.001, **P < 0.01, *P < 0.05, $^{\Psi}$P <0.08 and significant environmental effects are bolded. Significant main effects that were observed in the main-effect-only logistic regression models are represented as $^{m}$.**

| | Seedling Number | Seedling Est. | Survival to Rosette | Survival to Rep. | Per-Capita Rep. Output | Seedling Class Flucts (ANOVA) | Seedling Class Flucts (LOG REG) | Population Persistence |
|---|---|---|---|---|---|---|---|---|
| Dormant | | | | | | | | |
| Envt | **0.0027**** | 0.2162 | **0.0036**,$^{m}$** | 0.0630$^{\Psi}$ | **0.0007***** | 0.1188 | 0.1882 | 0.2015$^{m}$ |
| Year | 0.0001*** | 0.0005***,$^{m}$ | 0.9532$^{m}$ | -- | -- | 0.6041 | 0.2174 | 0.0004***,$^{m}$ |
| Envt*Year | 0.5728 | **0.0344*** | 0.8055 | -- | -- | 0.8119 | 0.3284 | 0.7062 |
| Mixed | | | | | | | | |
| Envt | **0.0243*** | **0.0436*,$^{m}$** | **0.0114*,$^{m}$** | **0.0142*** | **<0.0001***** | 0.5792 | 0.4249 | 0.2985$^{m}$ |
| Year | 0.1442 | 0.0012**,$^{m}$ | 0.9480$^{m}$ | -- | -- | 0.2731 | 0.9471 | 0.0008***,$^{m}$ |
| Envt*Year | 0.3559 | 0.1198 | 0.7132 | -- | -- | **0.0093**** | 0.7373 | 0.5536 |
| Non-Dormant | | | | | | | | |
| Envt | 0.2688 | 0.9259 | 0.2144$^{m}$ | 0.0975 | 0.4125 | 0.8620 | 0.2318 | 0.2887$^{m}$ |
| Year | 0.1396 | 0.0003***,$^{m}$ | 0.9439$^{m}$ | -- | -- | 0.8074 | 0.0732 | <0.0001***,$^{m}$ |
| Envt*Year | 0.4467 | **0.0348*** | **0.0285*** | -- | -- | 0.5207 | 0.6264 | 0.2294 |
| Dorm*Envt (Table 2) | 0.7236 | 0.4235 | 0.9180 | 0.5933 | 0.6803 | 0.5165 | 0.5968 | 0.7575 |

**Table S11. Sensitivities and asymptotic lambdas based on average vital rates matrices for each of four scenarios/models of environmental quality and seed-bank presence. The average vital rates matrices were calculated from the average of the population vital rates for each dormancy*environment combination. Matrix elements are as follows: top left, the contribution of fresh seeds at time t to fresh seeds at time t+2; top right, the contribution of fresh seeds at time t to seedbank seeds at time t+2; bottom left, the contribution of seedbank seeds to fresh seeds at time t+2; bottom right, the contribution of seedbank seeds at time t to seedbank seeds at time t+2 (see Fig. S3 for more details).**

**Model 1: No between year dormancy, two consecutive favorable years for survival and reproduction**

| | Control | | High Humidity | | Native Soil | | Litter | |
|---|---|---|---|---|---|---|---|---|
| Dormant | 1 | 0.069 | 1 | 0.020 | 1 | 0.002 | 0 | 0 |
| | 0 | 0 | 0 | 0 | 0 | 0 | 0 | 1 |
| | $\lambda = 26.03$ | | $\lambda = 437.28$ | | $\lambda = 3674.97$ | | $\lambda = 0$ | |
| Mixed | 1 | 0.011 | 1 | 0.014 | 1 | 0.002 | 1 | 0.61 |
| | 0 | 0 | 0 | 0 | 0 | 0 | 0 | 0 |
| | $\lambda = 781.46$ | | $\lambda = 442.48$ | | $\lambda = 892.45$ | | $\lambda = 0.15$ | |
| Non-Dormant | 1 | 0.136 | 1 | 0.176 | 1 | 0.000 | 1 | 0.689 |
| | 0 | 0 | 0 | 0 | 0 | 0 | 0 | 0 |
| | $\lambda = 4.13$ | | $\lambda = 1.55$ | | $\lambda = 20243.26$ | | $\lambda = 0.07$ | |

**Model 2: Maximum between year dormancy, two consecutive favorable years for survival and reproduction**

| | Control | | High Humidity | | Native Soil | | Litter | |
|---|---|---|---|---|---|---|---|---|
| Dormant | 0.732 | 0.032 | 0.963 | 0.018 | 0.996 | 0.002 | 0 | 0 |
| | 6.104 | 0.268 | 1.928 | 0.037 | 2.269 | 0.004 | 0 | 1 |
| | $\lambda = 64.72$ | | $\lambda = 472.69$ | | $\lambda = 3701.08$ | | $\lambda = 0$ | |
| Mixed | 0.979 | 0.010 | 0.980 | 0.014 | 0.997 | 0.002 | 0.656 | 0.189 |
| | 1.975 | 0.021 | 1.389 | 0.020 | 1.607 | 0.003 | 1.191 | 0.344 |
| | $\lambda = 816.38$ | | $\lambda = 460.83$ | | $\lambda = 898.51$ | | $\lambda = 0.660$ | |

| | | | | | | | | |
|---|---|---|---|---|---|---|---|---|
| Non-Dormant | 0.853 | 0.096 | 0.748 | 0.087 | 1.000 | 0.000 | 0.589 | 0.123 |
| | 1.312 | 0.147 | 2.158 | 0.252 | 1.961 | 0.000 | 1.970 | 0.411 |
| | $\lambda = 6.03$ | | $\lambda = 3.53$ | | $\lambda = 20262.19$ | | $\lambda = 0.775$ | |

**Model 3: No between year dormancy, 1 favorable year followed by an unfavorable year.**

| | Control | | High Humidity | | Native Soil | | Litter | |
|---|---|---|---|---|---|---|---|---|
| Dormant | NA | NA | NA | NA | NA | NA | 0 | 0 |
| | NA | NA | NA | NA | NA | NA | 0 | 1 |
| | $\lambda = 0$ | | $\lambda = 0$ | | $\lambda = 0$ | | $\lambda = 0$ | |
| Mixed | NA | NA | 1 | 90.288 | NA | NA | NA | NA |
| | NA | NA | 0 | 0 | NA | NA | NA | NA |
| | $\lambda = 0$ | | $\lambda = 0.07$ | | $\lambda = 0$ | | $\lambda = 0$ | |
| Non-Dormant | NA | NA | NA | NA | NA | NA | NA | NA |
| | NA | NA | NA | NA | NA | NA | NA | NA |
| | $\lambda = 0$ | | $\lambda = 0$ | | $\lambda = 0$ | | $\lambda = 0$ | |

**Model 4: Maximum between year dormancy, 1 favorable year followed by an unfavorable year.**

| | Control | | High Humidity | | Native Soil | | Litter | |
|---|---|---|---|---|---|---|---|---|
| Dormant | 0 | 0.076 | 0 | 0.480 | 0 | 0.438 | 0 | 0 |
| | 0 | 1 | 0 | 1 | 0 | 1 | 0 | 1 |
| | $\lambda = 23.68$ | | $\lambda = 18.05$ | | $\lambda = 13.08$ | | $\lambda = 0$ | |
| Mixed | 0 | 0.485 | 0.008 | 0.687 | 0 | 0.618 | 0 | 0.263 |
| | 0 | 1 | 0.011 | 0.992 | 0 | 1 | 0 | 1 |
| | $\lambda = 17.65$ | | $\lambda = 9.34$ | | $\lambda = 3.04$ | | $\lambda = 0.35$ | |
| Non-Dormant | 0 | 0.538 | 0 | 0.230 | 0 | 0.510 | 0 | 0.091 |
| | 0 | 1 | 0 | 1 | 0 | 1 | 0 | 1 |
| | $\lambda = 1.04$ | | $\lambda = 1.19$ | | $\lambda = 9.47$ | | $\lambda = 0.54$ | |
